# Energy Aware Technology Mapping of Genetic Logic Circuits

**DOI:** 10.1101/2024.06.27.601038

**Authors:** Erik Kubaczka, Maximilian Gehri, Jérémie J. M. Marlhens, Tobias Schwarz, Maik Molderings, Nicolai Engelmann, Hernan G. Garcia, Christian Hochberger, Heinz Koeppl

## Abstract

Energy and its dissipation are fundamental to all living systems, including cells. Insufficient abundance of energy carriers -as caused by the additional burden of artificial genetic circuits-shifts a cell’s priority to survival, also impairing the functionality of the genetic circuit. Moreover, recent works have shown the importance of energy expenditure in information transmission. Despite living organisms being non-equilibrium systems, non-equilibrium models capable of accounting for energy dissipation and non-equilibrium response curves are not yet employed in genetic design automation (GDA) software. To this end, we introduce Energy Aware Technology Mapping, the automated design of genetic logic circuits with respect to energy efficiency and functionality. The basis for this is an energy aware non-equilibrium steady state (NESS) model of gene expression, capturing characteristics like energy dissipation -which we link to the entropy production rate- and transcriptional bursting, relevant to eukaryotes as well as prokaryotes. Our evaluation shows that a genetic logic circuit’s functional performance and energy efficiency are disjoint optimization goals. For our benchmark, energy efficiency improves by 37.2% on average when comparing to functionally optimized variants. We discover a linear increase in energy expenditure and overall protein expression with the circuit size, where Energy Aware Technology Mapping allows for designing genetic logic circuits with the energy efficiency of circuits that are one to two gates smaller. Structural variants improve this further, while results show the Pareto dominance among structures of a single Boolean function. By incorporating energy demand into the design, Energy Aware Technology Mapping enables energy efficiency by design. This extends current GDA tools and complements approaches coping with burden *in vivo*.

**TOC Graphic:** 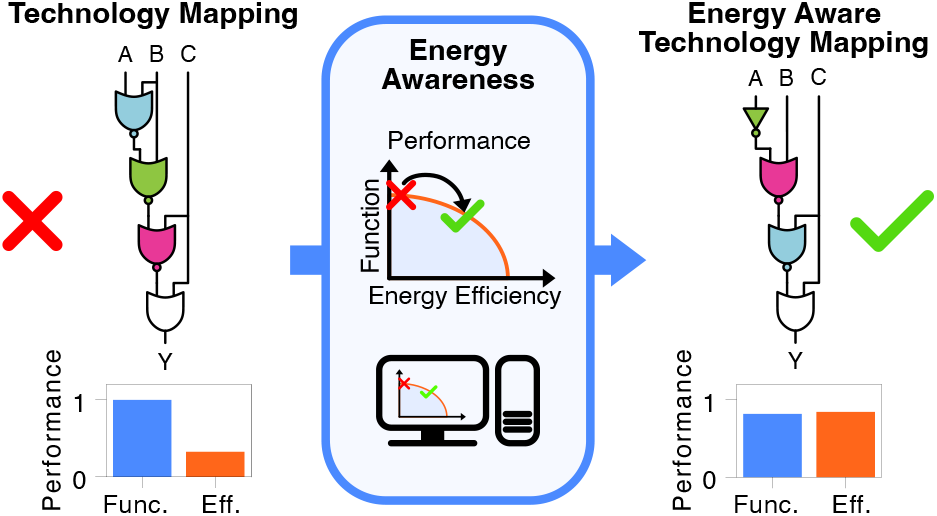

## 1 Introduction

Life is non-equilibrium (*1*), and so energy and its dissipation is essential for life and the function of biological organisms and systems (*2, 3*). While the absence of energy is incompatible with life (*1, 2*), a decrease in its availability has detrimental effects on a cell’s metabolism, and consequently, on its growth, fitness, and gene expression (*4–10*). Such a decrease can be sourced in insufficient nutrition (*4*) of cells, but also in the burden imposed by the insertion of engineered genetic circuits (*8, 10*) and the expression of heterologous proteins (*5*). Metabolic burden, also observed in *S. cerevisiae* and *E. coli*, refers to the diversion of resources from the host to synthetic constructs, affecting the availability of energy, nutrients, ribosomes, and RNA polymerase, as well as reducing cellular fitness (*5, 6*). This allocation away from maintenance and growth compromises the host organism’s physiological functions. As the functionality of synthetic constructs such as engineered genetic circuits depends on sufficient dynamics of proteins and other molecules (*11*), reliable and well-functioning host organisms are essential (*7, 10*). Consequently, a trade-off between function and energy efficiency emerges, affecting reaction levels even at the promoter scale (*12–14*).

The described importance of energy did not hinder the wide application of equilibrium gene expression models (*15–18*). Developed in the context of bacterial transcription, these models assume that regulatory mechanisms, such as transcription factor binding to DNA, operate at thermodynamic equilibrium (*15–17, 19*). While this is reasonable from a modeling perspective for prokaryotes, non-equilibrium processes and the inherent energy dissipation are essential for gene regulation (*1, 3, 19–23*). In particular, the sharpness and sensitivity of the eukaryotic gene response surpasses the so-called Hopfield barriers (*3, 12, 21*), which set an upper limit in the equilibrium case and were first described by J. Hopfield in the context of kinetic proofreading (*24*). Besides this, transcriptional bursting is characteristic of eukaryotic gene expression (*12, 25–28*), but also occurs in prokaryotes (*23, 29*). These bursts are characterized by durations of transcriptional activity significantly longer than the binding of single transcription factors, which lasts a few seconds (*21, 26, 28*). Mechanisms for this may include multistep activation, where transcription factor abundance modulates transcriptional activity, or cooperative exchange, where the burst period is determined by transcription factors rapidly swapping positions due to cooperative binding (*25, 26, 28*).

Despite these challenges, the targeted engineering of biological systems advances rapidly, with standards and tools aiding in or automating their design being created (*11, 31–36*), while the awareness with respect to resources such as energy increases (*8–10, 37–39*). One branch of tools is Genetic Design Automation (GDA) software (*11, 36, 40, 41*). These tools solve the task of creating genetic logic circuits realizing Boolean functions with modules characterized in a gate library (*11, 40, 41*), the so called technology mapping. The term originates from the synthesis of electronic circuits. Compared to electronics, technology mapping in GDA needs to solve additional, more complex tasks for typically smaller circuits. Particularly, it may not draw gates from the library multiple times, since it needs to consider crosstalk among genetic components. GDA tools optimize the characteristics of genetic logic circuits by evaluating them with the use of models on the basis of *in silico* experiments. The primary objective is functionality, which expresses the circuits’ capabilities of implementing the Boolean function in terms of a distance related to the minimum fold-change between the two Boolean states (*11, 30*). To be meaningful, the employed models have to capture the characteristics inherent to the cells and constructs under consideration, such as the input-output characteristics of gene expression cascades, non-equilibrium attributes like energy dissipation, and transcriptional bursting.

Otero-Muras et al. (*42*) present a tool for automating the engineering of synthetic metabolic pathways on the basis of Pareto optimal designs. These metabolic pathways are defined by the user, with genetic logic circuits being a possible branch. iBioSim (*36, 43*) is a tool for the automatized construction of genetic circuits, their simulation and model representation. The absence of an automatized flow from Boolean specifications to a DNA sequence highlights that this tool does not primarily target the automated design of genetic logic circuits. Cello (*11, 40, 41*) implements a complete user interface for engineering genetic logic circuits. The user can specify the desired Boolean function using a traditional hardware design language. The given function is then transformed to a functional equivalent circuit which only uses elements of the provided libraries for *S. cerevisiae* and *E. coli*. The gate assignment is optimized to maximize the smallest fold change between circuit output values corresponding to the distinct Boolean states ON and OFF, the Cello-score. ARCTIC (*18, 30*) extends the focus on the robustness of the resulting genetic logic circuits. By introducing particle based simulation, the E-score, and structural variants, this tool accounts for the stochastic nature of genetic gates and extends GDA’s design space by circuit topologies. The 2023 version of this tool introduced context-awareness and accounts for effects such as crosstalk between transcription factors and non-cognate promoters, or titration of transcription factors to non-cognate binding sites.

While the awareness of energy, in particular, and resources, in general, increases (*8– 10, 37–39*), it is not yet part of GDA. The same holds for non-equilibrium models of gene expression, accounting for energy expenditure and sharpness beyond the Hopfield barriers (*12*). To this end, we introduce **Energy Aware Technology Mapping**, the design of genetic logic circuits with respect to functionality and energy efficiency. This is based on a probabilistic non-equilibrium steady state (NESS) model of gene expression, accounting for non-equilibrium characteristics like energy dissipation (*2, 3*) and promoter architectures varying in the number of binding sites, activation steps, and cognate transcription factors. To characterize gene expression, we derive the functional and energetic response curve as a function of transcription factor abundance. In particular, we relate the entropy production rate of our model to its thermodynamic energy dissipation rate.

Energy Aware Technology Mapping uses this model for the *in silico* simulation of genetic logic circuits. This allows us to explore the trade-off between functionality and energy efficiency for genetic logic circuits, where we introduce energy efficiency as the reciprocal of a circuit’s energy demand. To this end, we first consider the Boolean circuits presented in (*11*) and continue with the evaluation of the impact of structural variants and the associated Pareto fronts. The Pareto fronts give rise to the structure’s performance independently of the optimization objective considered. By presenting means for multi-objective optimization, we allow to trade-off the objectives function and energy efficiency in a joint optimization.

The results and methods we present here are implemented in the technology mapping framework ARCTIC. ARCTIC is available at https://www.rs.tu-darmstadt.de/ARCTIC.

## 2 Results and Discussion

### 2.1 An Energy Aware Gene Expression Model

Gene expression is an inherently non-equilibrium process (*1, 44*), depending on the presence of energy carriers and building blocks like ATP and charged tRNA. While gene expression is subject to regulation at various levels (*3, 45, 46*), we focus on transcriptional regulation through the binding of activating or inhibiting transcription factors. In this section, we present the response characteristics of a single protein-coding gene whose promoter has one or more binding sites for cognate transcription factors. To this end, we describe the dynamics of the promoter and of the abundances of its associated mRNA and protein via a stochastic chemical reaction network (CRN). In contrast to combinatorial (equilibrium) promoter models, the use of a kinetic model enables the description of energy-dissipating (non-equilibrium) promoters. In particular, we obtain the mean energy dissipation rate and the NESS mean and variance of protein abundance as a function of the transcription factor concentration. These non-equilibrium response curves are used to characterize the genes realizing the gates for the technology mapping and possess attributes like increased sharpness in comparison to their equilibrium counterparts (*12*). We will first elucidate the kinetic model and then introduce the thermodynamic concepts necessary to relate the kinetic model to its energy dissipation rate.

### Model Description as a Chemical Reaction Network

A CRN is composed of the chemical species X_1_, …, X_*N*_, and elementary reactions ℛ_1_, …, ℛ_*M*_ with stoichiometric balanced equations

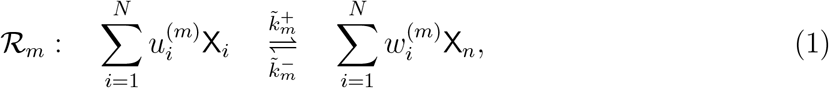

where the backward reaction microscopically reverses the forward reaction. Assu functions) 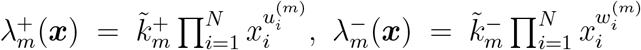 to each reaction ℛ_*m*_, where 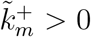 and 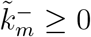 are the rate constants. A reaction is called microscopically reversible if 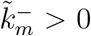. The concept of microscopic reversibility is important for the later thermodynamic treatment. For brevity, we simply write reversible instead of microscopically reversible in the following.

We partition the vector ***X*** of random variables into the promoter state ***Z*** and the RNA and protein abundances *R* and *P*. The promoter can switch reversibly between a finite number of states *z*_*i*_ and in each state RNA synthesis events occur according to the state’s transcriptional activity *a*_*i*_ ≥ 0. We represent the promoter states by different species, so that at any time ***Z***(*t*) is a unit vector with a one at position *i* in each case. RNA synthesis, protein synthesis, and degradation are modelled as single reactions, neglecting their multistep construction (*2*). Although each of the elementary steps of these synthesis and degradation reactions is reversible, we assume that the rate constants of the mechanistically reversed composite reactions are negligibly small, enforcing microscopic irreversibility. In Figures 2**A, B** we represent an exemplary instantiation of the gene expression model, where the promoter (**A**) has three transcription factor binding sites and two distinct levels of transcriptional activity. Given the concentration vector of the transcription factors ***c***, the transition rates of the promoter (as in Figure 2**A**) relate to Equation (1) as 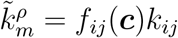 for ***ν***_*m*_ = ***e***_*j*_ − ***e***_*i*_, where *ρ* = + for *j > i* and *ρ* = − for *j < i*. The functions *f*_*ij*_ describe the dependence of the transition rate from state *z*_*i*_ to *z*_*j*_ on the transcription factor concentrations.

**Figure 1:**
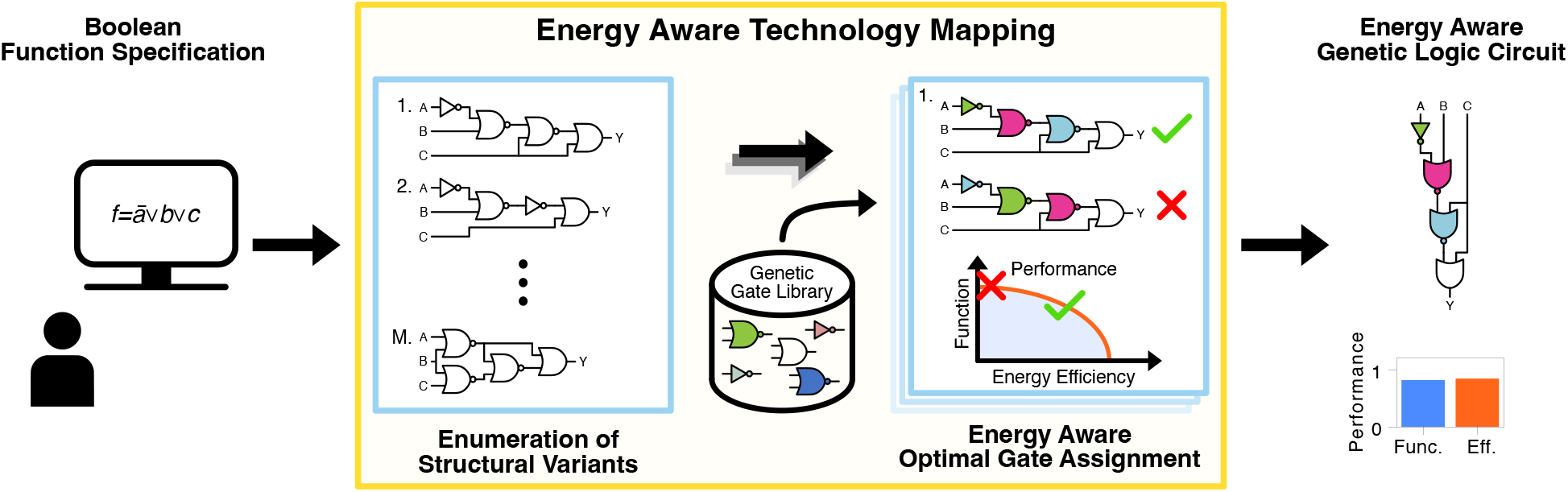
Energy Aware Technology Mapping. We here present the technology mapping pipeline as an example of the proposed Energy Aware Technology Mapping. With the specification of the Boolean function to realize as input, the technology mapping first enumerates structural variants (*30*). For each circuit structure obtained, we perform *in silico* an energy aware gate assignment. This takes into account the genetic logic circuit’s performance with respect to both, energy efficiency and functionality. After successful completion of the process, the user receives the automatically designed genetic logic circuit.

**Figure 2:**
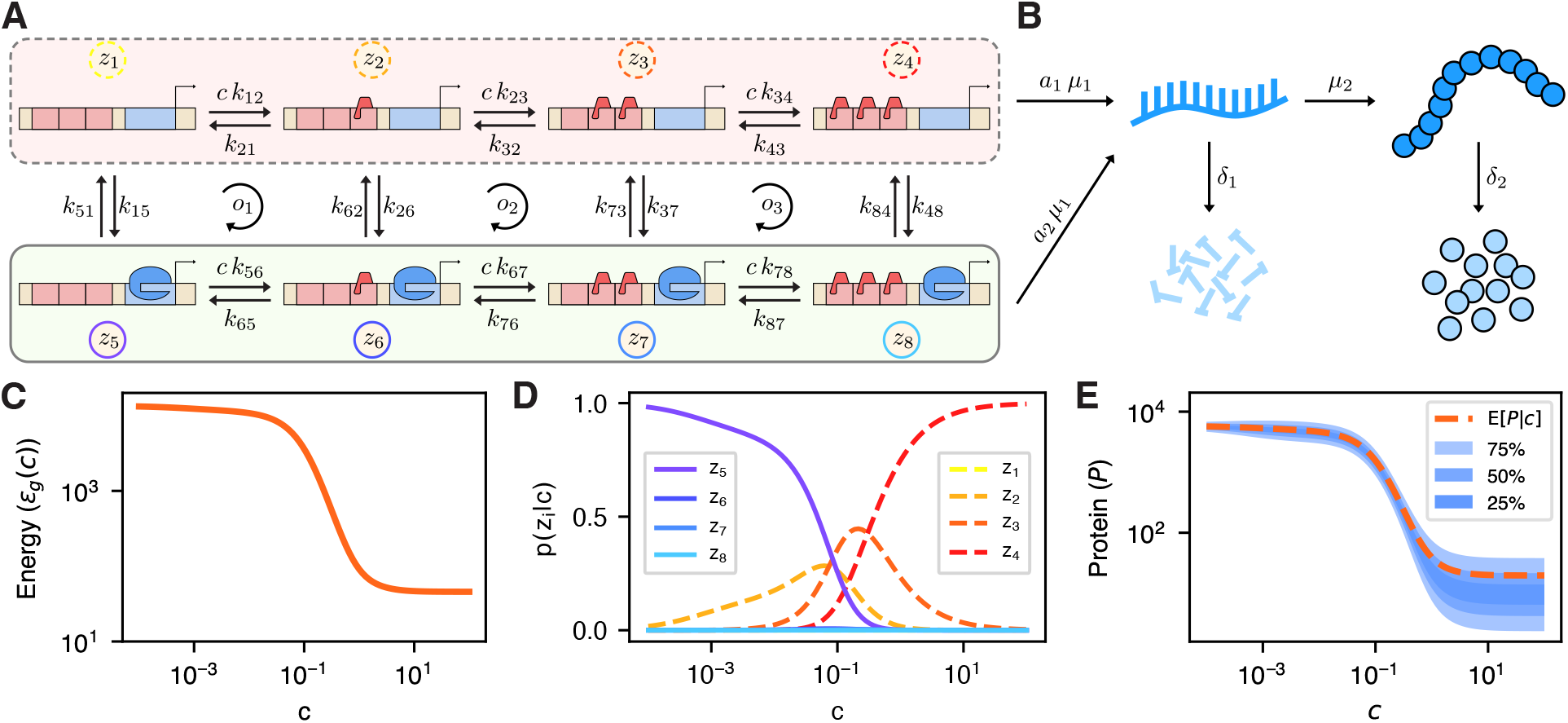
Energy aware gene expression model. **AB:** Schematic description of the proposed model consisting of the promoter model (**A**) and the reactions describing the RNA and protein dynamics (**B**). **A:** The promoter model is instantiated with two levels of transcriptional activity and allows for binding up to three transcription factors. The transcription factor concentration enters via the variable *c*, with the transcriptional active states (*z*_*i*_ with *i* = 5, 6, 7, 8) featuring transcription rate *a*_2_ *µ*_1_ and the inactive states (*z*_*i*_ with *i* = 1, 2, 3, 4) the basal rate *a*_1_ *µ*_1_ (*a*_2_ *> a*_1_). **B:** Besides transcription, the dynamics include translation (*µ*_2_) and the respective degradation reactions (*δ*_1_ and *δ*_2_). **C-E:** Exemplary response characteristic of the model as a function of the transcription factor abundance *c*. We here showcase inhibitory behavior of the transcription factor, while the model can also capture activatory behavior. **D** visualizes the steady state probabilities of each state, with the probability mass shifting from state *z*_5_ to *z*_2_ and *z*_3_ before concentrating in *z*_4_ as *c* increases. The corresponding protein distribution is shown in **E** by its mean and three quantile intervals. **C** presents the expected energy dissipation rate of the overall model. Comparing **C** and **E**, one notices the proportionality between energy dissipation rate and protein abundance.

The CRN representation of a promoter model is more versatile in general, as it may describe not only the number of bound transcription factors but also different DNA conformations in a potentially multi-step transcriptional activation (*12*). Such conformations can be, for example, open and closed loop complexes or different chromatin states (*12, 20, 48*). Since transcriptional activity is independent of the binding states of the transcription factors, this model can represent transcriptional bursts in both the multistep activation model and the exchange model (*12, 25–29*). By turning the reaction rate constants themselves into random variables, one can extend this model to include extrinsic noise due to external context factors in the sense of (*49–51*). It should be noted that reversible ATP-dependent reactions ATP ⇌ ADP + P_*i*_ always produce ATP in the reverse direction. Consequently, if any promoter state change is predominantly ATP-consuming in one direction, but not ATP-producing in the reverse direction, then there are at least two distinct reactions involved.

### Kinetic Response Curve of a Single Gene

The evolution of the distribution of the state of the gene expression system p(***X***(*t*) = ***x***) is described by the chemical master equation given in SI Section S1.3. Using the Chapman-Kolmogorov backward equation, we derive the mean and variance of the non-equilibrium steady state RNA and protein distributions in SI section S2, and derive the promoter’s steady state distribution ***π***(***c***) = lim_*t*→∞_ p(***Z***(*t*) = ***e***_*i*_) by applying the methods in section 4.2 to the promoter’s propensity matrix **Λ** = **Λ**(***c***) with Λ_*ij*_(***c***) = *f*_*ij*_(***c***)*k*_*ij*_. Together with the transcription activity parameter *a*_*i*_ associated with each promoter state *z*_*i*_, the average promoter activity is

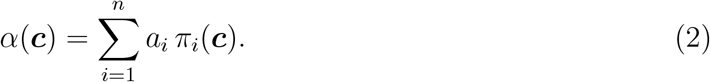

The protein’s mean and variance and the RNA’s mean are given by

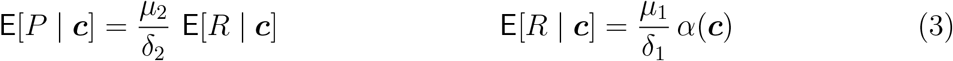

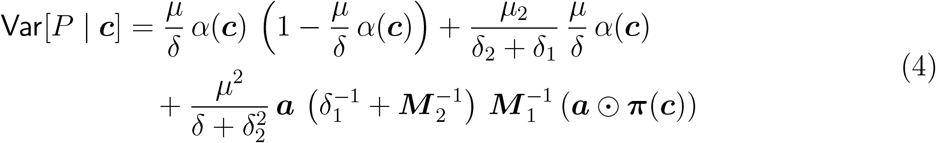

where *µ* = *µ*_1_ *µ*_2_, *δ* = *δ*_1_ *δ*_2_, ***a*** = [*a*_1_ … *a*_*n*_], ⊙ is the Hadamard product, and ***M*** _*i*_ = (*δ*_*i*_ ***I***_*n*_ − **Λ**^*T*^) with ***I***_*n*_ the *n* times *n* identity matrix. The quantities in Equations (3) and (4) are the characteristic response curves of a gene depending on the transcription factor concentrations ***c***. Later on, we use them for the NESS simulation of the gene circuit realizing the genetic logic circuit within the technology mapping. To allow for energy awareness in the technology mapping, we proceed by establishing a link between the kinetic model and its energy dissipation rate to derive an energetic response curve as a function of transcription factor abundance ***c***.

### Stochastic Thermodynamics of Open CRNs

In stochastic thermodynamics, each state ***x*** of the system is associated with its Gibbs free energy *g*(***x***). Upon reaction ℛ_*m*_ in forward direction, the Gibbs free energy change is 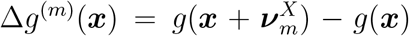, where 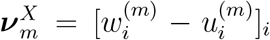 is the stoichiometric change vector. Further, the system is assumed to be in contact with a heat bath of temperature *T*. A system consisting of a CRN and a heat bath is called a closed CRN, which -following the zeroth law of thermodynamics-relaxes to equilibrium (*52*). In equilibrium, the mean energy dissipation rate is zero (*47*). A system that exhibits a zero net change in Gibbs free energy along any closed cycle in its state space can be considered a closed system. For instance, this is the case in the promoter in Figure 2**A**, if none of the transitions require additional energy carriers. Otherwise these energy carrying species need to be accounted for in the stoichiometric equations. Furthermore, for the dynamics of RNA and protein, it is necessary to take into account the building blocks whose recycling after degradation requires the expenditure of energy.

We thus assume that the reactions ℛ_*m*_ actually include species Y_1_, …, Y_*L*_, which are controlled by the cell to always have constant concentrations. Usually, these are energy carriers like ATP and its hydrolysis products like ADP and P_i_. By coupling our CRN to these species, we obtain an open CRN described by the stoichiometric balanced equations

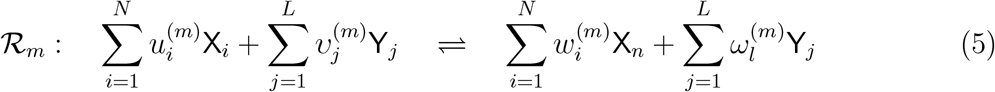

where, the species Y_1_, …, Y_*L*_ act as chemostats. Figure 3 illustrates the composition of closed and open CRNs. The Gibbs free energy change of the chemostat species associated with reaction ℛ_*m*_ is 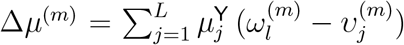, where 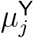 is the chemical potential of species Y_*j*_. Identifying 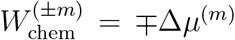 as the chemical work done on the core system by the chemostat upon reaction ℛ_*m*_ in the forward and backward direction, respectively, we introduce the energy dissipated into the environment per reaction as the difference of chemical work and Gibbs free energy change

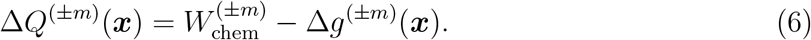

The mean energy dissipation rate 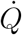 is then given by

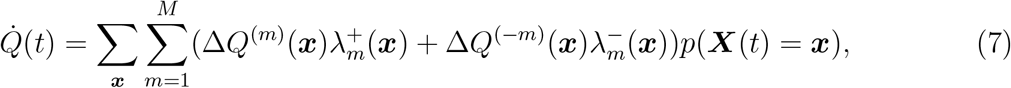

as we show in SI Section S3.2. For open CRNs the propensity functions of any reversible reaction ℛ_*m*_ (i.e. 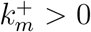 implies 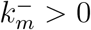) are related to the energy changes by the thermodynamic consistency relation (*52*)

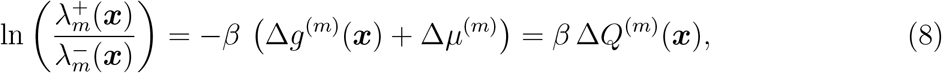

which associates the rates of the reactions with the corresponding energy dissipation and *β* = (*k*_B_*T*)^−1^ parameterizing the heat bath. If all reactions of an open CRN {ℛ_1_, …, ℛ_*M*_ } are reversible, then we may use Equation (8) to identify the energy dissipation rate with the entropy flow into the environment

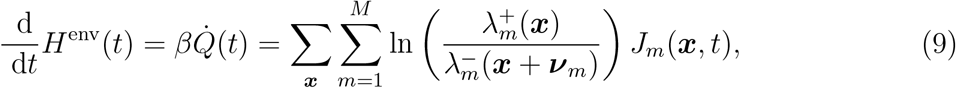

where 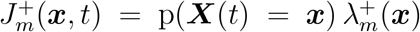 and 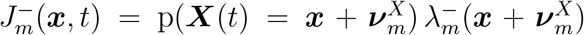 represent the forward and backward probability fluxes of reaction ℛ_*m*_ with net flux 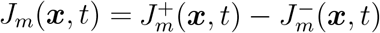.

**Figure 3:**
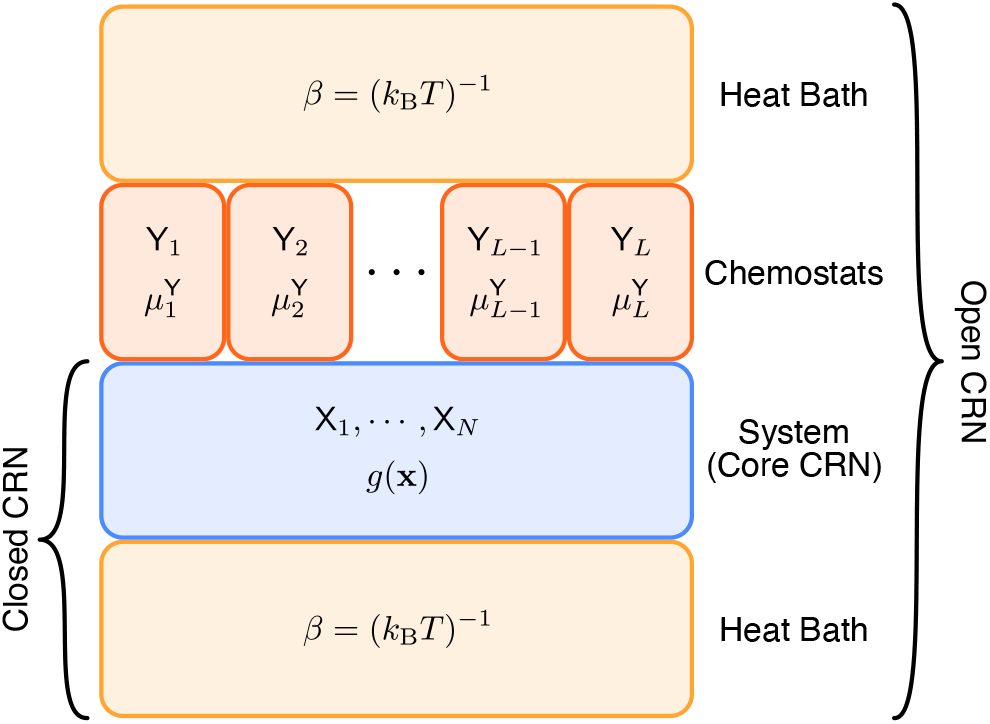
Schematic description of an open CRN. Contrasting open and closed chemical reaction networks, one observes well the difference sourced in the addition of chemostat species Y_*i*_. The abundance of the chemostat species is kept constant, in our case by the cellular environment, and leading to a chemical potential driving the core CRN. In this work, the chemostat species refer to cellular energy carriers like ATP and the products of corresponding hydrolysis reactions.

### Relation Between Energy Dissipation and Entropy Production Rate of the NESS

The entropy production rate (*47, 53, 54*) is the sum of the entropy change rate of the system and the entropy flow into the environment, i.e., 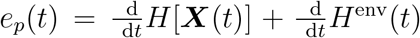, where *H*[***X***(*t*)] = − Σ_***x***_ *p*(***X***(*t*) = ***x***) ln(*p*(***X***(*t*) = ***x***)). At steady state (or in the limit *t* → ∞) we have 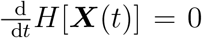 and hence the mean energy dissipation rate (in units of *k*_B_ *T*) of the NESS is given by the entropy production rate, i.e.,

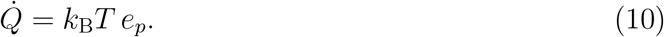

Due to the central relevance of this relationship, we recommend SI Section S3 and in particular Section S3.3 to the reader, where we derive and discuss the relationship between entropy production rate and energy dissipation rate in depth.

### Energetic Response Curve of a Single Gene

Following the above, we express the energetic response curve to the transcription factor concentrations ***c*** in terms of the overall mean energy dissipation rate of the NESS *ϵ*_*g*_(***c***).

This rate is given as the sum

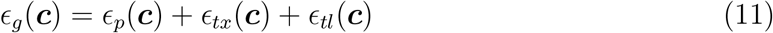

of the contributions of the promoter *ϵ*_*p*_(***c***), the RNA dynamics *ϵ*_*tx*_(***c***), and the protein dynamics *ϵ*_*tl*_(***c***).

In particular, we use the relationship between energy dissipation and entropy production to quantify the energy dissipation rate of the promoter as *ϵ*_*p*_(***c***) = *k*_B_*T e*_*p*_, since it is assumed to satisfy the microscopic reversibility requirement. Note that since the entropy production rate is calculated without the knowledge of chemical potentials 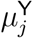 or the Gibbs free energy changes Δ*g*^(*m*)^, the expression is valid for a variety of different promoter structures, provided the promoter topology is accurate. To derive the NESS entropy production rate of the finite state promoter, we make use of Schnakenberg’s method (*53*), which we outline in SI Section S4. Using this method we obtain

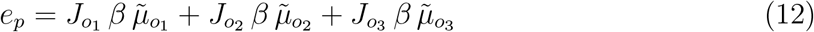

as derived in SI Section S4.1 and consequently the promoter’s energy dissipation rate is *ϵ*_*p*_(***c***) = *e*_*p*_*/β*. Here, 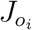 denotes the probability flux along the cycle *o*_*i*_ as presented in Figure 2**A** and 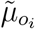 is the change in chemical potential associated with a single run through cycle *o*_*i*_. The products of each flux 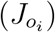 and force 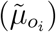 pair resemble the well known expression for electrical power, i.e., the product of a voltage (force) and the induced electric current (flux).

Since our transcription and translation model lacks microscopic reversibility, the thermodynamic consistency relation (Equation (8)) is not well defined. However, under the assumptions given in SI Section S3.4, we only need to know the chemical work per reaction in addition to the rate constants to compute the energy dissipation of those reactions via Equation (7). Here we derive these quantities from an intuitive heuristic perspective, while the corresponding considerations involving Equation (7) are presented in SI Section S3.4.

Considering the energy required per RNA molecule first, we subsume all length dependent energy requirements for RNA synthesis and degradation in *e*_*r*_ and the length independent ones in 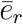. Taking the RNA length *l*_*r*_ in nucleotides into account, the energy per RNA molecule is 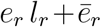. For the protein synthesis and degradation, we define the energies *e*_*p*_ and 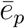 analogously and set *l*_*p*_ as the protein length in amino acids. To obtain the actual energy dissipation, we have to account for the synthesis and degradation rates. In particular, these rates are given by *δ*_1_ E[*R* | ***c***] and *δ*_2_ E[*P* | ***c***] for RNA and protein, respectively. Multiplying the rate of molecule synthesis and degradation with the associated energy yields

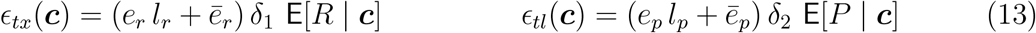

as the expected energy dissipation rates of the RNA and protein dynamics.

#### 2.2 Energy Aware Technology Mapping

With the energy aware NESS model of gene expression at hand, we now present the steps for incorporating it into the technology mapping process. This gives rise to Energy Aware Technology Mapping.

### From Genetic Logic Circuits to Genes and Back

The efficient technology mapping of genetic logic circuits gets enabled by large scale *in silico* experiments. Depending on the focus, *in silico* evaluations can target different levels of abstraction as exemplified in Figure 4**A**. These abstractions not only differ with respect to the modules they are composed of, but also in the signals considered. While Boolean functions take Boolean signals into account, the genetic logic circuits we consider here use the promoter activity in RPU, with protein abundance serving as signal carrier in the underlying gene circuits. With the model we present in this work, we target the gene expression level, where genes express proteins acting as transcription factors and repressing the expression level of the genes associated to their cognate promoter. As such, this approach can reassemble the actual implementation of a genetic logic circuit within cells as presented in (*40*).

**Figure 4:**
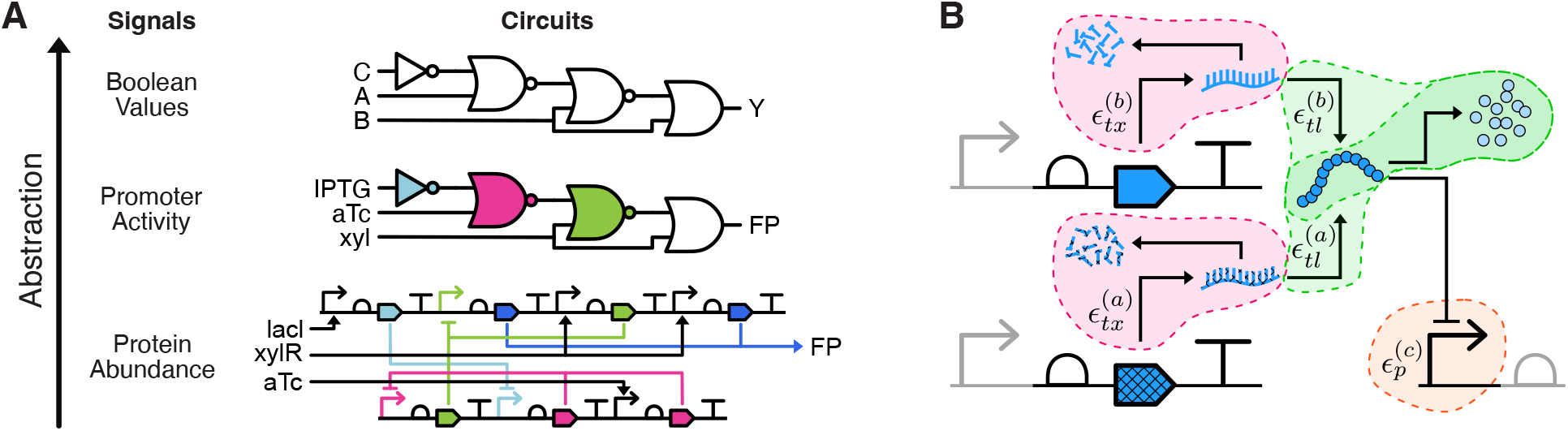
From genes to logic circuits. **A:** Visualization of the abstraction levels encountered in genetic design automation (GDA). The gene circuit at the bottom realizes the behavior within the cell and is the circuit our model is applied to. It relies on protein concentrations as signal carriers. Defined on top, the genetic logic circuit provides a convenient interpretation in terms of gates, familiar to engineering disciplines and the basis for GDA. The Boolean logic circuit shadows all implementational details and presents the function to realize. **B:** Overview on the energy demands of a genetic NOR gate following the implementation of (*40*). The gate consists of the genes *a, b*, and *c*, with the preceding and succeeding gates greyed out. *ϵ*_*tx*_ denotes the energy dissipation rate of the RNA dynamics, *ϵ*_*tl*_ of the protein dynamics, and *ϵ*_*p*_ is the energy dissipation rate of the promoter.

To make use of our NESS gene expression model within the genetic logic circuit centered technology mapping, we first obtain the corresponding gene circuit. In the next step, the Boolean input values are translated to the respective promoter activities, from which we then derive the corresponding inducer concentrations. The gene circuit is evaluated by applying our model in topological order to the genes. By deriving the promoter activity associated to the abundance of the reporter protein, we finally obtain the output representation at the genetic logic circuit level.

### The Functional Performance of a Genetic Logic Circuit

In order to complete the loop, we could apply a thresholding to the promoter activity values and obtain the associated Boolean value. Within the technology mapping, the performance of this thresholding approach is subsumed in the score *S*, effectively measuring the distance between activity levels representing *ON* and *OFF* states. Formally, we define the genetic logic circuit as the tuple (*γ, q*), where *γ* is the circuit’s structure and *q* the corresponding assignment of genetic gates. We derive the score *S* = *S*(*γ, q*) by evaluating the genetic logic circuit *in silico* for all Boolean input conditions ℐ (with 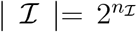 for *n*_*I*_ inputs). The obtained circuit output values in terms of promoter activities are subsumed in 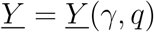 in case they shall represent an *OFF* state and 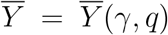 otherwise. Identifying the elementary scoring function with *s*, the score *S*(*γ, q*) is given by

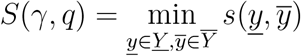

where 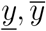 can be scalar values as in the case of the cello-score (*11*) or empirical distributions as for the E-score (*30*). Within Section 4.4, we extend on the definition of the E-score and it’s application to empirical distributions.

### The Energy Dissipation of a Genetic Logic Circuit

Besides the functional characterization, our gene expression model allows for insights into the energetics of the genetic logic circuit. For this, we define ℒ (*γ, q*) as the set of genetic logic gates included in the circuit (*γ, q*). Furthermore, we identify the genes expressing the transcription factor associated to gate *l* ∈ ℒ (*γ, q*) with the index set 𝒢_*C*_(*l*) and the genes with cognate promoters with the index set 𝒢_*P*_ (*l*). Again, the energy dissipation rate is the aggregation of the single parts contributions, resulting in

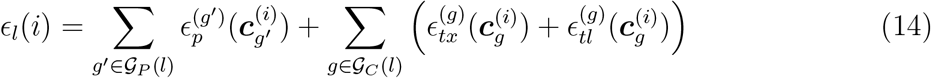

where *i* ∈ ℐ identifies the Boolean input condition and 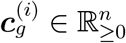 denotes the cognate transcription factors’ concentrations of gene *g* for the respective input condition. Within Figure 4**B**, we present this as an example of a genetic NOR gate following the implementation of (*40*). Taking all gates ℒ(*γ, q*) of the genetic logic circuit into account, the energy dissipation rate of the whole circuit for the *i*th input assignment is

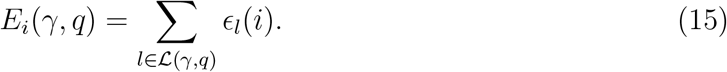

We define the average and maximum expected energy dissipation rates *E*(*γ, q*) and *E*_*max*_(*γ, q*) of the genetic logic circuit (*γ, q*) as

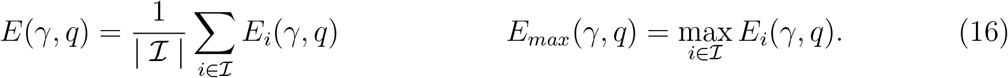

Depending on the type of application and the constraints enforced, both, *E* and *E*_*max*_, are valid objective functions for the energetic optimization. In the case of stochastic evaluation, these aggregation functions extend naturally.

### Technology Mapping of Genetic Logic Circuits

The previous two sections have introduced different metrics for the evaluation of genetic logic circuits. It is the task of the technology mapping to design a topology from the available gate types that realizes the desired Boolean function. To this end, our tool systematically enumerates all possible variants of gate selection and topologies. For each of these possible solutions, a gate assignment has to be carried out, which is then heuristically optimized. To incorporate both metrics into the scoring of the resulting circuits, either a multi-objective or a constrained optimization can be done.

#### 2.3 The Genetic Logic Circuit’s Trade-Off between Energy and Function

With the energy aware gene expression model included into the technology mapping framework ARCTIC (*18, 30*), we can explore the design space of genetic logic circuits with respect to both, functionality and energy efficiency. For this purpose, we consider the genetic logic gates introduced with Chen et al. (*40*). In particular, we calibrate the parameters of our model to capture the response characteristics encoded in the cytometry data provided by (*40*) as described in Methods 4.3. We then use the obtained parametrizations to derive a genetic gate library for *S. cerevisiae*, consisting of 12 genetic logic gates utilizing nine independent transcription factors.

We start our evaluation by investigating the energy dissipation rates encountered in mapping logic circuits and their relation to the circuit’s functionality. Based on the insights gained, a systematic evaluation of the circuit structure’s effect on the energy of genetic logic circuits follows up. In this context, we also consider the Pareto fronts and the application of multi-objective optimization. To emphasize comparability among genetic logic circuits as well as function and energy, we introduce the energy efficiency, or briefly efficiency, 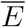 as the inverse of the energy measure (i.e. 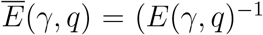 or 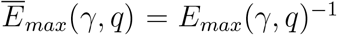)and apply normalization to the best scores encountered in the considered benchmark set.

### Function and Energy Optima are Disjoint

To assess the energy landscape of genetic logic circuits, we evaluate the technology mapping results of 33 logic circuits, each realizing a different Boolean function. The choice of circuits follows (*11*) and encompasses circuits ranging from two to seven gates. For the purpose of this evaluation, we optimize each circuit for the two objectives function and energy and constrain the other in the respective case. While the minimum functional score to achieve is identical for all circuits, the upper limit on the energy dissipation rate during functional optimization is chosen in dependence to the gate count (see Methods 4.4).

Starting with a small circuit first, Figure 5**A** presents the function and energy optimized genetic logic circuits for 0×2F (3 gates). Despite the same structure, the different gate assignments feature significant differences in function and energy. In particular, the energy optimized version decreases the functionality by 70.7% to improve energy efficiency by 44.4% in comparison to the version optimized for function. This is a decrease in fold-change by a factor of 3.4 and a decrease in energy resource consumption by 30.8%. In addition, the disjoint optimality of function and energy indicates a trade-off between the two optimization goals considered. Increasing the number of gates does not change this impression, as we observe by considering the results for circuits 0xDF (4 gates), 0×20 (5 gates), and 0×81 (7 gates) in Figure 5**D**. However, the absolute level of energy dissipation changes in both, the functionally and efficiency optimized circuits. While we estimate an energy dissipation rate of 29, 401 *k*_B_ *T s*^−1^ for circuit 0×2F, this increases to 38, 251 *k*_B_ *T s*^−1^ for circuit 0xDF, 40, 901 *k*_B_ *T s*^−1^ for circuit 0×20, and 64, 865 *k*_B_ *T s*^−1^ for 0×81 in the energy optimized case. With an average contribution of 98.8%, our model predicts protein synthesis and degradation to account for the largest portion of energy expenditure.

**Figure 5:**
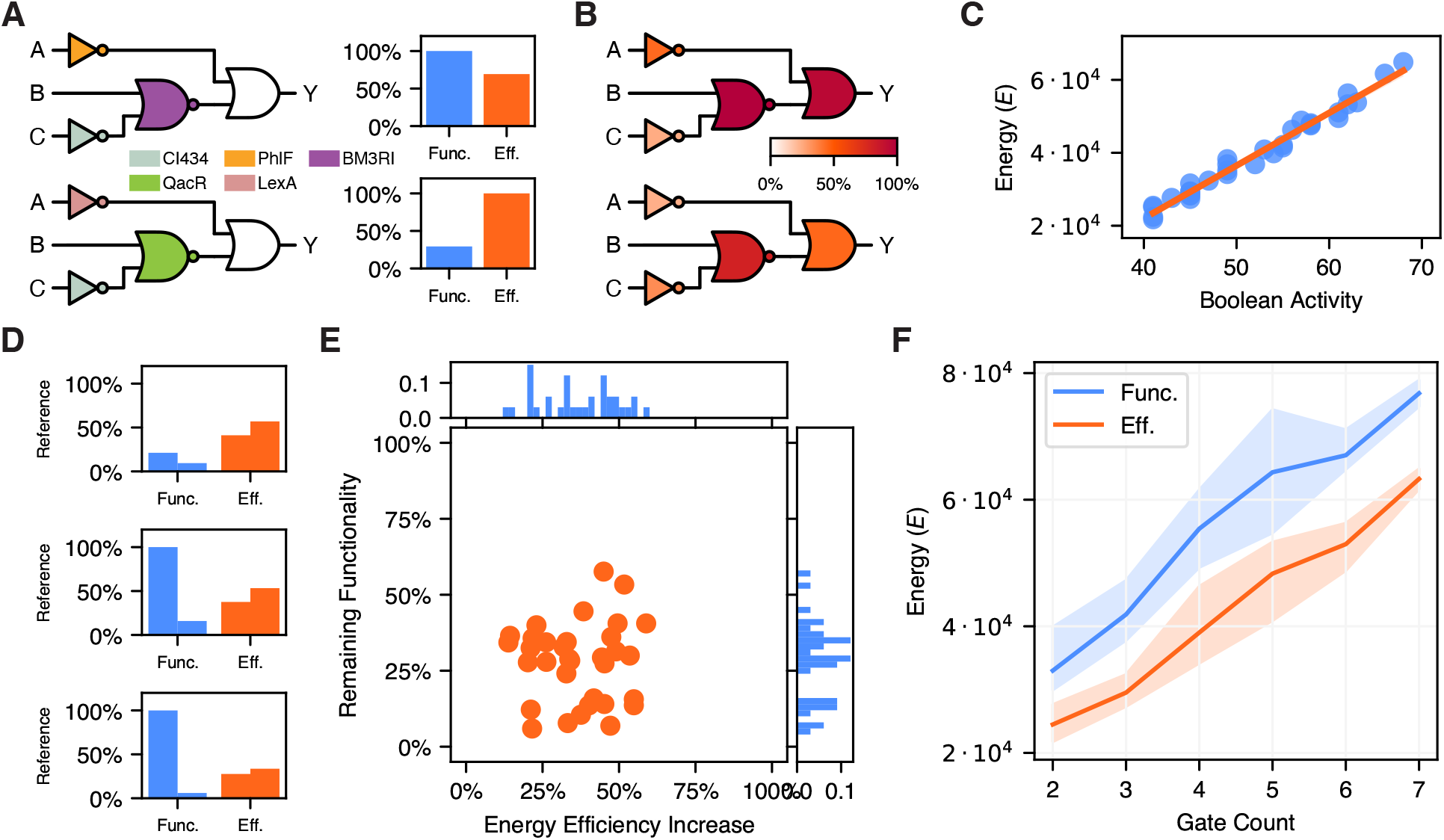
Function and energy as disjoint optimization targets. **A:** Circuit 0×2F (3 gates) is optimized for function (top) and energy efficiency (bottom). The bar plots on the right present the performance of the genetic logic circuits depicted on the left for both evaluation criterions. Clearly, the optima are disjoint. **B:** Visualization of the mean energy dissipation rate per gate (lower is better) for 0×2F over all Boolean input conditions, optimized for functionality (top) and energy efficiency (bottom) and normalized to the largest value. The energy dissipation of the NOR and OR gate differ significantly between the two implementations, impacting the overall energy efficiency significantly (see Figure **A**). **C:** Boolean activity as an heuristic for the energy of a genetic logic circuit. Here compared to the energy of circuits with the objective energy efficiency. **D:** The optimization results of circuits 0xDF (4 gates), 0×20 (5 gates), and 0×81 (7 gates), normalized to the maximum functional score and energy efficiency observed in the benchmark. The left of a bar pair refers to the results of the functional and the right to the energetic optimization. Again, optimizing for either of the two objectives decreases the genetic logic circuits performance in relation to the other. **E:** Analysis of the cost of energy efficiency increase on the benchmark. While the optimization for energy improves energy efficiency up to 58.9%, the functionality score is often decreased significantly. **F:** Relationship between expected energy dissipation rate of the circuit and it’s gate count. The bold line is the mean and the shaded areas present the minimum and maximum intervals, with the colors indicating the optimization objective. This figure reveals a near linear relationship between energy dissipation rate and gate count. However, the optimization objective determines the offset. Comparing the objectives, optimization for energy efficiency allows to implement genetic logic circuits of six gates with the energetic requirements of functionally optimized four gate circuits.

Figure 5**B** provides information on the differences of energy dissipation for the respective optimization goals. Presenting the energy per gate averaged over all input assignments, we observe that the two NOT gates differ only slightly between the functionally (top) and energetically (bottom) optimized variants. With 61.8%, the OR gate conduces the majority of energy saving. However, this is a direct result from the change in gate assignment of the respective preceding gates, as in comparison to the functionally optimized version their maximum expression levels are roughly halved. In consequence, the circuit’s score, which assesses the fold-change, drops proportionally. For functions 0xDF, 0×20, and 0×81, we observe the same behavior, with most energy savings resulting from gates closer to the output and only minor improvements and sometimes even worsening from gates close to the input of the circuit. Thereby, the largest energy saves are achieved by reducing the expression levels of Boolean ON states significantly.

Shifting the perspective to the set of all circuits, the optimization for energy increases energy efficiency up to 58.9% and on average by 37.2%. This is possible as the efficiency of most of the circuits can be improved well (see Figure 5**E**). However, this comes at a cost, as there is often only a small gap between the functionality constraint and the actual functionality score, as observable in the average remaining score of 28.5%. Figure 5**E** highlights this by visualizing the relationship between function and energy efficiency for the two cases of optimization. Reconsidering the relation between circuit size and energy, Figure 5**F** indicates the linear relationship between these quantities for the circuits taken into account for both optimization objectives. Comparing the results for efficiency and functionally optimized circuits among different circuit sizes, we see that a fixed energy budget allows for larger circuits when optimized for energy efficiency. On the other side, circuits designed by Energy Aware Technology Mapping feature an energetic advantage of at least one gate.

These results indicate an inherent trade-off between the two objectives energy efficiency and functionality for genetic logic circuits. Considering the *in vivo* realization of circuits, the expression level based signalling requires sufficient fold change to differentiate between distinct logic levels. In the presence of basal expression, fold change is achieved by sufficiently high levels of protein abundance, with the energy expenditure increasing proportionally. As part of the optimization for energy efficiency, the expression levels are reduced to a minimum viable level. However, the lower limit on the score ensures that despite the drop in functional performance, the circuit’s function is preserved. The observation of highest energy savings at the circuit outputs can result from level separation as a requirement for the function of gate cascades (*11, 18*) and minor optimization potential at the circuit inputs. Yet (*18*) points out that these last gates are the most important for the functional performance of a circuit. Again, this points to the exclusivity of the objectives considered. An aspect not explicitly considered yet but relevant for the overall energy expenditure is the gate technology used. The gate library used (*40*) provides transcription factor based gates, requiring the expression of heterologous proteins. Since our model and other studies (*5, 9, 55–57*) consider heterologous protein expression rate proportional to energy expenditure and protein dynamics dominate the energy consumed by gene expression, RNA-based gates could significantly reduce the overall energy consumed.

Driven by these insights and the relation between circuit size and energy dissipation, we present an even more precise estimate of a genetic logic circuit’s energy dissipation rate. This estimate is given by the **Boolean activity** of a structure, which reduces the estimation to counting the states which require protein expression. More formally, we consider the truth tables of the gates constituting the circuit structure. For each NOT and NOR gate, the number of Boolean ON states of the respective input gates feature is counted. To the circuit’s inputs and outputs, special treatment applies. The inputs are counted as if they were always ON, as their *in vivo* implementation makes use of constitutive promoters. For the outputs (simple or implicit OR), the number of ON states is counted as this refers to the protein level ideally representing this state. The overall Boolean activity is then obtained by summing up all the individual contributions for all possible Boolean input conditions. Figure 5**C** presents the relationship between the Boolean activity and the energy dissipation rate of genetic logic circuits optimized with respect to energy efficiency. With a Pearson correlation coefficient of 0.989 (0.940 when optimized for functionality), this easy-to-use heuristic is highly predictive.

### Promoter’s Energy Expenditure peaks in Transition Region

The circuit level trade-off is dominated by the protein expression level. In order to gain insight into the gate respectively promoter level, we focus here on the promoter energy, the entropy production rate. For two exemplary genetic gates, Figure 6 presents the entropy production rate and average promoter activity as a function of transcription factor abundance. All gates (see Figure S7) feature a peaked entropy production rate, concentrating energy dissipation mostly to the transition region of the gate’s response curve and being more peaked the sharper the transition is. In addition, the entropy production rate is often highest (dashed line) close to the steepest descend (marker) of promoter activity (see Figure 6**A**), with Figure 6**B** presenting a counterexample to this.

**Figure 6:**
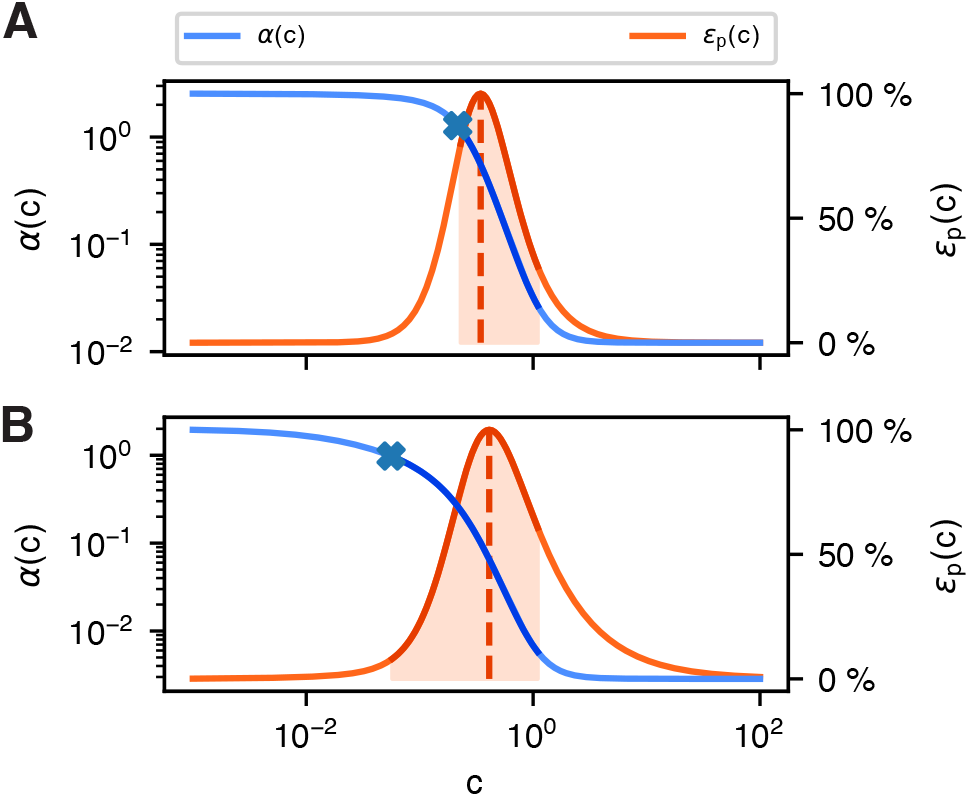
Peaked entropy production rate in transition region. The average promoter activity and the corresponding entropy production rates of two exemplary promoter models as a function of transcription factor abundance *c*. We highlight the transition region, which connects the two saturated regions of promoter activity. In most cases, entropy (*ϵ*_*p*_(*c*) = *e*_*p*_(*c*) *β*^−1^) is highest (dashed line) within this transition region. Besides, this entropy production peak is often close to the steepest descend (marker) of promoter activity (see **A**) but not in all cases (see **B**).

Considering the saturated regions for low and high input transcription factor levels, the promoters’ energy dissipation rates decrease significantly. In the context of genetic logic circuits, these regions represent the Boolean states one wants the gates to attain, giving rise to only minor contributions of the promoters to the overall energy dissipation of well functioning genetic logic circuits. Despite this, 21 of 33 efficiency optimized circuits feature a lower promoter energy dissipation rate in comparison to their functionally optimized pendants.

### Circuit’s Structure shapes Energy Dissipation

The previous results indicate a strong connection between a circuit’s size and its energy dissipation rate. However, not all circuits of a distinct size perform equally well, which brings the genetic logic circuit’s structure into focus. The structural variants approach implemented in ARCTIC (*18, 30*) allows for optimizing the structure in addition to the gate assignment. In doing so, (*30*) demonstrates the benefits of including the circuit’s structure in the optimization process for functionality. Guided by these insights, we use the structural variants approach to evaluate the structure’s impact on circuit performance with respect to both, functionality and energy efficiency.

For the evaluation of the structural variants, we consider exemplary the three Boolean functions 0xEC (3 gates), 0×02 (4 gates), and 0xE7 (5 gates), and allow for a single excess gate compared to the minimal structure. We apply the same constrained optimization as in the previous section, but optimize only for energy efficiency. Figure 7**A** provides a first intuition of the effect of structural variants by presenting the results of function 0xEC. The 16 different structures differ significantly in their functional but also energetic characteristics. The worst structure has a 54.3% higher energy dissipation rate than the best, and on average, the structural variants improve energy efficiency by 30.7%. Among the presented variants also the functionality varies, with an observed maximum improvement of 52.4%. Even more interesting is, that some structures (i.e. 11, 13, and 15) are significantly better than others in terms of both, energy efficiency and functionality.

**Figure 7:**
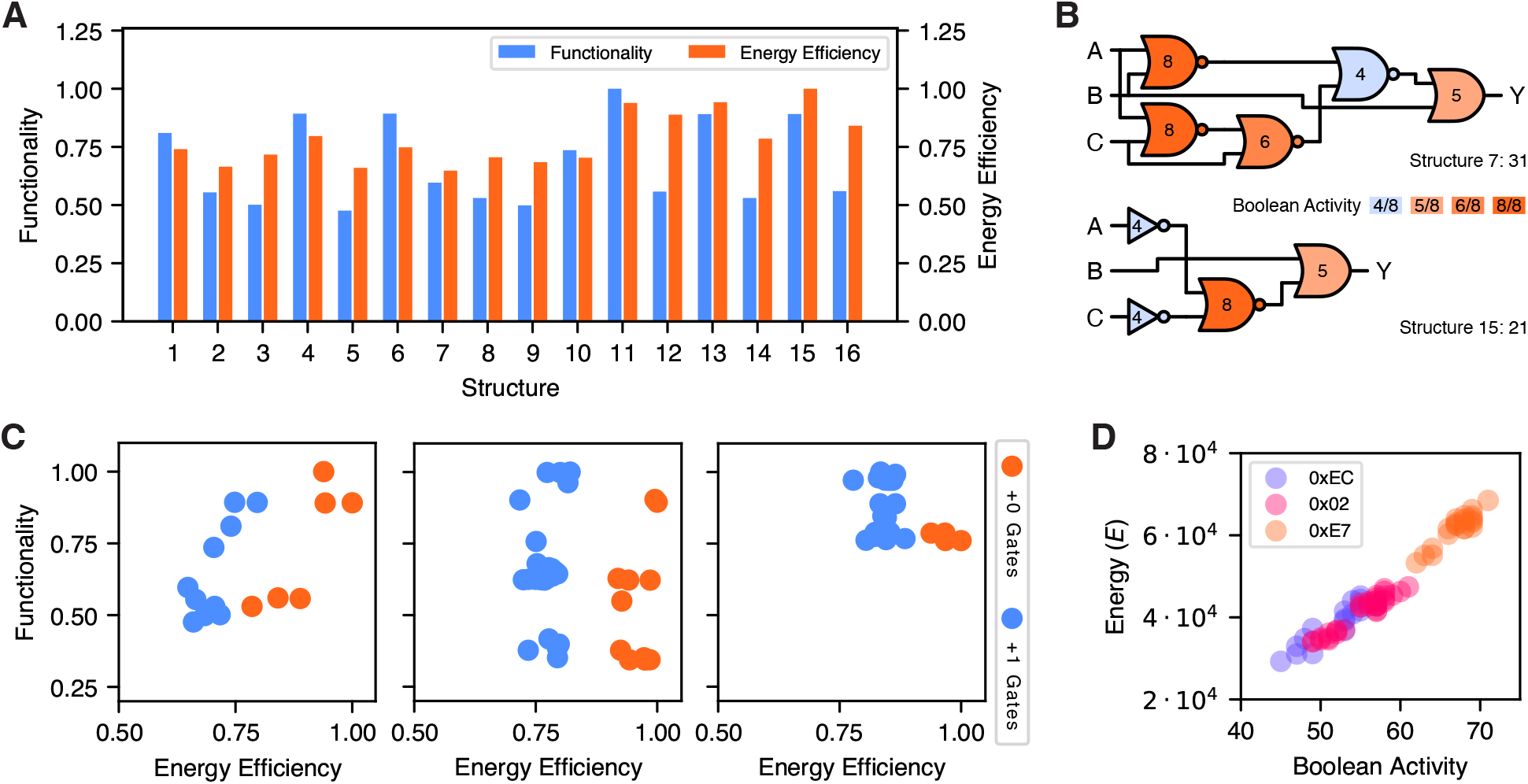
The logic circuit structure’s impact on energy efficiency. **A:** We here present the functionality and energy efficiency of the structural variants of function 0xEC optimized with respect to energy efficiency. The structures feature different characteristics in both, the energy efficiency and functionality, with an average improvement of energy efficiency by 30.7% when considering the best structure. Normalization refers in both cases to the best value encountered. **B:** The Boolean activity per gate for structures 7 and 15 of function 0xEC, which exhibit the worst and best energy efficiency. In comparison to structure 7, structure 15 features 10 active states less. This manifests in a significantly higher energy efficiency of structure 15 (see **A**). **C:** Overview on the distribution of energy optimized variants for the three functions 0xEC (3 gates), 0×02 (4 gates), and 0xE7 (5 gates), where the gate counts refer to the smallest structure obtained. Within the figures, the colors code for the number of excess gates, showing that smaller structures are beneficial for energy efficiency. However, for larger circuits the consideration of excess gates proves beneficial for functionality. The values are normalized with respect to the best values obtained for each function, while the clustering results from the discrete nature of gate assignment. **D:** Also for the structural variants, the correlation between Boolean activity and the energy dissipation is significant albeit varies among the different functions (0.927 for 0xEC, 0.96 for 0×02, and 0.934 for 0xE7).

Figure 7**B** provides a detailed insight into the Boolean activity of the least (7) and most (15) energy efficient structures. For each gate, its accumulated Boolean activity over all Boolean circuit input assignments is depicted. The difference of 10 active states between the two circuits manifests itself in a significantly differing energy efficiency. Examining the structures in detail, two causes for this difference can be found. First, structure 15 features one gate less compared to structure 7. Second, the way in which the Boolean function is computed leads to a lower average activity of the gates in structure 15 (6.2 for structure 7 vs. 5.25 for structure 15). This is partly caused by the usage of NOT gates over NOR gates in structure 15. In the given gate architecture, NOR gates feature an independently expressed transcription factor for each input, leading to a potential doubling of energy expenditure in one of four output states.

To emphasize our understanding of the relationship between energy efficiency, functionality and structure, Figure 7**C** includes for each of the considered functions a visualization showcasing the distribution of values. Structures with less gates are more energy efficient than their larger counterparts. However, except for 0xEC, the larger circuits feature the better functional performance despite the smaller ones being rather close. Considering the energy efficiency improvements possible, these decrease with increasing circuit size as the average improvement of 0×02 is 21.4% and of 0xE7 16.1%. Besides, some structures lead to superior designs with respect to both objectives.

Reconsidering the Boolean activity, we here investigate whether this heuristic is expressive for structures with the same Boolean function. Presented in Figure 7**D**, Boolean activity and energy expenditure again correlate significantly, with a correlation coefficient of 0.982. However, when considering single functions, the correlation reduces to 0.927 in the worst case.

The predictive power of Boolean activity and the dependence on the Boolean function indicate a circuit structure’s relevance to energy efficiency. We here confirms this by pointing out the improvements possible by considering structural variants in the context of energy efficiency. As the structures differ in the number of states requiring protein expression, they allow for increasing energy efficiency and can be beneficial for functionality.

### Pareto Optimality among Structures

Until now, we only considered the extremes of being either energetically or functional optimal. In a design approach, one would trade-off these objectives to achieve the best performance in relation to the energy spend. To this end, we analyze the Pareto front of all the structures for the Boolean function 0xF7. In particular, we perform a parameter sweep over the energy constraint for each of the 10 structures. This gives rise to the functional best genetic circuit adhering the energy constraint. The constraint is chosen to equally distribute into 20 samples over the interval of energy levels identified by an initial random sampling.

The Pareto fronts obtained showcase the exclusiveness of energetically and functional optimality by giving rise to an anti proportional characteristic. Considering exemplary the Pareto front of structure 4 (Figure 8**A**), which we present in Figure 8**C**, we observe that an increase in efficiency leads to a decrease of functionality and vice versa. In addition, we observe characteristic clustering at different levels of functional performance, likely being caused by different genetic gates at the later positions of the circuit (*18*). Despite this, the Pareto front presents a near convex behavior and allows for a smooth trade-off between function and energy. Our evaluations of exemplary structures for the Boolean function 0×26 featuring six genetic gates each also presents the anti-proportionality but does not exhibit the observed smoothness.

**Figure 8:**
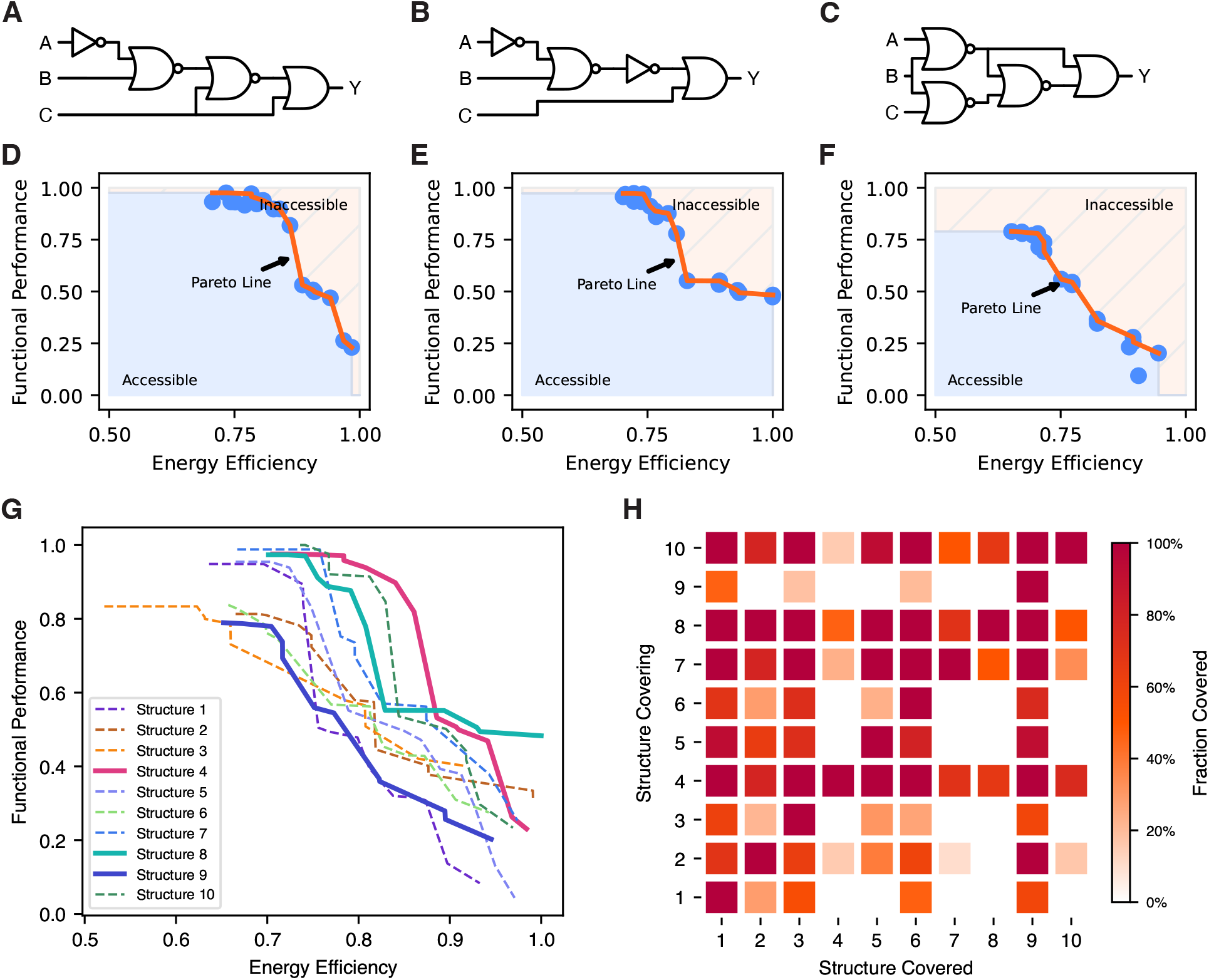
The Pareto front of the boolean function 0xF7. **A-C:** Three exemplary structures of the Boolean function 0xF7. In particular, Structure 4 in **A**, Structure 8 in **B** and Structure 9 in **C. D-F:** The Pareto fronts of the structures presented in **A-C**, where **D** presents the one of Structure 4, **E** the one of Structure 8, and **F** corresponds to Structure 9, respectively. Normalized to the same interval, the Pareto fronts (Figures **D-F**) exhibit different characteristics with the discontinuities and non-monotonicity resulting from discrete gate assignment and stochastic optimization. **D** combines high functional performance with moderate energy requirements while **E** allows the highest energy efficiency still preserving a moderate functional performance. The Pareto front of Structure 9 exhibits a near linear relationship between function and energy efficiency, with an inferior overall performance. **G:** We here illustrate the Pareto fronts of all the structures of 0xF7 jointly to emphasize comparability and the differences among them. Highlighting the Pareto fronts presented in Figures **D-F** by bold lines, their relationship to one another gets obvious. In addition, one observes that structure 9’s Pareto front is inferior to others. Figure **H** illustrates this quantitatively, by stating the portion of Pareto front covered by another Pareto front. Thereby, the rows indicate the Structure which’s Pareto front covers the respective Pareto front denoted in the column. This comparison emphasizes the dominating behavior of structures 4, 8 and 10 over the others with respect to both optimization goals.

Continuing with the Pareto fronts of structures 8 and 9 as visualized in Figures 8**E** and **F**, we observe that the general relationship persists while the exact form changes. In particular, the front in **E** appears inferior to the one in **D**, and this is surely the case for the areas with high functional performance, as structure 4 allows for higher energy efficiency there. However, when it comes close to the limits of energy efficiency, structure 8 (8**E**) is more robust in terms of the functional score achieved. Structure 9 (8**F**) features an almost linear transition region, but is obviously inferior to both of the previous with respect and to any optimization objective and the joint optimization as well.

Inspired by the inferior performance of structure 9 along the whole Pareto front, we compare the Pareto fronts of all structural variants of Boolean function 0xF7 in Figure 8**G**. The Pareto fronts are even more diverse than the ones already considered (highlighted by bold lines). Structures 4 and 8 cover almost all of the relevant optimization space, regardless of the trade-off considered. In contrast, seven out of ten structures do not allow the optimization to reach the best solution. Figure 8**H** visualizes this quantitatively by representing for each structure the portion of the Pareto points of any other structure covered. At first sight, one notices the four rows featuring a large number of dark squares. In particular, these are the rows of structures 4, 7, 8, and 10, which cover the Pareto fronts of the superior structures either completely or almost completely and thus exhibit a sort of Pareto dominance.

Despite considering energy efficiency and function jointly, the current optimizations only optimize either of the objectives while constraining the other. Multi-objective optimization provides a means to overcome this, as it allows for the joint optimization of both objectives (*58*). Dealing with the two objectives function and energy, represented by the scores *S* and *E*, one approach to realize multi-objective optimization is scalarization. In the case of scalarization, the objective functions are aggregated into a single objective, to which one can apply conventional optimization methods as the simulated annealing presented in (*30*).

Due to the well behaving shape of the Pareto fronts examined, we here suggest a linear scalarization, which is also known as the weighting method (*58, 59*). For it’s efficient application ahead of identifying the whole Pareto front, the scores need to be normalized (*58*). To this end, we introduce *S*_*r*_ and 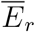 as reference maxima of the respective scores, which one obtains by an initial evaluation. In dependence to the weight parameter *ϕ* ∈ [0, 1], we express the scalarized performance of the genetic logic circuit (*γ, q*) by *V* (*γ, q*), which we define as

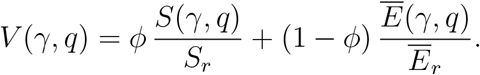

By the choice of the parameter *ϕ*, one implements the trade-off between the two objectives energy efficiency and function.

## 3 Conclusion and Outlook

In this work, we present an energy aware gene expression model characterizing the non-equilibrium steady state in terms of the first and second order moments and the associated energy requirements, which we relate to the model’s entropy production rate. This model can account for promoter architectures varying in the number of binding sites, activation steps, and cognate transcription factors.In contrast to the widely employed equilibrium models, the presented model captures non-equilibrium characteristics like increased sensitivity and sharpness, which are especially relevant in the context of eukaryotes. This includes the dissipation of energy, which is essential to life.In combination with the probabilistic NESS description and the presented relation between energy and entropy production rate, this allows for further evaluation of the trade-off between function and energy, beyond the scope of genetic logic circuits.

With this model at hand, we establish **Energy Aware Technology Mapping**, the design of genetic logic circuits with respect to function and energy efficiency. The *in silico* evaluation based on 33 Boolean circuits improves energy efficiency by 37.2% on average, but reduces the functionality to 28.5% compared to the functionally optimized variants. These improvements result from decreases in expression levels representing Boolean ON states, simultaneously decreasing the circuit’s functional performance. With respect to structural variants, we show that different structures of a single Boolean function vary in their energetic characteristic and can be beneficial for energy efficiency (up to 22.7% on average) and functionality. This also extends to the case of multi-objective optimization, as shown in the Pareto evaluation.

Based upon the insights gained in this study, one can further improve the design of genetic logic circuits with respect to energy by the creation of energy aware genetic gate libraries. This can include protein based gates focusing on robust functioning despite lower expression levels, the use of shorter transcription factors as signalling molecules, gates omitting translation by employing RNA based signals or any combination thereof.

To validate the theoretical framework of our proposed Energy Aware Technology Mapping for genetic logic circuits, we outline potential experimental evaluations. The simplest method to evaluate energy efficiency involves observing cell growth and viability by measuring optical density at 600 nm (OD600) and cell count (*60*). An alternative is the direct measurement of metabolites related to the energetic state of the cell *in vivo*, for which ATP sensors could be employed (*61, 62*). A more complex, yet insightful, method is RNA sequencing (RNA-seq). Sequencing the transcriptome of cells harboring synthetic circuits would not only reveal the transcripts associated with the circuits but also allows to check for upregulation and downregulation of genes linked to overall cell fitness (*63*). As this approach addresses the increased metabolic burden that larger circuits impose, incorporating energy dissipation while maintaining circuit functionality could enable the development of larger circuits that operate effectively *in vivo*.

In conclusion, our results provide strong evidence for the exclusivity of the objectives functionality and energy efficiency in the context of genetic logic circuits. This matches previous results on the relation between energy expenditure and precision and sharpness. Additionally, the consideration of structural variants proves relevant also in the case of energy efficiency. Since it is beneficial to consider structural variants regardless of the optimization goal, disregarding them is likely to lead to sub-optimal solutions in terms of both, energy efficiency and functionality. Given the proportionality between energy consumption and the rate of heterologous protein expression, optimization of energy can effectively reduce the expression of heterologous proteins, thereby allowing for energy efficiency by design. This is particularly relevant in the context of metabolic burden, as it allows cellular fitness to be maintained by reducing the amount of resources diverted from the host by synthetic constructs. In that sense, the well being of the host organism ensures the desired functionality of the genetic logic circuit.The energetic advantage of one to two gates provided by Energy Aware Technology Mapping can be crucial for implementing the circuit *in vivo* in a resource constrained host organism.

## 4 Methods

The here presented methods present the techniques most relevant to our work. Within the referenced supplementary information, this is complemented.

### 4.1 Gene Circuit Simulation

The input to the gene circuit are the concentrations of the inducer molecules. While *in vivo* the inducer binds to constitutively expressed repressive transcription factors, we realize the dependency on the inducer levels by a lookup table based approach, matching a degenerate promoter model with the desired promoter activity to the provided inducer concentration. For the simulation of the gene circuit, we apply our energy aware gene expression model one after another to the genes within the circuit. Thereby, one has to preserve topological order, meaning that a gene can only be considered when all genes expressing the cognate transcription factor have already been evaluated. To obtain the concentration of the protein encoded by a gene, we apply a log-normal closure to the moments provided by Equations (3) and (4) and draw a sample. In case multiple genes express the same protein, the protein’s final concentration is the superposition of the single genes’ contributions. By performing this approach many times in parallel, one obtains an empirical distribution predicting the population dynamics of cells implementing the particular genetic logic circuit. While this is a sampling based approach, one can instead propagate the mean or the median as the protein abundance to yield a point estimate. For a more formal treatment of probabilistic genetic logic circuit simulation, we refer the interested reader to (*30*).

### 4.2 Steady State Distribution of CTMCs and Kirchhoff’s Theorem

The unique steady state distribution of an ergodic CTMC with *n* states can be derived by using its propensity matrix **Λ**, defined in row sum zero form. The first step is to identify the null-space, respectively solving for the left eigenvector ***v*** = [*v*_1_ … *v*_*n*_] corresponding to the eigenvalue 0 of **Λ**.

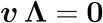

The steady state distribution ***π*** is then defined as

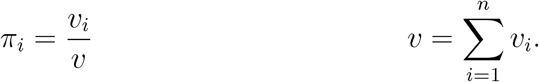

Schnakenberg (*53*) and later (*64*) describe an alternative approach, called Kirchhoff’s theorem -as initially described by Kirchhoff-based on the enumeration of spanning trees.

The subsequent contains a rather informal description, wherefore we refer the interested reader to (*53*) and (*64*).

Before we start with the actual method, we recall the definition of a spanning tree first. A spanning tree of an undirected graph is a tree and a subgraph, containing all the vertices of the graph. The undirected graph being a prerequisite, we derive it by identifying the vertices of the graph with the states in the state space of our CTMC. Next, the forward and backward reactions between two states are aggregated into a single undirected edge, connecting the vertices associated to the states.

We continue with enumerating all possible spanning trees of the graph. In total, there are *n*_*t*_ trees and we refer to the *j*th with *T*_*j*_. Next, we introduce the function Π_*i*_(*T*_*j*_), which maps the spanning tree *T*_*j*_ to the subgraph of the CTMC’s state space that includes the reactions being part of the spanning tree and directed to state *z*_*i*_. In particular, this is done by setting *z*_*i*_ as the root node and successively traversing the edges in the spanning tree *T*_*j*_. Actually, each edge corresponds to two reactions, where the one pointing in the direction of the root node is preserved and the other is dropped. The result is a directed tree, consisting of the reactions that lead from any initial state to the root state *z*_*i*_. With the previous, we assign the value

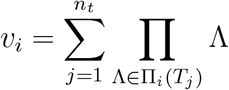

to each state *z*_*i*_, where the product runs over the propensities of the reactions in Π_*i*_(*T*_*j*_). As before the steady state distribution is given by

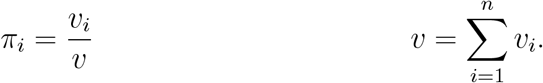

While in theory this method is not limited by the size of the CTMC’s state space, the practical applicability depends on the complexity of the obtained graph’s topology (*53*). However, this approach provides an appealing way to derive the steady state distribution symbolically. As such, our implementation makes use of the steady state distribution derived from Kirchhoff’s theorem for the promoter architectures considered, while it also provides the simpler eigenvector approach for other architectures.

### 4.3 Parameter Estimation

The estimation of a parameter set matching the characteristics of the part to capture is essential for the model’s expressiveness within a technology mapping application. As our model gives rise to mean and variance of the response characteristic, data sources giving rise to both are ideal. One possible source is cytometry data for different inducer concentrations. As the histograms provide rich information on the distribution, the mean and variance can be derived by evaluating

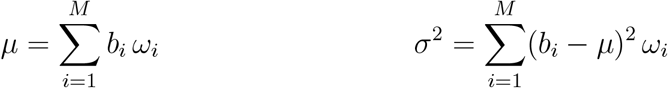

where (*b*_1_, …, *b*_*M*_) are the bin centers and (*ω*_1_, …, *ω*_*M*_) are the frequencies in a histogram with *M* bins.

As model instantiation, we use a promoter architecture featuring three binding sites for a single cognate transcription factor and having only two activation steps with respect to transcriptional activity as depicted in Figure 2**AB**. Since we collapse states with equal number of transcription factors bound into a single state, we obtain a CTMC of eight states and 20 reactions transitioning between these states. In addition, we constrain all the states corresponding to the same transcriptional activation level to have the same promoter activity *a*_*i*_. By fixing the reaction rate constants for RNA and protein dynamics are fixed in dependence to the organism (see SI Section S7), our model instantiation features 22 parameters, 20 for the reaction rate constants of the promoter’s CTMC and two for the two promoter activity levels. From an intuitive perspective, the rate constants between two states of differing promoter activity levels balance the output dynamic maximally achievable. The reactions depending on the transcription factor abundance and their reverse reactions then determine the location of the transition. In foregoing evaluations, this instantiation features a well balance between the complexity of the model and the capabilities of modeling response characteristics of interest, especially the sharpness encountered. However, the variance in the data is not completely explained by this, likely caused by extrinsic noise (*49–51*) not accounted for in this model.

For parameter estimation, we apply gradient based optimization and minimize the logarithmic difference between the model’s prediction and the reference data from (*40*). We constrain the reaction rate constants to the range [10^−5^, 10^5^], penalize non monotonic response characteristics, and by weighting the error prioritize model quality in the saturated regions. In this context, it is important to note that the parameter estimation considers mean and variance. The mean dynamics itself are insufficient to uniquely determine a set of reaction rate constants as outlined at the end of SI Section S2.2. For a detailed description of the parameter estimation process please refer to SI Section S5.2.

### 4.4 ARCTIC

ARCTIC is the technology mapping framework for genetic logic circuits used in this work. It takes a combinational Boolean specification as input and constructs all possible structural variants for this specification based on a given library of gates. It then uses different scoring methods to optimize the assignment of library gates to the elements of the topologies. In this way it can search for the best performing genetic circuit for a given Boolean specification.

ARCTIC offers different circuit models and scores that focus on robust genetic circuits. It implements different optimization methods to leverage these models and to explore the design space. In this work, the Simulated Annealing heuristic has been used. It applies a neighborhood structure leveraging functional proximity of gates and is flexible in terms of the used design objective.

We employed deterministic and sampling based simulations. The evaluation on the benchmark set as well as the structural variants made use of the deterministic approach taking the average into account. The constraint optimization used a Cello (*11*) or E-score (*30*) of 200 as lower bound on the functionality, which corresponds to a 200 fold-change of the median values. This bound is sufficient for circuit functionality and, in combination with a sampling based approach and the E-score, still allows for robust circuit designs. During optimization for circuit function, the upper bound on the energy expenditure was set in dependence to the circuits size, allowing for 20.000 *k*_B_ *T s*^−1^ per gate. In the Pareto evaluation, we employ the sampling based approach with the E-score and consider 1000 samples for each output distribution.

#### E-score

The E-score (*30*) is a generalization of Cello’s circuit score (*11*) to the case where circuit outputs are random variables. This takes stochastic effects in populations of cellular hosts into account. Intuitively, the more “distance” lies between distributions that represent ON states and those that represent OFF states, the higher a circuit is scored. The E-score is calculated as the exponential of a modified 1-Wasserstein distance between empirical distributions of the logarithms of the concentrations or copy-numbers of the output chemical species. Consider first two sets of *N* samples <u>*y*</u> ≡ {<u>*y*</u>^(1)^, <u>*y*</u>^(2)^, …, <u>*y*</u>^(*N*)^} and 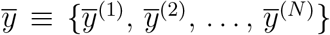 that represent two empirical circuit output distributions. The set *y* consists of those samples corresponding to an anticipated circuit output of logical zero (OFF) and 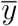 consists of those samples corresponding to an anticipated circuit output of logical one (ON). We then calculate the modified 1-Wasserstein distance 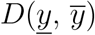 on the samples’ logarithms by

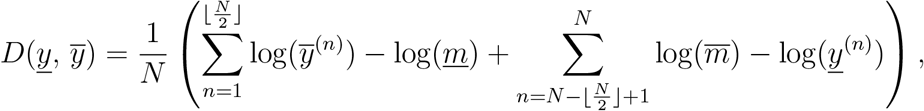

where *m* and 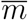 are the medians of *y* and 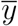 respectively. Consider now having several sample sets <u>*Y*</u> ≡ <u>*Y*</u> (*γ, q*) = {<u>*y*</u>_1_, <u>*y*</u>_2_, …, <u>*y*</u>_*K*_ } representing logical OFF states and 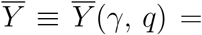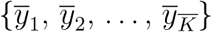 logical ON states, where each individual sample set corresponds to one of 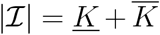 different Boolean input conditions of a circuit with structure *γ* and assignment *q*. The circuit’s E-score is then obtained by

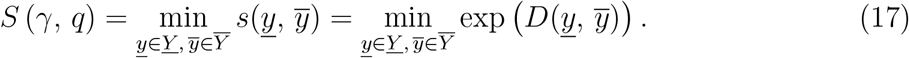

It is shown in (*30*) that *S* (*γ, q*) simplifies to Cello’s circuit score for *γ* and *q* if *N* = 1 and the single samples correspond to Cello’s approximations for the median. Note, that the exponentiation in Equation (17) can be applied after taking the minimum to obtain the score *S*.

## 5 Code Availability

The model is implemented as part of the technology mapping framework ARCTIC, which is available at https://www.rs.tu-darmstadt.de/ARCTIC.

## 6 Associated Content

### 6.1 Supplementary Information

Derivations of moment equations, the chemical reaction network of the model presented, background on chemical reaction networks, further methods and extended figures and tables.

## 7 Author Contributions

H.K., C.H., and H.G. provided the research idea and contributed methodology. E.K. derived the model, implemented the model and the Energy Aware Technology Mapping in ARCTIC, and designed and did evaluation of Energy Aware Technology Mapping. M.G. and E.K. performed mathematical analysis of stochastic thermodynamics. T.S. provided the interface for ARCTIC and executed evaluation. J.M. and M.M. contributed to the biological background.

N.E. contributed to ARCTIC and the E-score. All authors contributed to the writing of the paper.

## 8 Acknowledgments

The authors acknowledge all people in the Self-Organizing Systems lab for the valuable discussions. We especially thank the anonymous reviewers for their valuable comments and suggestions and Eike Mentzendorff for advice on multi-objective optimization. For the use of SBOLCanvas, we acknowledge Terry et al. (*65*), Quinn et al. (*32*), and the SBOL community (*31*). Funded by the European Union. Views and opinions expressed are however those of the author(s) only and do not necessarily reflect those of the European Union or the European Research Council Executive Agency. Neither the European Union nor the granting authority can be held responsible for them. E.K. was supported by ERC-PoC grant PLATE (101082333). J.M. was supported by the MSCA-DN SYNSENSO (GA 101072980). N.E. acknowledges funding by the German Research Foundation (DFG) as part of the project B4 within the Collaborative Research Center (CRC) 1053 – MAKI. H.G.G. was supported by NIH R01 Awards R01GM139913 and 1R01GM152815, by the Koret-UC Berkeley-Tel Aviv University Initiative in Computational Biology and Bioinformatics, and by a Winkler Scholar Faculty Award. H.G.G. is also a Chan Zuckerberg Biohub Investigator (Biohub – San Francisco).

