## Supplementary Information for "Energy Aware Technology Mapping of Genetic Logic Circuits"

### Contents


**S1 The Model's Chemical Master Equation** **S4**

S1.1 Chemical Reaction Networks . . . S4

S1.2 The Model's Chemical Reaction Network Representation . . . S5

S1.3 The Model's Chemical Master Equation . . . S7

S1.4 The Promoter's Marginal Distribution . . . S9

**S2 The Model's Moment Equations** **S11**

S2.1 Chapman Kolmogorov Backward Equation . . . S11

S2.2 Steady State Promoter Dynamics . . . S12

S2.3 Steady State RNA Dynamics . . . S17

S2.4 Steady State Protein Dynamics . . . S21

**S3 Stochastic Thermodynamics of Open Chemical Reaction Networks** **S25**

S3.1 Closed Stochastic Chemical Reaction Networks . . . S26

S3.2 Open Chemical Reaction Networks – Coupling to Chemostats . . . S29

S3.3 Relation of Entropy Production Rate and Energy Dissipation . . . S34

S3.4 Energy Dissipation of RNA and Protein Dynamics . . . S36

S3.5 Entropy Production Rate on a Finite State Space . . . S38

**S4 Schnakenberg's Cycle Representation of the Entropy Production Rate** **S39**

S4.1 Entropy Production Rate of the Eight State Promoter . . . S40

**S5 ARCTIC and Energy Aware Technology Mapping** **S44**

S5.1 Library Creation . . . S46

S5.2 Parameter Estimation . . . S49

**S6 Relative Promoter Units, Relative Expression Units, and the Characterization of Genetic Logic Gates in Cello** **S50**

**S7 Gene Expression Kinetics in *S. cerevisiae*** **S51**

S7.1 Transcription Rates and Energy Dependence . . . . . . . . . . . . . . . . . . . . . . S51

S7.2 Translation Rates and Energy Dependence . . . . . . . . . . . . . . . . . . . . . . . S52

S7.3 Degradation and Dilution Rates . . . . . . . . . . . . . . . . . . . . . . . . . . . . . S53

**S8 Extended Figures and Tables** **S55**

**References** **S62**

### S1 The Model's Chemical Master Equation

In this section, we state the chemical master equation presented in the main paper. In addition, we provide some background on chemical reaction networks and show how our system can be modelled by these.

$$
\begin{aligned}
\frac{d}{dt}\,\mathrm{p}(z_i, r, p \mid \boldsymbol{c}, t) = &\sum_{j=1, j\neq i}^{n} \Lambda_{ji}\, \mathrm{p}(z_i, r, p \mid \boldsymbol{c}, t) - \sum_{j=1, j\neq i}^{n} \Lambda_{ij}\, \mathrm{p}(z_i, r, p \mid \boldsymbol{c}, t) \\
&+ \mu_1\, a_i\, \mathrm{p}(z_i, r-1, p \mid \boldsymbol{c}, t) - \mu_1\, a_i\, \mathrm{p}(z_i, r, p \mid \boldsymbol{c}, t) \\
&+ \delta_1\, (r+1)\, \mathrm{p}(z_i, r+1, p \mid \boldsymbol{c}, t) - \delta_1\, r\, \mathrm{p}(z_i, r, p \mid \boldsymbol{c}, t) \\
&+ \mu_2\, r\, \mathrm{p}(z_i, r, p-1 \mid \boldsymbol{c}, t) - \mu_2\, r\, \mathrm{p}(z_i, r, p \mid \boldsymbol{c}, t) \\
&+ \delta_2\, (p+1)\, \mathrm{p}(z_i, r, p+1 \mid \boldsymbol{c}, t) - \delta_2\, p\, \mathrm{p}(z_i, r, p \mid \boldsymbol{c}, t)
\end{aligned}
$$

#### S1.1 Chemical Reaction Networks

Within this work, we oftenly consider chemical reaction networks. Formally, they are defined by the set $\mathcal{R} = \{\mathcal{R}_1, \ldots, \mathcal{R}_M\}$ of chemical reactions on the $N \in \mathbb{N}$ chemical species $\mathsf{\mathbf{X}} = (\mathsf{X}_1, \ldots, \mathsf{X}_N)$. The single reactions are defined as

$$
\mathsf{R}_m : \quad \sum_{n=1}^{N} a_n^{(m)} \mathsf{X}_n \quad \xrightarrow{k_m} \quad \sum_{n=1}^{N} b_n^{(m)} \mathsf{X}_n
$$

where $k_m$ is the reaction rate constant and $a_n^{(m)}, b_n^{(m)} \in \mathbb{N}$ are the substrate and product coefficients for species $\mathsf{X}_n$ in reaction $\mathcal{R}_m$, which are subsumed in the matrices $\boldsymbol{A}$ and $\boldsymbol{B}$, with $\boldsymbol{A}, \boldsymbol{B} \in \mathbb{N}^{M\times N}$. The stoichiometric matrix $\overline{\boldsymbol{N}} \in \mathbb{Z}^{M\times N}$ of the system is given by $\overline{\boldsymbol{N}} = \boldsymbol{B} - \boldsymbol{A}$. The stoichiometric change vectors $\nu_m$ follow directly as the column vectors of $\overline{\boldsymbol{N}}$ and give rise to the change in the species vector $\mathsf{\mathbf{X}}$ induced by reaction $\mathcal{R}_m$.

Not yet specified further is the rate of the reaction. Within this work, we assume the rates to follow mass action kinetics (1–3) and describe the resulting rate by the propensity function $\lambda_m(\boldsymbol{X}) \in \mathbb{R}_{\geq 0}$, where $\boldsymbol{X}$ is the vector of molecule counts associated to the species in $\mathsf{\mathbf{X}}$.

#### S1.2 The Model's Chemical Reaction Network Representation

We can subdivide the presented model into three parts. The promoter, the RNA dynamics and the protein dynamics. To allow for a convenient treatment later on, we here present how to capture the model and especially the promoter in terms of a chemical reaction network.

Beginning with the promoter, we represent by $Z$ the promoter's state. At a single point in time, this variable can take a single value of $z_1, \ldots, z_n$ for an $n$ state promoter. In addition, only transitions starting at the current state (e.g. $z_i$) are allowed. We can realize this conveniently by representing the promoter's state by a one-hot encoding, where we introduce for each state $z_i$ a chemical species $\mathsf{Z}_i$ and enforce $\sum_{i=1}^{n} Z_i = 1$ for the species' molecule counts. The current state of the promoter is then the species vector $\boldsymbol{Z} = (Z_1, \ldots, Z_n)$.

By representing the state in terms of a one-hot encoding, we can make use of the law of mass action to describe the rates of the reactions. In particular, the propensities arise naturally from our definition as

$$\Lambda_{ij}(\boldsymbol{Z}) = \Lambda_{ij} = Z_i \, \tilde{k}_{ij} \tag{S1}$$

with $Z_i \in \{0, 1\}$ and $\tilde{k}_{ij}$ being a pseudo reaction rate constant defined as $\tilde{k}_{ij} = f(\boldsymbol{c})\, k_{ij}$, where $k_{ij}$ is the actual reaction rate constant and $f(\boldsymbol{c})$ a monomial function realizing the dependency on the transcription factors of relevance.

Consequently, this interpretation has to propagate to the distribution characterizing our system. Practically, $\pi_i$ is still the probability of finding the system in state $z_i$ in the steady state. However, now this gives rise to the probability of $Z_i = 1$ instead of $Z = z_i$.

For the RNA and protein dynamics, the formulation is straight forward and follows directly from the model definition.

$$
\begin{aligned}
\text{I}: &\qquad \mathsf{Z}_i \xrightarrow{\mu_1\, a_i} \mathsf{Z}_i + RNA \qquad i = 1,\ldots,n\\
\text{II}: &\qquad RNA \xrightarrow{\mu_2} RNA + protein\\
\text{III}: &\qquad RNA \xrightarrow{\delta_1} \varnothing\\
\text{IV}: &\qquad protein \xrightarrow{\delta_2} \varnothing
\end{aligned}
$$

By introducing $\mathsf{Z}_i$ as substrate species for the transcription, we implicitly add the dependency on this species for the respective transcription rate via the law of mass action, while the corresponding transcriptional activity is indicated by $a_i$. In particular, the reaction I exists $n$ times, such that the instantaneous rate of RNA production is

$$
\begin{aligned}
\sum_{j=1}^{n} \mu_1\, a_j\, Z_j &= \mu_1 \sum_{j=1}^{n} a_j\, Z_j\\
&= \mu_1\, a_i\, Z_i + \mu_1 \sum_{j=1, j\neq i}^{n} a_j\, Z_j\\
&= \mu_1\, a_i
\end{aligned}
\tag{S2}
$$

which is the desired rate of RNA production in state $z_i$. For the transition to the last step, we exploit that $Z_i = 1$ implies $Z_j = 0$ for all $j \neq i$, which is a direct consequence of the one-hot encoding. In the later, we use $R$ and $P$ to the abundance of RNA and protein.

**Stoichiometric Change Vectors and Propensities** We here extend on the stoichiometric change vectors $\boldsymbol{\nu}$ and the propensities of our CTMC. For the promoter, the stoichiometric change vector realizes the transition from state $z_i$ to state $z_j$. Following the one-hot encoding, a state $z_i$ is represented by $\boldsymbol{e}_i$, wich is the $i$th unit vector with all zeros except a single one at position $i$. Consequently, the stoichiometric change vector associated to the transition

$z_i$ to $z_j$ and acting on the promoter state representation $\boldsymbol{Z}$ is given by

$$\boldsymbol{\nu}_{ij} = \boldsymbol{e}_j - \boldsymbol{e}_i \tag{S3}$$

and extends to the overall state vector $\boldsymbol{X}$ by adding zeros at the positions of RNA and protein. For the RNA and protein dynamics, the change vectors for the state vector $\begin{bmatrix} R & P \end{bmatrix}^T$ are

$$\nu_{\mathrm{I}} = \begin{bmatrix} 1 \\ 0 \end{bmatrix} \qquad \nu_{\mathrm{II}} = \begin{bmatrix} 0 \\ 1 \end{bmatrix} \qquad \nu_{\mathrm{III}} = \begin{bmatrix} -1 \\ 0 \end{bmatrix} \qquad \nu_{\mathrm{IV}} = \begin{bmatrix} 0 \\ -1 \end{bmatrix}.$$

As before, they extend to $\boldsymbol{X}$ by adding zeros at the positions of the promoter species.

Left over are the propensities of the reactions. For the promoter, the propensities are given by Equation (S1) where the dependency on external transcription factor concentrations is included in $\tilde{k}_{ij}$. For the RNA and protein dynamics, the propensities are

$$\lambda_{\mathrm{I}}(\boldsymbol{X}) = Z_i\,\mu_1\,a_i \qquad \lambda_{\mathrm{II}}(\boldsymbol{X}) = R\,\mu_2 \qquad \lambda_{\mathrm{III}}(\boldsymbol{X}) = R\,\delta_1 \qquad \lambda_{\mathrm{IV}}(\boldsymbol{X}) = P\,\delta_2$$

with $R, P$ as the current RNA and protein abundance levels. As shown previously, the overall instantaneous RNA production rate is $\mu_1\,a_i$ for the promoter being in state $z_i$ as given by Equation (S2).

#### S1.3 The Model's Chemical Master Equation

On the basis of the chemical reaction network presented previously, we can derive the corresponding chemical master equation. This master equation gives rise to the time evolution of the system's probability distribution. In particular, it describes the in and outflux of probability mass into a distinct state of the system. Formally, the chemical master equation for a chemical reaction network as defined previously is given by

$$\frac{d}{dt}\mathrm{P}[\boldsymbol{X}(t)=\boldsymbol{x}] = \sum_{j=1}^{M}\lambda_j(\boldsymbol{x}-\boldsymbol{\nu}_j)\ \mathrm{P}[\boldsymbol{X}(t)=\boldsymbol{x}-\boldsymbol{\nu}_j] - \sum_{j=1}^{M}\lambda_j(\boldsymbol{x})\ \mathrm{P}[\boldsymbol{X}(t)=\boldsymbol{x}]$$

where the sum goes over all reactions and $\boldsymbol{X}(t)$ is the vector of species, explicitly including the time dependency.

In the context of our model, with $\boldsymbol{Z}$ being the random variable vector for the promoter state in one-hot encoding and $R$ and $P$ being the random variables for RNA and protein, we get $\boldsymbol{X}(t) = (\boldsymbol{Z}(t), R(t), P(t))$, while the corresponding probability reads $\mathrm{P}[\boldsymbol{Z}(t) = \boldsymbol{e}_i, R(t) = r, P(t) = p \mid \boldsymbol{c}]$, with $\boldsymbol{c}$ being the vector of cognate transcription factor concentrations and $\boldsymbol{e}_i$ denoting the $i$th unit vector. For reasons of brevity, we use $\mathrm{p}(\boldsymbol{e}_i, r, p \mid \boldsymbol{c}, t) = \mathrm{P}[\boldsymbol{Z}(t) = \boldsymbol{e}_i, R(t) = r, P(t) = p \mid \boldsymbol{c}]$ and obtain the chemical master equation

$$\begin{aligned}\frac{d}{dt}\mathrm{p}(\boldsymbol{e}_i, r, p \mid \boldsymbol{c}, t) = &\sum_{j=1, j\neq i}^{n}\Lambda_{ji}\ \mathrm{p}(\boldsymbol{e}_j, r, p \mid \boldsymbol{c}, t) - \sum_{j=1, j\neq i}^{n}\Lambda_{ij}\ \mathrm{p}(\boldsymbol{e}_i, r, p \mid \boldsymbol{c}, t)\\ &+ \mu_1\, a_i\ \mathrm{p}(\boldsymbol{e}_i, r-1, p \mid \boldsymbol{c}, t) - \mu_1\, a_i\ \mathrm{p}(\boldsymbol{e}_i, r, p \mid \boldsymbol{c}, t)\\ &+ \delta_1\,(r+1)\ \mathrm{p}(\boldsymbol{e}_i, r+1, p \mid \boldsymbol{c}, t) - \delta_1\, r\ \mathrm{p}(\boldsymbol{e}_i, r, p \mid \boldsymbol{c}, t)\\ &+ \mu_2\, r\ \mathrm{p}(\boldsymbol{e}_i, r, p-1 \mid \boldsymbol{c}, t) - \mu_2\, r\ \mathrm{p}(\boldsymbol{e}_i, r, p \mid \boldsymbol{c}, t)\\ &+ \delta_2\,(p+1)\ \mathrm{p}(\boldsymbol{e}_i, r, p+1 \mid \boldsymbol{c}, t) - \delta_2\, p\ \mathrm{p}(\boldsymbol{e}_i, r, p \mid \boldsymbol{c}, t)\end{aligned}$$

where the first line corresponds to the promoter dynamics governed by the propensities $\Lambda_{ij}$, the second and third row describe the RNA and the fourth and fifth row the protein dynamics. Reidentifying the one-hot encoding of the promoter with the states $z_1, \ldots, z_n$ and the state variable $Z$, we get

$$\mathrm{P}[Z(t) = z_i, R(t) = r, P(t) = p \mid \boldsymbol{c}] = \mathrm{P}[\boldsymbol{Z}(t) = \boldsymbol{e}_i, R(t) = r, P(t) = p \mid \boldsymbol{c}].$$

Consequently, the desired chemical master equation presented in the main part follows by using $\mathrm{p}(z_i, r, p \mid \boldsymbol{c}, t) = \mathrm{P}[Z(t) = z_i, R(t) = r, P(t) = p \mid \boldsymbol{c}]$.

$$\begin{aligned}\frac{d}{dt}\mathrm{p}(z_i, r, p \mid \boldsymbol{c}, t) = &\sum_{j=1, j\neq i}^{n} \Lambda_{ji}\, \mathrm{p}(z_i, r, p \mid \boldsymbol{c}, t) - \sum_{j=1, j\neq i}^{n} \Lambda_{ij}\, \mathrm{p}(z_i, r, p \mid \boldsymbol{c}, t)\\ &+ \mu_1\, a_i\, \mathrm{p}(z_i, r-1, p \mid \boldsymbol{c}, t) - \mu_1\, a_i\, \mathrm{p}(z_i, r, p \mid \boldsymbol{c}, t)\\ &+ \delta_1\,(r+1)\, \mathrm{p}(z_i, r+1, p \mid \boldsymbol{c}, t) - \delta_1\, r\, \mathrm{p}(z_i, r, p \mid \boldsymbol{c}, t)\\ &+ \mu_2\, r\, \mathrm{p}(z_i, r, p-1 \mid \boldsymbol{c}, t) - \mu_2\, r\, \mathrm{p}(z_i, r, p \mid \boldsymbol{c}, t)\\ &+ \delta_2\,(p+1)\, \mathrm{p}(z_i, r, p+1 \mid \boldsymbol{c}, t) - \delta_2\, p\, \mathrm{p}(z_i, r, p \mid \boldsymbol{c}, t)\end{aligned} \tag{S4}$$

#### S1.4 The Promoter's Marginal Distribution

We here show, that the distribution of the promoter can be directly derived from the promoter's propensity matrix. To this end, we start with the definition of the promoters marginal distribution.

$$\mathrm{P}[Z(t) = z_i \mid \boldsymbol{c}] = \sum_{r,p=0}^{\infty} \mathrm{P}[Z(t) = z_i, R(t) = r, P(t) = p \mid \boldsymbol{c}]$$

Making use of the compressed notation $\mathrm{p}(z_i, r, p \mid \boldsymbol{c}, t)$ again and taking the derivative of both sides yields

$$\begin{aligned}\frac{d}{dt}\mathrm{p}(z_i \mid \boldsymbol{c}, t) &= \frac{d}{dt}\sum_{r,p=0}^{\infty} \mathrm{p}(z_i, r, p \mid \boldsymbol{c}, t)\\ &= \sum_{r,p=0}^{\infty} \frac{d}{dt}\mathrm{p}(z_i, r, p \mid \boldsymbol{c}, t).\end{aligned}$$

Inserting the chemical master equation (Equation (S4)) and rearranging the sums a bit gives

$$\begin{aligned}
\frac{d}{dt}\,\mathrm{p}(z_i \mid \boldsymbol{c}, t) = &\sum_{j=1, j\neq i}^{n} \Lambda_{ji} \sum_{r,p=0}^{\infty} \mathrm{p}(z_i, r, p \mid \boldsymbol{c}, t) - \sum_{j=1, j\neq i}^{n} \Lambda_{ij} \sum_{r,p=0}^{\infty} \mathrm{p}(z_i, r, p \mid \boldsymbol{c}, t) \\
&+ \mu_1\, a_i \sum_{r,p=0}^{\infty} \mathrm{p}(z_i, r-1, p \mid \boldsymbol{c}, t) - \mu_1\, a_i \sum_{r,p=0}^{\infty} \mathrm{p}(z_i, r, p \mid \boldsymbol{c}, t) \\
&+ \delta_1 \sum_{r,p=0}^{\infty} (r+1)\; \mathrm{p}(z_i, r+1, p \mid \boldsymbol{c}, t) - \delta_1 \sum_{r,p=0}^{\infty} r\; \mathrm{p}(z_i, r, p \mid \boldsymbol{c}, t) \\
&+ \mu_2 \sum_{r,p=0}^{\infty} r\; \mathrm{p}(z_i, r, p-1 \mid \boldsymbol{c}, t) - \mu_2 \sum_{r,p=0}^{\infty} r\; \mathrm{p}(z_i, r, p \mid \boldsymbol{c}, t) \\
&+ \delta_2 \sum_{r,p=0}^{\infty} (p+1)\; \mathrm{p}(z_i, r, p+1 \mid \boldsymbol{c}, t) - \delta_2 \sum_{r,p=0}^{\infty} p\; \mathrm{p}(z_i, r, p \mid \boldsymbol{c}, t)
\end{aligned}$$

which we can simplify further by defining $\mathrm{p}(z_i, r, p \mid \boldsymbol{c}, t) = 0$ for $r, p < 0$ giving rise to the following identities:

$$\begin{aligned}
\sum_{r=0}^{\infty} \mathrm{p}(z_i, r-1, p \mid \boldsymbol{c}, t) &= \mathrm{p}(z_i, -1, p \mid \boldsymbol{c}, t) + \sum_{r=1}^{\infty} \mathrm{p}(z_i, r-1, p \mid \boldsymbol{c}, t) \\
&= \sum_{r=0}^{\infty} \mathrm{p}(z_i, r, p \mid \boldsymbol{c}, t) \\
\sum_{p=0}^{\infty} \mathrm{p}(z_i, r, p-1 \mid \boldsymbol{c}, t) &= \sum_{p=0}^{\infty} \mathrm{p}(z_i, r, p \mid \boldsymbol{c}, t) \\
\sum_{r=0}^{\infty} (r+1)\, \mathrm{p}(z_i, r+1, p \mid \boldsymbol{c}, t) &= \sum_{r=1}^{\infty} r\, \mathrm{p}(z_i, r, p \mid \boldsymbol{c}, t) \\
&= \sum_{r=0}^{\infty} r\, \mathrm{p}(z_i, r, p \mid \boldsymbol{c}, t) \\
\sum_{p=0}^{\infty} (p+1)\, \mathrm{p}(z_i, r, p+1 \mid \boldsymbol{c}, t) &= \sum_{p=0}^{\infty} p\, \mathrm{p}(z_i, r, p \mid \boldsymbol{c}, t)
\end{aligned}$$

Making use of promoter marginal distribution definition and reshaping after inserting the previous identities, we obtain

$$\frac{d}{dt}\,\mathrm{p}(z_i \mid \boldsymbol{c}, t) = \sum_{j=1, j\neq i}^{n} \Lambda_{ji}\; \mathrm{p}(z_i \mid \boldsymbol{c}, t) - \sum_{j=1, j\neq i}^{n} \Lambda_{ij}\; \mathrm{p}(z_i \mid \boldsymbol{c}, t)$$

which is the intended result. In particular it states, that the probability evolution of the promoter only depends on the propensities given by the $\Lambda_{ij}$. Because of this, it is applicable to derive the steady state distribution of the promoter following the descriptions presented in the methods of this work.

### S2 The Model's Moment Equations

On the basis of the chemical reaction network and the chemical master equation introduced previously, we derive in this section the moment equations for RNA and protein characterizing our model. In particular, these are the average promoter activity

$$\alpha(\boldsymbol{c}) = \sum_{i=1}^{n} a_i \, \pi_i.$$

and the protein's mean and variance and the RNA's mean

$$\mathsf{E}[P \mid \boldsymbol{c}] = \frac{\mu_2}{\delta_2} \, \mathsf{E}[R \mid \boldsymbol{c}] \qquad \mathsf{E}[R \mid \boldsymbol{c}] = \frac{\mu_1}{\delta_1} \, \alpha(\boldsymbol{c})$$

$$\begin{aligned} \mathsf{Var}[P \mid \boldsymbol{c}] = \frac{\mu}{\delta} \, \alpha(\boldsymbol{c}) \left(1 - \frac{\mu}{\delta} \, \alpha(\boldsymbol{c})\right) + \frac{\mu_2}{\delta_2 + \delta_1} \, \frac{\mu}{\delta} \, \alpha(\boldsymbol{c}) \\ + \frac{\mu^2}{\delta + \delta_2^2} \, \boldsymbol{a} \left(\delta_1^{-1} + \boldsymbol{M}_2^{-1}\right) \boldsymbol{M}_1^{-1} \left(\boldsymbol{a} \odot \boldsymbol{\pi}(\boldsymbol{c})\right) \end{aligned}$$

as presented in the main part of this work.

#### S2.1 Chapman Kolmogorov Backward Equation

The general Chapman Kolmogorov backward equation is given by

$$\frac{d}{dt} \mathsf{E}[g(\mathbf{X}(t))] = \sum_{m=1}^{M} \mathsf{E}[\lambda_m(\mathbf{X}(t)) \left(g(\mathbf{X}(t) + \boldsymbol{\nu}_m) - g(\mathbf{X}(t))\right)]$$

and provides a formal method to derive the moments, here given by the monomial function $g(\cdot)$, of a system modelled by a CTMC. For example, $g(\mathbf{X}(t)) = P$ returns the mean dynamics of protein, where we drop the explicit dependency on $t$ for reasons of brevity. As before, $\lambda_j(\cdot)$ is the propensity function of the $j$th reaction and $\nu_j$ is the associated change vector.

In the steady state, the previous simplifies to

$$\sum_{m=1}^{M} \mathsf{E}[\lambda_m(\mathbf{X}(t))\,(g(\mathbf{X}(t)+\boldsymbol{\nu}_m) - g(\mathbf{X}(t)))] = 0$$
$$\sum_{m=1}^{M} \mathsf{E}[\lambda_m(\mathbf{X}(t))\,g(\mathbf{X}(t)+\boldsymbol{\nu}_m)] = \sum_{m=1}^{M} \mathsf{E}[\lambda_m(\mathbf{X}(t))\,g(\mathbf{X}(t))]$$

which we will use to derive the first and second order moments of the RNA and protein distribution. A fortunate aspect of the chemical reaction network under consideration is, that all propensities are zero or first order and so linear. This yields closed equations for all moments and allows to derive the moments of interest exactly. In systems with second or higher order reactions, moment equations would depend on higher order moments, which need to be approximated by a moment closure technique.

#### S2.2 Steady State Promoter Dynamics

The transcriptional activity of the promoter depends on the state the promoter is in. In particular, we remember that we assigned the transcriptional activity $a_i \in \mathbb{R}_{\geq 0}$ to promoter state $z_i$. The instantaneous overall transcriptional activity, which we denote by $a(\boldsymbol{c})$ follows from the previous as

$$a(\boldsymbol{c}) = \sum_{j=1}^{n} a_j\,Z_j$$
$$= a_i\,Z_i$$

for $i$ such that $Z_i = 1$. The property that $Z_i = 1$ implies $Z_j = 0$ for all $j \neq i$ follows directly from the one-hot encoding and we will make use of it extensively within the following.

In the context of the moment equations presented later, the promoter has a special role as it is fully described by its steady state distribution. To emphasize this, we denote the steady state distribution of the promoter by $\boldsymbol{\pi} = (\pi_1, \ldots, \pi_n)$ with $\boldsymbol{\pi}\,\boldsymbol{\Lambda} = 0$, where $\boldsymbol{\Lambda} = \boldsymbol{\Lambda}(\boldsymbol{c})$ is the propensity matrix. In particular $\pi_i = \lim_{t \Rightarrow \infty} \mathrm{p}(z_i \mid \boldsymbol{c}, t)$, where we drop the dependence on $\boldsymbol{c}$ to emphasize brevity.

**Mean Promoter Activity** With the steady state distribution and the instantaneous promoter activity at hand, we can derive the average promoter activity $\alpha(\boldsymbol{c})$.

$$\begin{aligned}
\alpha(\boldsymbol{c}) &= \mathsf{E}[a(\boldsymbol{c}) \mid \boldsymbol{c}] \\
&= \mathsf{E}\left[\sum_{j=1}^{n} a_j \, Z_j \middle| \boldsymbol{c}\right] \\
&= \sum_{j=1}^{n} a_j \; \mathsf{E}[Z_j \mid \boldsymbol{c}] \\
&= \sum_{j=1}^{n} a_j \, \pi_j
\end{aligned}$$

For this, we make use of

$$\begin{aligned}
\mathsf{E}[Z_j \mid \boldsymbol{c}] &= \sum_{x=0}^{1} x \; \mathrm{P}[Z_j = x \mid \boldsymbol{c}, t] \\
&= 0\,(1 - \pi_j) + 1\,\pi_j \\
&= \pi_j
\end{aligned}$$

where $\mathrm{P}[Z_j = 0 \mid \boldsymbol{c}, t] = 1 - \pi_j$ follows from the one-hot encoding as $Z_j = 0$ implies $Z_i = 1$ for some $i \neq j$.

**Distributions and Moments of the Promoter's States** Already used in the previous equation, we here state the distribution of the state species $Z_i$ completely.

$$
\begin{aligned}
\mathrm{P}[Z_i = x_i \mid \boldsymbol{c}, t] &= \begin{cases} 1 - \pi_i & x_i = 0 \\ \pi_i & x_i = 1 \end{cases} \\
&= (1 - x_i)\,(1 - \pi_i) + x_i\,\pi_i
\end{aligned}
$$

As required later, we next consider the conditional distribution between two state species $Z_i$ and $Z_j$. However, this requires the distinction of the two cases $i = j$ and $i \neq j$. For $i = j$, $\mathrm{P}[Z_i = x_i \mid Z_j = x_j, \boldsymbol{c}, t]$ is given by

$$
\begin{aligned}
\mathrm{P}[Z_i = x_i \mid Z_j = x_j, \boldsymbol{c}, t] &= \begin{cases} 1 & x_i = 0 \text{ and } x_j = 0 \\ 0 & x_i = 1 \text{ and } x_j = 0 \\ 0 & x_i = 0 \text{ and } x_j = 1 \\ 1 & x_i = 1 \text{ and } x_j = 1 \end{cases} \\
&= \delta(x_j - x_i)
\end{aligned}
$$

and for $k \neq j$ by

$$
\begin{aligned}
\mathrm{P}[Z_i = x_i \mid Z_j = x_j, \boldsymbol{c}, t] &= \begin{cases} 1 - \frac{\pi_i}{1-\pi_j} & x_i = 0 \text{ and } x_j = 0 \\ \frac{\pi_i}{1-\pi_j} & x_i = 1 \text{ and } x_j = 0 \\ 1 & x_i = 0 \text{ and } x_j = 1 \\ 0 & x_i = 1 \text{ and } x_j = 1 \end{cases} \\
&= (1 - x_i) + (1 - x_j)\,(2\,x_i - 1)\,\frac{\pi_i}{1 - \pi_j}.
\end{aligned}
$$

With these two distributions, we can evaluate the moment $\mathsf{E}[Z_i\, Z_j]$ as follows.

$$\begin{aligned}
\mathsf{E}\left[Z_i\, Z_j \mid \boldsymbol{c}\right] &= \sum_{x_i=0}^{1}\sum_{x_j=0}^{1} x_i\, x_j\ \mathrm{P}[Z_i = x_i, Z_j = x_j \mid \boldsymbol{c}, t] \\
&= \sum_{x_i=0}^{1}\sum_{x_j=0}^{1} x_i\, x_j\ \mathrm{P}[Z_i = x_i \mid Z_j = x_j, \boldsymbol{c}, t]\ \mathrm{P}[Z_j = x_j \mid \boldsymbol{c}, t] \\
&= \mathrm{P}[Z_i = 1 \mid Z_j = 1, \boldsymbol{c}, t]\ \mathrm{P}[Z_j = 1 \mid \boldsymbol{c}, t] \\
&= \begin{cases} \pi_j & k = j \\ 0 & k \neq j \end{cases}
\end{aligned}$$

Finally, we consider the moment $\mathsf{E}[a(\boldsymbol{c})\, Z_i]$ for which we use the result of $\mathsf{E}\left[Z_i\, Z_j\right]$.

$$\begin{aligned}
\mathsf{E}[a(\boldsymbol{c})\, Z_i \mid \boldsymbol{c}] &= \mathsf{E}\left[ Z_i \sum_{j=1}^{n} a_j\, Z_j \middle| \boldsymbol{c} \right] \\
&= \sum_{j=1}^{n} a_j\ \mathsf{E}\left[Z_i\, Z_j \mid \boldsymbol{c}\right] \\
&= a_i\ \mathsf{E}\left[Z_i\, Z_i \mid \boldsymbol{c}\right] + \sum_{j=1, j\neq k}^{n} a_j\ \mathsf{E}\left[Z_i\, Z_j \mid \boldsymbol{c}\right] \\
&= a_i\, \pi_i
\end{aligned}$$

**Rational Fraction Representation of the Promoter Activity** Considering a promoter with $n$ states and a single cognate transcription factor whose abundance is represented by $c$. Following the method of Kirchoff for deriving the steady state distribution, probability of being in state $z_i$ is of the form

$$\pi_i = \frac{\sum_{j=0}^{n} \alpha_j^{(i)}\, c^j}{\sum_{j=0}^{n} \left(\alpha_j^{(i)} + \beta_j^{(i)}\right)\, c^j}$$

with $\alpha_j^{(i)}, \beta_j^{(i)} \in \mathrm{R}_{\geq 0}$ and $m$ being the number of transitions depending on the concentration $c$, which is here assumed to enter linearly. Assigning one of multiple classes to each state, the probability of encountering a state to which class $k$ is associated is of the form

$$p_k = \frac{\sum_{j=0}^{m} \bar{\alpha}_j^{(k)}\, c^j}{\sum_{j=0}^{m} \left(\bar{\alpha}_j^{(k)} + \bar{\beta}_j^{(k)}\right) c^j}$$

where the $\bar{\alpha}_j^{(k)}$ and $\bar{\beta}_j^{(k)}$ are given by

$$\bar{\alpha}_j^{(k)} = \sum_{i \in \mathcal{I}_k} \alpha_j^{(i)} \qquad\qquad \bar{\beta}_j^{(k)} = \sum_{i=1, i \notin \mathcal{I}_k}^{m} \alpha_j^{(i)}$$

where $\mathcal{I}_k$ is the index set of all states class $k$ is assigned to and the $\alpha_j^{(i)}$ and $\beta_j^{(i)}$ as introduced previously.

Splitting the promoter in ON and OFF states with respective transcription rates $a_1$ and $a_2$ (without loss of generality $a_2 > a_1$) and assign on of these states as class to the promoter states, the average transcription rate is given by

$$y = \frac{\sum_{j=0}^{m} (a_2\, \overline{\alpha}_j + a_2\, \underline{\alpha}_j)\, c^j}{\sum_{j=0}^{m} (\overline{\alpha}_j + \underline{\alpha}_j)\, c^j}$$

where the $\overline{\alpha}_j = \sum_{i \in \overline{\mathcal{X}}} \alpha_j^{(i)}$ are the coefficients of the ON states and $\underline{\alpha}_j = \sum_{i \in \underline{\mathcal{X}}} \beta_j^{(i)}$ the one of the OFF states. As the $\overline{\alpha}$ and $\underline{\alpha}_j$ are sums of non-negative values, they are itself non-negative. This provides a convenient representation of the average promoter activity in terms of a rational fraction of two positive definite polynomials, depending in total on $2\,(m+1)+2$ parameters. We will extend on the consequence of this in the following example considering the three binding site promoter model presented in the main part of this work.

**Three Binding Site Model** The three binding site model acting as example in this work features eight states and six of the twenty reactions depend on the transcription factor concentration, thus $m = 6$. This yields $y$ as the fraction of two sixth degree polynomials consisting of $2 \cdot 7 = 14$ parameters defining the shape and two parameters defining the output range. This is 16 parameters in total. The corresponding CTMC features 20 reactions and as such 20 reaction rate constants describing the behaviour of the CTMC. In the non-equilibrium case, they can be chosen arbitrarily but positive.

Comparing the 16 parameters defining the average response curve with the 22 parameters of the full model, we observe that the model is not fully determined by considering the mean only. However, this gives space for the incorporation of further moments such as the variance, which depends on the whole set of reaction rate constants as we will see later on.

#### S2.3 Steady State RNA Dynamics

For the RNA dynamics, the transcription and RNA degradation reactions are of relevance. We refer to the random variable representing the RNA abundance by $R$ and use the instantaneous and average RNA production rates $a(\boldsymbol{c})$ and $\alpha(\boldsymbol{c})$ from the previous. For RNA degradation, we include the degradation rate $\delta_1$.

**First Raw Moment** For the first raw moment, we set $g(\mathbf{X}(t)) = R$ and obtain the Chapman Kolmogorov Backward equation as

$$\begin{aligned}
\frac{d}{dt}\mathsf{E}[R \mid \boldsymbol{c}] &= \sum_{i,j=1,j\neq i}^{n} \mathsf{E}[\Lambda_{ij}\,((R+0)-R) \mid \boldsymbol{c}] \\
&\quad + \mathsf{E}[\mu_1\, a(\boldsymbol{c})\,((R+1)-R) \mid \boldsymbol{c}] + \mathsf{E}[\delta_1\, R\,((R-1)-R) \mid \boldsymbol{c}] \\
&\quad + \mathsf{E}[\mu_2\, R\,((R+0)-R) \mid \boldsymbol{c}] + \mathsf{E}[\delta_2\, P\,((R+0)-R) \mid \boldsymbol{c}] \\
&= \mu_1\, \mathsf{E}[a(\boldsymbol{c}) \mid \boldsymbol{c}] - \delta_1\, \mathsf{E}[R \mid \boldsymbol{c}]
\end{aligned}$$

where the steady state condition (e.g. $\frac{d}{dt}\mathsf{E}[R \mid \boldsymbol{c}] = 0$) and $\mathsf{E}[a(\boldsymbol{c}) \mid \boldsymbol{c}] = \alpha(\boldsymbol{c})$ yields

$$\mathsf{E}[R \mid \boldsymbol{c}] = \frac{\mu_1}{\delta_1}\,\alpha(\boldsymbol{c})$$

as the RNA's mean.

**Second Raw Moment** To derive the protein's variance, we can take the intermediate step of determining the second raw moment first. The monomial function associated to this moment is $g(\mathbf{X}(t)) = R^2$ and we derive

$$\begin{aligned}\frac{d}{dt}\,\mathsf{E}[R^2 \mid \boldsymbol{c}] &= \sum_{i,j=1,j\neq i}^{n} \mathsf{E}[\Lambda_{ij}\,((R+0)^2 - R^2) \mid \boldsymbol{c}] \\ &\qquad + \mathsf{E}[\mu_1\, a(\boldsymbol{c})\,((R+1)^2 - R^2) \mid \boldsymbol{c}] + \mathsf{E}[\delta_1\, R\,((R-1)^2 - R^2) \mid \boldsymbol{c}] \\ &\qquad + \mathsf{E}[\mu_2\, R\,((R+0)^2 - R^2) \mid \boldsymbol{c}] + \mathsf{E}[\delta_2\, P\,((R+0)^2 - R^2) \mid \boldsymbol{c}] \\ &= \mu_1\; \mathsf{E}[a(\boldsymbol{c})\,(R^2 + 2\,R + 1 - R^2) \mid \boldsymbol{c}] + \delta_1\; \mathsf{E}[R\,(R^2 - 2\,R + 1 - R^2) \mid \boldsymbol{c}] \\ &= \mu_1\,(2\;\mathsf{E}[a(\boldsymbol{c})\,R \mid \boldsymbol{c}] + \mathsf{E}[a(\boldsymbol{c}) \mid \boldsymbol{c}]) + \delta_1\,(\mathsf{E}[R \mid \boldsymbol{c}] - 2\;\mathsf{E}[R^2 \mid \boldsymbol{c}]).\end{aligned}$$

In the steady state, we obtain after some rearrangement the relation

$$\mathsf{E}[R^2 \mid \boldsymbol{c}] = \frac{1}{2}\;\mathsf{E}[R \mid \boldsymbol{c}] + \frac{\mu_1}{2\,\delta_1}\;\mathsf{E}[a(\boldsymbol{c}) \mid \boldsymbol{c}] + \frac{\mu_1}{\delta_1}\;\mathsf{E}[a(\boldsymbol{c})\,R \mid \boldsymbol{c}]$$

Inserting the moments already known and the ones derived subsequently, we obtain

$$\mathsf{E}[R^2 \mid \boldsymbol{c}] = \frac{\mu_1}{\delta_1}\,\alpha(\boldsymbol{c}) + \frac{\mu_1^2}{\delta_1}\,\boldsymbol{a}^T\left(\delta_1\,\boldsymbol{I}_n - \tilde{\boldsymbol{K}}^T\right)^{-1}(\boldsymbol{a}\odot\boldsymbol{\pi})$$

where $\boldsymbol{a} = [a_1 \quad \dots \quad a_n]^T$ is the vector of state activities and $\odot$ denotes the Hadamard product of two vectors.

**Proportionality of Promoter Activity and RNA Abundance** The moment $g(\boldsymbol{X}(t)) = a(\boldsymbol{c})\, R$ can be stated in terms of the moment $\mathsf{E}\,[Z_j\, R]$ as illustrated by the relation

$$\begin{aligned}
\mathsf{E}[a(\boldsymbol{c})\, R \mid \boldsymbol{c}] &= \mathsf{E}\left[ \sum_{j=1}^{n} a_j\, Z_j\, R \middle| \boldsymbol{c} \right] \\
&= \sum_{j=1}^{n} a_j\; \mathsf{E}\,[Z_j\, R \mid \boldsymbol{c}] \\
&= \boldsymbol{a}^T \begin{bmatrix} \mathsf{E}[Z_1\, R \mid \boldsymbol{c}] \\ \vdots \\ \mathsf{E}[Z_n\, R \mid \boldsymbol{c}] \end{bmatrix}
\end{aligned}$$

To determine the moments $\mathsf{E}\,[Z_j\, R \mid \boldsymbol{c}]$, we again make use of the Chapman Kolmogorov backward equation for $g(\boldsymbol{X}(t)) = Z_k\, R$. In addition, we consider the moments derived in Section S2.2.

$$\begin{aligned}
\frac{d}{dt}\, \mathsf{E}[Z_k\, R \mid \boldsymbol{c}] &= \sum_{i,j=1, j\neq i}^{n} \mathsf{E}[Z_i\, \tilde{k}_{ij}\, ((Z_k + \nu_{ij_k})\, R - Z_k\, R) \mid \boldsymbol{c}] \\
&\quad + \mathsf{E}[\mu_1\, a(\boldsymbol{c})\, (Z_k\, (R+1) - Z_k\, R) \mid \boldsymbol{c}] + \mathsf{E}[\delta_1\, R\, (Z_k\, (R-1) - Z_k\, R) \mid \boldsymbol{c}] \\
&\quad + \mathsf{E}[\mu_2\, R\, (Z_k\, R - Z_k\, R) \mid \boldsymbol{c}] + \mathsf{E}[\delta_2\, P\, (Z_k\, R - Z_k\, R) \mid \boldsymbol{c}] \\
&= \mu_1\; \mathsf{E}[a(\boldsymbol{c})\, Z_k \mid \boldsymbol{c}] - \delta_1\; \mathsf{E}[Z_k\, R \mid \boldsymbol{c}] + \sum_{i,j=1, j\neq i}^{n} \nu_{ij_k}\; \mathsf{E}[Z_i\, \tilde{k}_{ij}\, R \mid \boldsymbol{c}] \\
&= \mu_1\; \mathsf{E}[a(\boldsymbol{c})\, Z_k \mid \boldsymbol{c}] - \delta_1\; \mathsf{E}[Z_k\, R \mid \boldsymbol{c}] + \sum_{i=1, i\neq k}^{n} \nu_{ik_k}\; \mathsf{E}[Z_i\, \tilde{k}_{ik}\, R \mid \boldsymbol{c}] + \sum_{i,j=1, j\neq i, j\neq k}^{n} \nu_{ij_k}\; \mathsf{E}[Z_i\, \tilde{k}_{ij}\, R \mid \boldsymbol{c}] \\
&= \mu_1\; \mathsf{E}[a(\boldsymbol{c})\, Z_k \mid \boldsymbol{c}] - \delta_1\; \mathsf{E}[Z_k\, R \mid \boldsymbol{c}] + \sum_{i=1, i\neq k}^{n} \mathsf{E}[Z_i\, \tilde{k}_{ik}\, R \mid \boldsymbol{c}] - \sum_{j=1, j\neq k}^{n} \mathsf{E}[Z_k\, \tilde{k}_{kj}\, R \mid \boldsymbol{c}] \\
&= \mu_1\; \mathsf{E}[a(\boldsymbol{c})\, Z_k \mid \boldsymbol{c}] - \delta_1\; \mathsf{E}[Z_k\, R \mid \boldsymbol{c}] + \sum_{i=1, i\neq k}^{n} \mathsf{E}[Z_i\, \tilde{k}_{ik}\, R \mid \boldsymbol{c}] - \mathsf{E}[R\, Z_k \sum_{j=1, j\neq k}^{n} \tilde{k}_{kj} \mid \boldsymbol{c}] \\
&= \mu_1\; \mathsf{E}[a(\boldsymbol{c})\, Z_k \mid \boldsymbol{c}] - \delta_1\; \mathsf{E}[Z_k\, R \mid \boldsymbol{c}] + \sum_{i=1, i\neq k}^{n} \mathsf{E}[Z_i\, \tilde{k}_{ik}\, R \mid \boldsymbol{c}] + \mathsf{E}[R\, Z_k\, \tilde{k}_{kk} \mid \boldsymbol{c}] \\
&= \mu_1\, a_k\, \pi_k - \delta_1\; \mathsf{E}[Z_k\, R \mid \boldsymbol{c}] + \sum_{i=1}^{n} \tilde{k}_{ik}\; \mathsf{E}[Z_i\, R \mid \boldsymbol{c}]
\end{aligned}$$

We obtain this result by using $\sum_{j=1,j\neq k}^{n} \tilde{k}_{kj} = -\tilde{k}_{kk}$ and

$$\nu_{ij_k} = \begin{cases} 1 & k = j \\ -1 & k = i \\ 0 & \text{else} \end{cases}$$

which follows from the definition of $\boldsymbol{\nu}_{ij}$ in Equation (S3).

Making use of the steady state assumption, we obtain

$$\mu_1\, a_k\, \pi_k = \delta_1\ \mathsf{E}[Z_k\, R \mid \boldsymbol{c}] - \sum_{i=1}^{n} \tilde{k}_{ik}\ \mathsf{E}[Z_i\, R \mid \boldsymbol{c}]$$

for each $k = 1, \ldots, n$, giving rise to a system of $n$ linear equations in the moments $\mathsf{E}[Z_k\, R \mid \boldsymbol{c}]$. Recognizing, that the sum goes over all elements in the $k$th column of matrix $\tilde{\boldsymbol{K}}$ and the product realizes a scalar product of two vectors, we obtain the $\mathsf{E}[Z_k\, R \mid \boldsymbol{c}]$ by solving

$$\left(\delta_1\, \boldsymbol{I}_n - \tilde{\boldsymbol{K}}^T\right) \begin{bmatrix} \mathsf{E}[Z_1\, R \mid \boldsymbol{c}] \\ \vdots \\ \mathsf{E}[Z_n\, R \mid \boldsymbol{c}] \end{bmatrix} = \mu_1 \begin{bmatrix} a_1\, \pi_1 \\ \vdots \\ a_n\, \pi_n \end{bmatrix}.$$

In turn, we obtain the desired quantities as

$$\begin{bmatrix} \mathsf{E}[Z_1\, R \mid \boldsymbol{c}] \\ \vdots \\ \mathsf{E}[Z_n\, R \mid \boldsymbol{c}] \end{bmatrix} = \mu_1 \left(\delta_1\, \boldsymbol{I}_n - \tilde{\boldsymbol{K}}^T\right)^{-1} \begin{bmatrix} a_1\, \pi_1 \\ \vdots \\ a_n\, \pi_n \end{bmatrix}.$$

**Variance** With the first and second centered moment at hand, we obtain the variance of the RNA abundance as

$$\begin{aligned}\mathsf{Var}[R \mid \boldsymbol{c}] &= \mathsf{E}[R^2 \mid \boldsymbol{c}] - E[R \mid \boldsymbol{c}]^2 \\ &= \frac{\mu_1}{\delta_1}\,\alpha(\boldsymbol{c})\left(1 - \frac{\mu_1}{\delta_1}\,\alpha(\boldsymbol{c})\right) + \frac{\mu_1^2}{\delta_1}\,\boldsymbol{a}^T\left(\delta_1\,\boldsymbol{I}_n - \tilde{\boldsymbol{K}}^T\right)^{-1}(\boldsymbol{a}\odot\boldsymbol{\pi}(\boldsymbol{c})).\end{aligned}$$

#### S2.4 Steady State Protein Dynamics

In this section, we derive the moment equations for the protein abundance, to which we refer as $P$. The protein abundance is affected by the two reactions translation (reaction II) and protein degradation (reaction IV). The important reaction rate constants are in this context $\mu_2$ for translation and $\delta_2$ for degradation.

**First Raw Moment** We obtain the first raw moment, respectively the average, by using $g(\mathbf{X}(t)) = P$. The corresponding Chapman Kolmogorov backward equation reads

$$\begin{aligned}\frac{d}{dt}\,\mathsf{E}[P \mid \boldsymbol{c}] &= \sum_{i,j=1,j\neq i}^{n} \mathsf{E}[\Lambda_{ij}\,(P - P) \mid \boldsymbol{c}] \\ &\quad + \mathsf{E}[\mu_1\,a(\boldsymbol{c})\,(P - P) \mid \boldsymbol{c}] + \mathsf{E}[\delta_1\,R\,(P - P) \mid \boldsymbol{c}] \\ &\quad + \mathsf{E}[\mu_2\,R\,((P+1) - P) \mid \boldsymbol{c}] + \mathsf{E}[\delta_2\,P\,((P-1) - P) \mid \boldsymbol{c}] \\ &= \mu_2\,\mathsf{E}[R \mid \boldsymbol{c}] - \delta_2\,\mathsf{E}[P \mid \boldsymbol{c}]\end{aligned}$$

and we obtain from the steady state condition $\frac{d}{dt}\,\mathsf{E}[P \mid \boldsymbol{c}] = 0$ the relationship

$$\begin{aligned}\mathsf{E}[P \mid \boldsymbol{c}] &= \frac{\mu_2}{\delta_2}\,\mathsf{E}[R \mid \boldsymbol{c}] \\ &= \frac{\mu_2}{\delta_2}\,\frac{\mu_1}{\delta_1}\,\alpha(\boldsymbol{c}).\end{aligned}$$

**Second Raw Moment** For the second raw moment we consider the monomial function $g(\mathbf{X}(t)) = P^2$ and obtain the following differential equation.

$$\begin{aligned}
\frac{d}{dt}\,\mathsf{E}[P \mid \boldsymbol{c}] &= \sum_{i,j=1,j\neq i}^{n} \mathsf{E}[\Lambda_{ij}\,(P^2 - P^2) \mid \boldsymbol{c}] \\
&\quad + \mathsf{E}[\mu_1\, a(\boldsymbol{c})\,(P^2 - P^2) \mid \boldsymbol{c}] + \mathsf{E}[\delta_1\, R\,(P^2 - P^2) \mid \boldsymbol{c}] \\
&\quad + \mathsf{E}[\mu_2\, R\,((P+1)^2 - P^2) \mid \boldsymbol{c}] + \mathsf{E}[\delta_2\, P\,((P-1)^2 - P^2) \mid \boldsymbol{c}] \\
&= \mathsf{E}[\mu_2\, R\,(P^2 + 2\,P + 1 - P^2) \mid \boldsymbol{c}] + \mathsf{E}[\delta_2\, P\,(P^2 - 2\,P + 1 - P^2) \mid \boldsymbol{c}] \\
&= \mu_2\,(2\;\mathsf{E}[R\,P \mid \boldsymbol{c}] + \mathsf{E}[R) \mid \boldsymbol{c}]) + \delta_2\,(\mathsf{E}[P \mid \boldsymbol{c}] - 2\;\mathsf{E}[P\,P \mid \boldsymbol{c}])
\end{aligned}$$

Making use of the steady state, we derive

$$\mathsf{E}[P^2 \mid \boldsymbol{c}] = \frac{1}{2}\;\mathsf{E}[P \mid \boldsymbol{c}] + \frac{\mu_2}{2\,\delta_2}\;\mathsf{E}[R \mid \boldsymbol{c}] + \frac{\mu_2}{\delta_2}\;\mathsf{E}[R\,P \mid \boldsymbol{c}]$$

and by inserting $\mathsf{E}[R\,P \mid \boldsymbol{c}]$ as derived subsequently and the moments already known from the previous, we get

$$\begin{aligned}
\mathsf{E}[P^2 \mid \boldsymbol{c}] &= \frac{\mu_2}{\delta_2}\,\frac{\mu_1}{\delta_1}\,\alpha(\boldsymbol{c}) + \frac{\mu_1\,\mu_2^2}{\delta_1\,\delta_2\,(\delta_1+\delta_2)}\,\alpha(\boldsymbol{c}) \\
&\quad + \frac{\mu_1^2\,\mu_2^2}{\delta_1\,\delta_2\,(\delta_1+\delta_2)}\,\boldsymbol{a}^T \left(\delta_1\,\boldsymbol{I}_n - \tilde{\boldsymbol{K}}^T\right)^{-1} (\boldsymbol{a} \odot \boldsymbol{\pi})) \\
&\quad + \frac{\mu_1^2\,\mu_2^2}{\delta_2\,(\delta_1+\delta_2)}\,\boldsymbol{a}^T \left(\delta_2\,\boldsymbol{I}_n - \tilde{\boldsymbol{K}}^T\right)^{-1} \left(\delta_1\,\boldsymbol{I}_n - \tilde{\boldsymbol{K}}^T\right)^{-1} (\boldsymbol{a} \odot \boldsymbol{\pi}) \\
&= \frac{\mu_2}{\delta_2}\,\frac{\mu_1}{\delta_1}\,\alpha(\boldsymbol{c}) + \frac{\mu_1\,\mu_2^2}{\delta_1\,\delta_2\,(\delta_1+\delta_2)}\,\alpha(\boldsymbol{c}) \\
&\quad + \frac{\mu_1^2\,\mu_2^2}{\delta_2\,(\delta_1+\delta_2)}\,\boldsymbol{a}^T \left(\frac{1}{\delta_1} + \left(\delta_2\,\boldsymbol{I}_n - \tilde{\boldsymbol{K}}^T\right)^{-1}\right) \left(\delta_1\,\boldsymbol{I}_n - \tilde{\boldsymbol{K}}^T\right)^{-1} (\boldsymbol{a} \odot \boldsymbol{\pi})
\end{aligned}$$

where again $\boldsymbol{a} = [a_1 \quad \ldots \quad a_n]^T$ is the vector of state activities and $\odot$ denotes the Hadamard product of two vectors.

**Proportionality of RNA and Protein Abundance** To derive the expression for $\mathsf{E}[R\,P \mid \boldsymbol{c}]$ we set $g(\boldsymbol{X}(t)) = R\,P$ for which the Chapman Kolmogorov equation follows as

$$\begin{aligned}\frac{d}{dt}\,\mathsf{E}[R\,P \mid \boldsymbol{c}] = \sum_{i,j=1,j\neq i}^{n} &\mathsf{E}[\Lambda_{ij}\,(R\,P - R\,P) \mid \boldsymbol{c}] \\ &+ \mathsf{E}[\mu_1\,a(\boldsymbol{c})\,((R+1)\,P - R\,P) \mid \boldsymbol{c}] + \mathsf{E}[\delta_1\,R\,((R-1)\,P - R\,P) \mid \boldsymbol{c}] \\ &+ \mathsf{E}[\mu_2\,R\,(R\,(P+1) - R\,P) \mid \boldsymbol{c}] + \mathsf{E}[\delta_2\,P\,(R\,(P-1) - R\,P) \mid \boldsymbol{c}] \\ = \mu_1\;&\mathsf{E}[a(\boldsymbol{c})\,P \mid \boldsymbol{c}] - \delta_1\;\mathsf{E}[R\,P \mid \boldsymbol{c}] + \mu_2\;\mathsf{E}[R^2 \mid \boldsymbol{c}] + \delta_2\;\mathsf{E}[R\,P \mid \boldsymbol{c}].\end{aligned}$$

Reshaping and the steady state condition give us then

$$\mathsf{E}[R\,P \mid \boldsymbol{c}] = \frac{\mu_1}{\delta_1 + \delta_2}\;\mathsf{E}[a(\boldsymbol{c})\,P \mid \boldsymbol{c}] + \frac{\mu_2}{\delta_1 + \delta_2}\;\mathsf{E}[R^2 \mid \boldsymbol{c}].$$

The derivation of $\mathsf{E}[a(\boldsymbol{c})\,P \mid \boldsymbol{c}]$ follows the same pattern as $\mathsf{E}[a(\boldsymbol{c})\,R \mid \boldsymbol{c}]$ before. We begin with inserting the definition of $a(\boldsymbol{c})$ into $\mathsf{E}[a(\boldsymbol{c})\,P \mid \boldsymbol{c}]$ to obtain

$$\begin{aligned}\mathsf{E}[a(\boldsymbol{c})\,P \mid \boldsymbol{c}] &= \mathsf{E}\left[\sum_{j=1}^{n} a_j\,Z_j\,P \,\middle|\, \boldsymbol{c}\right] \\ &= \sum_{j=1}^{n} a_j\;\mathsf{E}[Z_j\,P \mid \boldsymbol{c}] \\ &= \boldsymbol{a}^T \begin{bmatrix}\mathsf{E}[Z_1\,P \mid \boldsymbol{c}] \\ \vdots \\ \mathsf{E}[Z_n\,P \mid \boldsymbol{c}]\end{bmatrix}.\end{aligned}$$

For $\mathsf{E}[Z_j\,P \mid \boldsymbol{c}]$, we evaluate the corresponding Chapman Kolmogorov backward equation and get for each $k = 1, \ldots, n$

$$
\begin{aligned}
\frac{d}{dt}\,\mathsf{E}[Z_k\,P \mid \boldsymbol{c}] = &\sum_{i,j=1,j\neq i}^{n} \mathsf{E}[Z_i\,\tilde{k}_{ij}\,((Z_k+\nu_{ij_k})\,P - Z_k\,P) \mid \boldsymbol{c}] \\
&+ \mathsf{E}[\mu_1\,a(\boldsymbol{c})\,(Z_k\,P - Z_k\,P) \mid \boldsymbol{c}] + \mathsf{E}[\delta_1\,R\,(Z_k\,P - Z_k\,P) \mid \boldsymbol{c}] \\
&+ \mathsf{E}[\mu_2\,R\,(Z_k\,(P+1) - Z_k\,P) \mid \boldsymbol{c}] + \mathsf{E}[\delta_2\,P\,(Z_k\,(P-1) - Z_k\,P) \mid \boldsymbol{c}] \\
= &\,\mu_2\,\mathsf{E}[R\,Z_k \mid \boldsymbol{c}] - \delta_2\,\mathsf{E}[P\,Z_k \mid \boldsymbol{c}] + \sum_{i,j=1,j\neq i}^{n} \nu_{ij_k}\,\mathsf{E}[P\,Z_i\,\tilde{k}_{ij} \mid \boldsymbol{c}] \\
= &\,\mu_2\,\mathsf{E}[R\,Z_k \mid \boldsymbol{c}] - \delta_2\,\mathsf{E}[P\,Z_k \mid \boldsymbol{c}] + \sum_{i=1,i\neq k}^{n} \nu_{ik_k}\,\mathsf{E}[P\,Z_i\,\tilde{k}_{ik} \mid \boldsymbol{c}] + \sum_{i,j=1,j\neq i,j\neq k}^{n} \nu_{ij_k}\,\mathsf{E}[P\,Z_i\,\tilde{k}_{ij} \mid \boldsymbol{c}] \\
= &\,\mu_2\,\mathsf{E}[R\,Z_k \mid \boldsymbol{c}] - \delta_2\,\mathsf{E}[P\,Z_k \mid \boldsymbol{c}] + \sum_{i=1,i\neq k}^{n} \mathsf{E}[P\,Z_i\,\tilde{k}_{ik} \mid \boldsymbol{c}] - \sum_{j=1,j\neq k}^{n} \mathsf{E}[P\,Z_k\,\tilde{k}_{kj} \mid \boldsymbol{c}] \\
= &\,\mu_2\,\mathsf{E}[R\,Z_k \mid \boldsymbol{c}] - \delta_2\,\mathsf{E}[P\,Z_k \mid \boldsymbol{c}] + \sum_{i=1,i\neq k}^{n} \mathsf{E}[P\,Z_i\,\tilde{k}_{ik} \mid \boldsymbol{c}] - \mathsf{E}[P\,Z_k \sum_{j=1,j\neq k}^{n} \tilde{k}_{kj} \mid \boldsymbol{c}] \\
= &\,\mu_2\,\mathsf{E}[R\,Z_k \mid \boldsymbol{c}] - \delta_2\,\mathsf{E}[P\,Z_k \mid \boldsymbol{c}] + \sum_{i=1,i\neq k}^{n} \mathsf{E}[P\,Z_i\,\tilde{k}_{ik} \mid \boldsymbol{c}] + \mathsf{E}[P\,Z_k\,\tilde{k}_{kk} \mid \boldsymbol{c}] \\
= &\,\mu_2\,\mathsf{E}[R\,Z_k \mid \boldsymbol{c}] - \delta_2\,\mathsf{E}[P\,Z_k \mid \boldsymbol{c}] + \sum_{i=1}^{n} \mathsf{E}[P\,Z_i\,\tilde{k}_{ik} \mid \boldsymbol{c}].
\end{aligned}
$$

As before, the steady state gives rise to a system of $n$ equations, which can be represented by a single linear equation with the solution

$$
\begin{bmatrix} \mathsf{E}[Z_1\,P \mid \boldsymbol{c}] \\ \vdots \\ \mathsf{E}[Z_n\,P \mid \boldsymbol{c}] \end{bmatrix} = \mu_2\,\left(\delta_2\,\boldsymbol{I}_n - \tilde{\boldsymbol{K}}^T\right)^{-1} \begin{bmatrix} \mathsf{E}[Z_1\,R \mid \boldsymbol{c}] \\ \vdots \\ \mathsf{E}[Z_n\,R \mid \boldsymbol{c}] \end{bmatrix}.
$$

**Variance** After deriving the first and second order central moment, left over is their combination into the variance of the protein production. In particular, the variance is given by

$$
\begin{aligned}
\mathsf{Var}[P \mid \boldsymbol{c}] &= \mathsf{E}[P^2 \mid \boldsymbol{c}] - \mathsf{E}[P \mid \boldsymbol{c}]^2 \\
&= \frac{\mu_2}{\delta_2}\frac{\mu_1}{\delta_1}\alpha(\boldsymbol{c})\left(1 - \frac{\mu_2}{\delta_2}\frac{\mu_1}{\delta_1}\alpha(\boldsymbol{c})\right) + \frac{\mu_1\,\mu_2^2}{\delta_1\,\delta_2\,(\delta_1+\delta_2)}\alpha(\boldsymbol{c}) \\
&\quad + \frac{\mu_1^2\,\mu_2^2}{\delta_2\,(\delta_1+\delta_2)}\boldsymbol{a}^T\left(\frac{1}{\delta_1} + \left(\delta_2\,\boldsymbol{I}_n - \tilde{\boldsymbol{K}}^T\right)^{-1}\right)\left(\delta_1\,\boldsymbol{I}_n - \tilde{\boldsymbol{K}}^T\right)^{-1}(\boldsymbol{a}\odot\boldsymbol{\pi}(\boldsymbol{c}))
\end{aligned}
$$

**Protein and RNA Covariance** To adequately state the relationship between the distribution of RNA and the one of protein, the covariance is of importance. As everything is in place, we obtain after some reshaping

$$
\begin{aligned}
\mathsf{Cov}[R\,P \mid \boldsymbol{c}] &= \mathsf{E}[(R - \mathsf{E}[R \mid \boldsymbol{c}])\,(P - \mathsf{E}[P \mid \boldsymbol{c}]) \mid \boldsymbol{c}] \\
&= \mathsf{E}[R\,P \mid \boldsymbol{c}] - \mathsf{E}[R \mid \boldsymbol{c}]\;\mathsf{E}[P \mid \boldsymbol{c}] \\
&= \frac{\mu_2}{\delta_1+\delta_2}\frac{\mu_1}{\delta_1}\alpha(\boldsymbol{c}) - \frac{\mu_2}{\delta_2}\frac{\mu_1^2}{\delta_1^2}\alpha(\boldsymbol{c})^2 \\
&\quad + \mu_1^2\,\boldsymbol{a}^T\left(\frac{\mu_2}{\delta_1+\delta_2}\left(\delta_2\,\boldsymbol{I}_n - \tilde{\boldsymbol{K}}^T\right)^{-1} + \frac{1}{\delta_1}\right)\left(\delta_1\,\boldsymbol{I}_n - \tilde{\boldsymbol{K}}^T\right)^{-1}(\boldsymbol{a}\odot\boldsymbol{\pi}(\boldsymbol{c}))
\end{aligned}
$$

### S3 Stochastic Thermodynamics of Open Chemical Reaction Networks

In the main document, we utilized the entropy production rate of the (kinetic) promoter model to quantify the mean energy dissipation rate of the promoter. In this section, we pedagogically introduce the concepts of stochastic thermodynamics for open chemical reaction networks (CRNs) in order to provide an explanation of this commonly made identification. There is a theoretical gap between a kinetic model and its energetics. In the following, we point out that the energy changes associated with a microscopically reversible chemical reaction can be linked to the associated kinetic rates through a relation that establishes consistency between the kinetic and energetic descriptions. The application of this consistency relation, also referred to as local detailed balance, allows for the expression of the energy dissipation rate of a chemical reaction network that is coupled to a heat bath and a number of chemostats in terms of only its kinetic rates.

The most relevant equations in this section are the energy dissipated into the environment per reaction

$$\Delta Q^{(\pm m)}(\boldsymbol{x}) = W_{\text{chem}}^{(\pm m)} - \Delta g^{(\pm m)}(\boldsymbol{x}),$$

the mean energy dissipation rate

$$\dot{Q}(t) = \sum_{\boldsymbol{x}} \sum_{m=1}^{M} (\Delta Q^{(m)}(\boldsymbol{x})\lambda_m^+(\boldsymbol{x}) + \Delta Q^{(-m)}(\boldsymbol{x})\lambda_m^-(\boldsymbol{x}))p(\boldsymbol{X}(t) = \boldsymbol{x}),$$

the thermodynamic consistency relation

$$\ln\left(\frac{\lambda_m^+(\boldsymbol{x})}{\lambda_m^-(\boldsymbol{x})}\right) = -\beta\left(\Delta g^{(m)}(\boldsymbol{x}) + \Delta\mu^{(m)}\right) = \beta\,\Delta Q^{(m)}(\boldsymbol{x}),$$

and the identification of the energy dissipation rate with the entropy flow into the environment

$$\frac{\mathrm{d}}{\mathrm{d}t}H^{\text{env}}(t) = \beta\dot{Q}(t) = \sum_{\boldsymbol{x}} \sum_{m=1}^{M} \ln\left(\frac{\lambda_m^+(\boldsymbol{x})}{\lambda_m^-(\boldsymbol{x}+\boldsymbol{\nu}_m)}\right) J_m(\boldsymbol{x}, t).$$

#### S3.1 Closed Stochastic Chemical Reaction Networks

A stochastic chemical reaction network consists of $N \in \mathbb{N}$ chemical species $\mathsf{Z} = (\mathsf{Z}_1, \ldots, \mathsf{Z}_N)$ and $M \in \mathbb{N}$ reactions $\mathcal{R}_1, \ldots, \mathcal{R}_M$ with stoichiometric balance equations

$$\mathsf{R}_m : \quad \sum_{n=1}^{N} a_n^{(m)} \mathsf{Z}_n \quad \underset{\lambda_m^-}{\overset{\lambda_m^+}{\rightleftharpoons}} \quad \sum_{n=1}^{N} b_n^{(m)} \mathsf{Z}_n,$$

where $a^{(m)}, b^{(m)} \in \mathbb{N}^N$ contain the respective numbers of reacting molecules of each species and the numbers of the product molecules. The backward reaction microscopically reverses the forward reaction. The difference of output and input species of a reaction $\mathcal{R}_m$ forms the stoichiometric change vector $\nu_m = b^{(m)} - a^{(m)}$ of the forward reaction. Let $\{Z(t) \in \mathbb{N}^N : t \geq 0\}$ be the continuous-time Markov chain of the species' copy number vector. The forward and backward reactions are assumed to follow stochastic mass action kinetics (1–3). Consequently, the rates of the reactions, also referred to as propensity functions, have the form

$$\lambda_m^+(z) = k_m^+ \prod_{n=1}^{N} \frac{z_n!}{(z_n - a_n^{(m)})!}, \qquad \lambda_m^-(z) = k_m^- \prod_{n=1}^{N} \frac{z_n!}{(z_n - b_n^{(m)})!} \tag{S5}$$

with $z \in \mathbb{N}^N$ and stochastic rate constants $k_m^+ > 0$ and $k_m^- \geq 0$. These stochastic rate constants are related to the deterministic rate constants $\kappa_m^+ > 0, \kappa_m^- \geq 0$ and the volume $V$ of the solvent via

$$k_m^+ = \kappa_m^+ \frac{V}{\prod_{n=1}^{N} V^{a_n^{(m)}}}, \qquad k_m^- = \kappa_m^- \frac{V}{\prod_{n=1}^{N} V^{b_n^{(m)}}},$$

due to consistency of physical units in both the deterministic ODE model (of the concentrations) and the probability evolution equation of the stochastic model (4). A reaction is called microscopically reversible if $k_m^- > 0$. A CRN is called microscopically reversible if all reactions are microscopically reversible. In the following, a CRN is assumed to be microscopically reversible unless otherwise noted. We also make the simplifying assumption that all reactions are elementary, i.e., there are no reaction intermediates and each reaction has only a single transition state in the reaction coordinate diagram (5). Lastly, we assume that all reactants and products are resolved.

The probability mass function $p(z,t) = \mathrm{P}(Z(t) = z)$ of this model follows the chemical master equation (1–3)

$$\partial_t p(z,t) = \sum_{m=1}^{M} \lambda_m^+(z - \nu_m) p(z - \nu_m, t) + \lambda_m^-(z + \nu_m) p(z + \nu_m, t) - (\lambda_m^+(z) + \lambda_m^-(z)) p(z,t) \tag{S6}$$

This model makes several assumptions. Firstly, assuming a Markov process description means that the future dynamics of the mesoscopic system is sufficiently described by the current state and does not depend on the past. The microscopic states comprising any mesoscopic state are thus assumed to be degenerate. That is, the future dynamics do not depend on the particular microstate, the system is in, but just on the current mesostate $z$. Secondly, there are assumptions about the environment in which the reaction network is embedded. Since the propensity functions are independent of time and space, the embedding environment must be seen as equilibrated and spatially homogeneous from the perspective of the species $\mathsf{Z}$. That is, temperature, pressure, and volume of the solvent are constant over time and space. Similarly, the molecular composition of the solvent, which may influence the rate constants, remains constant over time and the surrounding of each molecule of $\mathsf{Z}$ homogenizes spatially on a much faster time scale than reactions occur.

A chemical reaction network is considered closed if it does not exchange molecules with the external environment and if it does not change the number of molecules of any of the chemical species that make up the solvent. A closed thermodynamic system may exchange heat with its environment. We interpret the solvent as a heat bath which maintains a constant temperature $T$.

The zeroth law of thermodynamics for chemical reaction networks dictates that a closed, microscopically reversible CRN always relaxes to equilibrium (*3*). Equilibrium statistical mechanics further requires that the equilibrium distribution of a closed chemical reaction network is (i) a Gibbs-Boltzmann distribution $\pi(z) = \lim_{t\to\infty} \mathrm{P}(Z(t) = z) \propto e^{-\beta g(z)}$, where $\beta = (k_\mathrm{B}T)^{-1}$ and $g(z)$ is the Gibbs free energy of the state $z$, and (ii) it satisfies detailed balance

$$\lambda_m^+(z)\,\pi(z) = \lambda_m^-(z + \nu_m)\,\pi(z + \nu_m) \tag{S7}$$

for all admissible states $z$ and all reversible reactions $\mathcal{R}_m$ (*3*). Plugging the Boltzmannn distribution into (S7) yields the local detailed balance condition, or thermodynamic consistency relation, for closed CRNs ([3]):

$$\ln\left(\frac{\lambda_m^+(z)}{\lambda_m^-(z+\nu_m)}\right) = -\beta(g(z+\nu_m) - g(z)) \tag{S8}$$

for all admissible states $z$ and all reversible reactions $\mathcal{R}_m$. It should be noted that equation (S8) relates kinetic and energetic constants, and therefore it is also satisfied during the transient nonequilibrium and independent of the initial conditions.

#### S3.2 Open Chemical Reaction Networks – Coupling to Chemostats

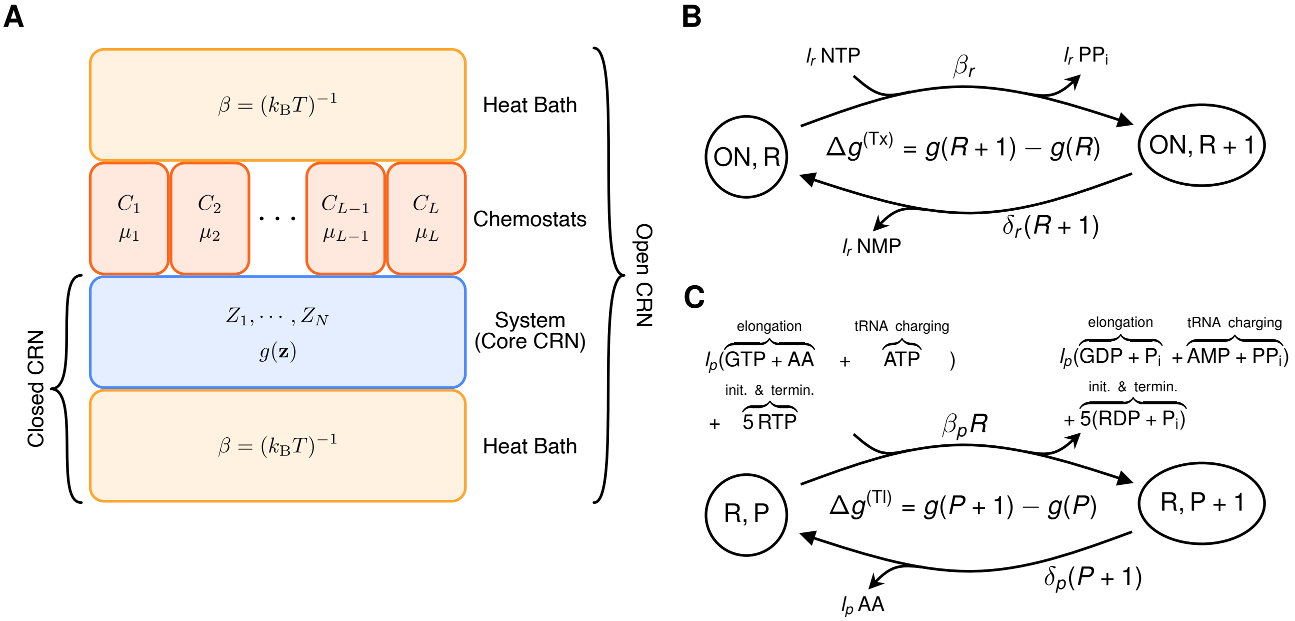


Figure S1: **Open chemical reaction networks. A:** Visualization of an open CRN, where the core system Z is coupled to a number of chemostat species. Each horizontal layer can exchange energy with its vertically adjacent layers. **BC:** State transition diagram of the copy numbers of RNA and protein with a switchable promoter, exhibiting ON and OFF states. The directed edges also show which species are exchanged with the chemostat.

So far, we have suggested that the reactions of the core reaction network may also depend on the molecular species of the solvent. For a closed network, we have assumed that the species of the solvent do not change due to the given reactions $\mathcal{R}_1, \ldots, \mathcal{R}_M$, so that the stochastic rate constants do not explicitly depend on the concentrations of the solvent species and thus do not change with time. Otherwise, for example, some solvent species might become depleted. In an open CRN, the exchange of molecules with an external reservoir is allowed. In the following, we assume that the core system is coupled to L chemostats $\mathsf{C}_1, \dots, \mathsf{C}_L$ via the solvent. The open CRN is visualized in Figure S1**A**. The copy number of chemostat molecules in the solvent is assumed to be high enough to enter the description of the system only via their real-valued concentrations $c = (c_1, \dots, c_L)$, which are constant over time. The coupled system follows the stoichiometric balance equations

$$\mathsf{R}_m : \quad \sum_{l=1}^{L} r_l^{(m)} \mathsf{C}_l + \sum_{n=1}^{N} a_n^{(m)} \mathsf{Z}_n \quad \underset{\lambda_m^-}{\overset{\lambda_m^+}{\rightleftharpoons}} \quad \sum_{l=1}^{L} s_l^{(m)} \mathsf{C}_l + \sum_{n=1}^{N} b_n^{(m)} \mathsf{Z}_n,$$

where $\gamma^{(m)} = s^{(m)} - r^{(m)}$ is the stoichiometric change vector of the chemostat species associated with $\mathcal{R}_m$. Immediately after each reaction, the chemostat is assumed to correct the copy number of all chemostat species. Consequently, the propensity functions of the coupled reactions maintain the form (S5) with the adjustment that the stochastic rate constants now depend on the chemostat concentrations insofar as they incorporate a deterministic law of mass action

$$k_m^+(c) = \kappa_m^+ \frac{V}{\prod_{n=1}^{N} V^{a_n^{(m)}}} \prod_{l=1}^{L} c_l^{r_l^{(m)}}, \qquad k_m^-(c) = \kappa_m^- \frac{V}{\prod_{n=1}^{N} V^{b_n^{(m)}}} \prod_{l=1}^{L} c_l^{s_l^{(m)}},$$

where the units of the deterministic rate constants $\kappa_m^+$, $\kappa_m^-$ are adapted accordingly. Therefore, the probability evolution is still described by (S6).

Suppose a real chemical network is empirically well described by a Markovian CRN. Let us further assume that the biomolecular species ATP, ADP and $\mathrm{P}_i$ are chemostat species and some of the state changes of the system involve ATP hydrolysis. In a microscopically reversible CRN, any ATP consuming reaction ATP $\rightleftharpoons$ ADP $+ \mathrm{P}_i$ always produces ATP in the reverse direction. Consequently, if any state change is predominantly ATP-consuming in one direction, but not ATP-producing in the reverse direction, then there are at least two distinct reactions involved. That is, the microscopically reversible CRN features two reactions with the same stochiometric change vector for the core system, but distinct change vectors for the chemostat species.

To describe the energetics of each reaction $\mathcal{R}_m$ we denote by $\mu_l$ the chemical potential of $\mathsf{C}_l$, i.e., the energy associated with a single molecule of species $\mathsf{C}_l$. Reaction $\mathcal{R}_m$, starting from state $z$, is associated with a change in the Gibbs free energy of the state

$$\Delta g^{(m)}(z) = g(z+\nu_m) - g(z) = -\Delta g^{(-m)}(z+\nu_m)$$

and the chemical work $W_{\text{chem}}^{(m)}$ done on the system by the chemostat

$$W_{\text{chem}}^{(\pm m)} = \mp \sum_l (s_l^{(m)} - r_l^{(m)})\mu_l,$$

where $(-m)$ denotes the reverse reaction. Thus the energy dissipated into the environment per reaction is

$$\Delta Q^{(\pm m)}(z) = W_{\text{chem}}^{(\pm m)} - \Delta g^{(\pm m)}(z) \tag{S9}$$

with $\Delta Q^{(-m)}(z+\nu_m) = -\Delta Q^{(m)}(z)$. Intuitively, the dissipated energy is the chemical work done on the system but not stored in the system's state. Let $0 < \tau_1^{(\pm m)} < \tau_2^{(\pm m)} < \cdots \le t$ be the random times, when reaction $\mathcal{R}_m$ happens in forward or backward direction. Then the energy dissipated by the random trajectory $\{Z(s) : 0 < s \le t\}$ of the open CRN within the time interval $(0, t]$ is

$$q(t) = \sum_{m=1}^{M} \left( \sum_{k:0<\tau_k^{(m)} \le t} \Delta Q^{(m)}(Z(\tau_k^{(m)}-)) \right) + \left( \sum_{k:0<\tau_k^{(-m)} \le t} \Delta Q^{(-m)}(Z(\tau_k^{(-m)}-)) \right) \tag{S10}$$

where $Z(t-) = \lim_{s\uparrow t} Z(s)$ denotes the left limit in time, such that $Z(\tau_k^{(m)}-)$ is the state of the CRN just before the $k$-th occurrence of reaction $\mathcal{R}_m$ in forward direction (2). The mean energy dissipation rate $\dot{Q}(t)$ of the open CRN can be stated as

$$\begin{aligned}\dot{Q}(t) = \frac{\mathrm{d}}{\mathrm{d}t}\mathsf{E}[q(t)] &= \sum_{m=1}^{M}\left(\mathsf{E}[\Delta Q^{(m)}(Z(t))\lambda_m^+(Z(t))] + \mathsf{E}[\Delta Q^{(-m)}(Z(t))\lambda_m^-(Z(t))]\right)\\ &= \sum_z \sum_{m=1}^{M} (\Delta Q^{(m)}(z)\lambda_m^+(z) + \Delta Q^{(-m)}(z)\lambda_m^-(z))p(z,t) \\ &= \sum_z \sum_{m=1}^{M} \Delta Q^{(m)}(z)\ (\lambda_m^+(z)p(z,t) - \lambda_m^-(z+\nu_m)p(z+\nu_m,t)),\end{aligned} \tag{S11}$$

where the expectation of $q$ is taken over the distribution of all trajectories $\{Z(s): 0 < s \leq t\}$. Defining the forward, the backward, and the net probability fluxes $J_m^+(z,t) = \mathrm{p}(Z(t) = z)\,\lambda_m^+(z)$, $J_m^-(z,t) = \mathrm{p}(Z(t) = z+\nu_m)\,\lambda_m^-(z+\nu_m)$, and $J_m(z,t) = J_m^+(z,t) - J_m^-(z,t)$, respectively, of reaction $\mathcal{R}_m$ , we can rewrite (S11) as

$$\dot{Q}(t) = \sum_z \sum_{m=1}^{M} \Delta Q^{(m)}(z)\ J_m(z,t)$$

The first line in Equation (S11) follows by the theory of point processes (*6*), the second line by evaluation of the expectations and the third line by shifting of the order of summation of the last term and using the property of Equation (S9) stated above. For completeness we elaborate on the first line of (S11) in the following proof.

*Proof of* (S11). We represent $Z(t)$ via its forward/backward reaction counting processes

$$R_m^\pm(t) = N_m^\pm\left(\int_0^t \lambda_m^\pm(Z(s))\ \mathrm{d}s\right),$$

where $\{N_m^\pm(t) \in \mathbb{N}, t > 0\}$ with $1 \leq m \leq M$ are independent unit rate Poisson processes. Then

$$Z(t) = Z(0) + \sum_{m=1}^{M} \nu_m\,(R_m^+(t) - R_m^-(t)) = Z(0) + \sum_{m=1}^{M}\int_0^t \nu_m\,(\ \mathrm{d}R_m^+(s) -\ \mathrm{d}R_m^-(s)).$$

We rewrite (S10) via stochastic integrals of the reaction counting processes

$$q(t) = \sum_{m=1}^{M} \int_0^t \Delta Q^{(m)}(Z(s-))\ \mathrm{d}R_m^+(s) + \int_0^t \Delta Q^{(-m)}(Z(s-))\ \mathrm{d}R_m^-(s).$$

Now with the definition in (*7*), page 27, the expectations of the stochastic integrals are rewritten as expectations of Lebesgue/Riemann integrals

$$\begin{aligned}\mathsf{E}[q(t)] &= \sum_{m=1}^{M} \mathsf{E}\left[\int_0^t \Delta Q^{(m)}(Z(s-))\ \mathrm{d}R_m^+(s) + \int_0^t \Delta Q^{(-m)}(Z(s-))\ \mathrm{d}R_m^-(s)\right]\\ &= \sum_{m=1}^{M} \mathsf{E}\left[\int_0^t \Delta Q^{(m)}(Z(s-))\lambda_m^+(Z(s-))\ \mathrm{d}s + \int_0^t \Delta Q^{(-m)}(Z(s-))\lambda_m^-(Z(s-))\ \mathrm{d}s\right],\end{aligned}$$

such that (S11) follows by exchanging the expectation with the integral via Fubini's theorem and taking the derivative. □

In open CRNs the thermodynamic consistency relation

$$\ln\left(\frac{\lambda_m^+(z)}{\lambda_m^-(z+\nu_m)}\right) = \beta\Delta\, Q^{(m)}(z) \tag{S12}$$

for all admissible states $z$ and all reversible reactions $\mathcal{R}_m$ generalizes the local detailed balance (S8) (*3*). To show consistency in the deterministic limit, we use the macroscopic limit of the Gibbs free energy of the state and the propensity functions for a large volume $V$

$$\begin{aligned} g(V\tilde{z}) &\approx V\sum_{n=1}^{N}\int_0^{\tilde{z}_n}(g_n^o + \beta^{-1}\ln(y_n))\ \mathrm{d}y_n\\ \Delta g^{(m)}(V\tilde{z}) &\approx \nu_m(V^{-1}\nabla_{\tilde{z}})g(V\tilde{z}) = \sum_{n=1}^{N}(b_n^{(m)} - a_n^{(m)})(g_n^o + \beta^{-1}\ln(\tilde{z}_n))\\ \lambda_m^+(V\tilde{z}) &\approx \kappa_m^+ V\left(\prod_{l=1}^{L} c_l^{r_l^{(m)}}\right)\left(\prod_{n=1}^{N}\tilde{z}_n^{a_n^{(m)}}\right), \qquad \lambda_m^-(V\tilde{z}) \approx \kappa_m^- V\left(\prod_{l=1}^{L} c_l^{s_l^{(m)}}\right)\left(\prod_{n=1}^{N}\tilde{z}_n^{b_n^{(m)}}\right),\end{aligned}$$

where $\tilde{z} = V^{-1}z$, $\nabla_{\tilde{z}}$ denotes the gradient operator w.r.t. the components of $\tilde{z}$, and we assumed independence of the chemical potentials of distinct species. Further we identify $\mu_l = \mu_l^o + \beta^{-1}\ln(c_l)$ and note that $-W_{\mathrm{chem}}^{(\pm m)}$ is exactly the change in Gibbs free energy of the state of the chemostat species in the macroscopic limit. Thus if the species $C_1, \ldots, C_L$ and the core system would form a closed system, then the macroscopic limit of $-\Delta Q^{(m)}(z)$ is exactly the change of macroscopic Gibbs free energy of the total system upon reaction $\mathcal{R}_m$ in forward direction. So in the macroscopic limit the thermodynamic consistency relation for open CRNs (S12), i.e.,

$$\ln\left(\frac{\lambda_m^+(V\tilde{z})}{\lambda_m^-(V\tilde{z})}\right) = \beta\left(\sum_l (s_l^{(m)} - r_l^{(m)})\mu_l - \Delta g^{(m)}(V\tilde{z})\right).,\tag{S13}$$

is consistent with (S8). Using the definition of $\mu_l$ and the expression for $\Delta g^{(m)}(V\tilde{z})$ in Equation (S13) yields the well known relation for the macroscopic rate constants (*8*)

$$\ln(K_{\text{eq}}^{(m)}) = \ln\left(\frac{\kappa_m^+}{\kappa_m^-}\right) = -\sum_{l=1}^{L}(s_l^{(m)} - r_l^{(m)})\beta\mu_l^o - \sum_{n=1}^{N}(b_n^{(m)} - a_n^{(m)})\beta g_n^o,\tag{S14}$$

where $K_{\text{eq}}^{(m)}$ is the equilibrium constant of reaction $\mathcal{R}_m$. For further consistency results we refer the reader to (*3*).

#### S3.3 Relation of Entropy Production Rate and Energy Dissipation

Plugging (S12) into (S11) reexpresses the mean energy dissipation rate (in units of $k_\text{B}T$)

$$\beta\dot{Q}(t) = \sum_z \sum_{m=1}^{M} \ln\left(\frac{\lambda_m^+(z)}{\lambda_m^-(z+\nu_m)}\right) J_m(z,t)\tag{S15}$$

based only on the kinetic parameters of the open CRN. Now (S15) is further identified as the (Shannon) entropy flow into the environment $\frac{\text{d}}{\text{d}t}H^{\text{env}}(t) = \beta\dot{Q}(t)$ (*2*, *3*). The entropy production of the whole system $H^{\text{tot}}(t)$, consisting of the core system and its environment, during the interval $[0,t]$ is given as

$$H^{\text{tot}}(t) = \int_0^t \left(\frac{\text{d}}{\text{d}s}H[Z(s)] + \frac{\text{d}}{\text{d}s}H^{\text{env}}(s)\right)\,\text{d}s,$$

where $H[Z(t)] = -\sum_z p(z,t)\ln(p(z,t))$ is the (Shannon) entropy of the core system at time $t \geq 0$. Using the CME (S6) we express the derivative of the entropy of the core system as

$$\begin{aligned}\frac{\mathrm{d}}{\mathrm{d}t}H[Z(t)] &= -\sum_z (\ln(p(z,t)) + 1)\partial_t p(z,t) \\ &= \sum_z \sum_{m=1}^{M} \ln\left(\frac{p(z,t)}{p(z+\nu_m,t)}\right)(\lambda_m^+(z)p(z,t) - \lambda_m^-(z+\nu_m)p(z+\nu_m,t)).\end{aligned}$$

Thus the entropy production rate is given by

$$\begin{aligned}\frac{\mathrm{d}}{\mathrm{d}t}H^{\mathrm{tot}}(t) &= \sum_z \sum_{m=1}^{M} \ln\left(\frac{p(z,t)\,\lambda_m^+(z)}{p(z+\nu_m,t)\,\lambda_m^-(z+\nu_m)}\right)(\lambda_m^+(z)p(z,t) - \lambda_m^-(z+\nu_m)p(z+\nu_m,t)) \\ &= \sum_z \sum_{m=1}^{M} \ln\left(\frac{J_m^+(z,t)}{J_m^-(z,t)}\right)J_m(z,t).\end{aligned} \tag{S16}$$

At stationarity we have $\lim_{t\to\infty} H[Z(t)] = -\sum_z \pi(z)\ln(\pi(z))$, implying $\lim_{t\to\infty}\frac{\mathrm{d}}{\mathrm{d}t}H[Z(t)] = 0$ and consequently

$$e_p = \lim_{t\to\infty}\frac{\mathrm{d}}{\mathrm{d}t}H^{\mathrm{tot}}(t) = \lim_{t\to\infty}\frac{\mathrm{d}}{\mathrm{d}t}H^{\mathrm{env}}(t) = \lim_{t\to\infty}\beta\dot{Q}(t),$$

which is what we wanted to show. The entropy production rate at stationarity quantifies the mean energy dissipation rate of the core chemical reaction network into the environment.

In the main document, we have provided the energy dissipation rate for an unspecified coupling to chemostat species and unspecified chemostat concentrations. It should be noted, however, that the deterministic rate constants depend explicitly on the coupling due to equation (S14). Consequently, a given open CRN with specified reactions and fixed chemostat concentrations has only one degree of freedom per reaction $\mathcal{R}_m$ and the stochastic rate constants obey

$$\frac{k_m^+}{k_m^-} = K_{\mathrm{eq}}^{(m)} \cdot \left(\prod_{n=1}^{N} V^{b_n^{(m)}-a_n^{(m)}}\right)\left(\prod_{l=1}^{L} c^{r_l^{(m)}-s_l^{(m)}}\right)$$

for a given coupling and fixed chemostat concentrations.

#### S3.4 Energy Dissipation of RNA and Protein Dynamics

Lastly, we derive the energy consumption rate of RNA and protein dynamics of a single gene. In contrast to the promoter reactions, transcription, RNA degradation, translation and protein degradation are modelled as microscopically irreversible reactions. Each is an effective reaction, which may consist of a sequence of microscopically reversible reactions. However, each of the reverse microscopic reactions is significantly less likely to occur than the corresponding forward reaction, implying that the rate of the reverse effective composite reaction is negligible.

We model the RNA and protein dynamics with the following stoichiometric balance equations, containing both, the chemostat species and the core species, $\mathsf{R}$ (RNA), $\mathsf{P}$ (protein) and the promoter in the on state ($\mathrm{G_{ON}}$):

$$\begin{aligned}
\text{Transcription:}\quad & l_r \cdot \mathrm{NTP} + \mathrm{G_{ON}} \xrightarrow{\lambda^{(\mathrm{Tx})}} \mathrm{RNA} + l_r \cdot \mathrm{PP_i} + \mathrm{G_{ON}} \\
\text{RNA deg.:}\quad & \mathrm{RNA} \xrightarrow{\lambda^{(\mathrm{Rdeg})}} l_r \cdot \mathrm{NMP} \\
\text{Translation:}\quad & \mathrm{RNA} + l_p(\mathrm{GTP} + \mathrm{AA} + \mathrm{ATP}) + 5\,\mathrm{RTP} \\
& \xrightarrow{\lambda^{(\mathrm{Tl})}} \mathrm{RNA} + \mathrm{Protein} + l_p(\mathrm{GDP} + \mathrm{P_i} + \mathrm{AMP} + \mathrm{PP_i}) + 5(\mathrm{RDP} + \mathrm{P_i}) \\
\text{Protein deg.:}\quad & \mathrm{Protein} \xrightarrow{\lambda^{(\mathrm{Pdeg})}} l_p \cdot \mathrm{AA}
\end{aligned}$$

The chemostat species are the building blocks of RNA and protein as well as energy-rich molecules and their degradation products. State transition diagrams for the RNA and protein states are depicted in Figure S1 b) and c). Denote $Z(t) = (X(t), R(t), P(t))$, where $X(t) \in \{0,1\}$ is the promoter state. Let $-\Delta\mu^{(\mathrm{Tx})}, -\Delta\mu^{(\mathrm{Tl})}$ be the chemical work done to produce an RNA molecule or a protein, respectively, i.e.,

$$\begin{aligned}
\Delta\mu^{(\mathrm{Tx})} =& l_r(\mu(\mathrm{PP_i}) - \mu(\mathrm{NTP})) \\
\Delta\mu^{(\mathrm{Tl})} =& l_r(\mu(\mathrm{GDP}) + \mu(\mathrm{P_i}) + \mu(\mathrm{AMP}) + \mu(\mathrm{PP_i}) - \mu(\mathrm{GTP}) - \mu(\mathrm{AA}) - \mu(\mathrm{ATP})) \\
& + 5(\mu(\mathrm{RDP}) + \mu(\mathrm{P_i}) - \mu(\mathrm{RTP}))
\end{aligned}$$

Further, let $\Delta g^{(\mathrm{Tx})}(Z(t))$, $\Delta g^{(\mathrm{Tl})}(Z(t))$ be the Gibbs free energy "stored" in, respectively, the RNA molecule and the protein. Lastly, $\Delta\mu^{(\mathrm{Rdeg})} = l_r\mu(\mathrm{NMP})$, $\Delta\mu^{(\mathrm{Pdeg})} = l_p\mu(\mathrm{AA})$ be the chemical energy released into the environment upon the respective degradation reactions. As an approximation we assume that the reaction free energies are independent of the copy number of the respective RNA or protein. Then

$$\begin{aligned}\dot{Q}^{(\mathrm{TxTl})}(t) = &\,\mathsf{E}[(-\Delta\mu^{(\mathrm{Tx})} - \Delta g^{(\mathrm{Tx})})\lambda^{(\mathrm{Tx})}(Z(t))] + \mathsf{E}[(-\Delta\mu^{(\mathrm{Rdeg})} + \Delta g^{(\mathrm{Tx})})\lambda^{(\mathrm{Rdeg})}(Z(t))]\\ &+ \mathsf{E}[(-\Delta\mu^{(\mathrm{Tl})} - \Delta g^{(\mathrm{Tl})})\lambda^{(\mathrm{Tl})}(Z(t))] + \mathsf{E}[(-\Delta\mu^{(\mathrm{Pdeg})} + \Delta g^{(\mathrm{Tl})})\lambda^{(\mathrm{Pdeg})}(Z(t))]\end{aligned}$$

such that with the stationarity conditions $\partial_t\,\mathsf{E}[R(t)] = \mathsf{E}[\lambda^{(\mathrm{Tx})}(Z(t))] - \mathsf{E}[\lambda^{(\mathrm{Rdeg})}(Z(t))] = 0$ and $\partial_t\,\mathsf{E}[P(t)] = \mathsf{E}[\lambda^{(\mathrm{Tl})}(Z(t))] - \mathsf{E}[\lambda^{(\mathrm{Pdeg})}(Z(t))] = 0$ and the system variables at stationarity $Z_\infty = (X_\infty, R_\infty, P_\infty) = \lim_{t\to\infty} Z(t)$ (limit in distribution) we obtain

$$\begin{aligned}\lim_{t\to\infty}\dot{Q}^{(\mathrm{TxTl})}(t) = &\lim_{t\to\infty}\mathsf{E}[(-\Delta\mu^{(\mathrm{Tx})} - \Delta\mu^{(\mathrm{Rdeg})})\lambda^{(\mathrm{Tx})}(Z_\infty)] + \lim_{t\to\infty}\mathsf{E}[(-\Delta\mu^{(\mathrm{Tl})} - \Delta\mu^{(\mathrm{Pdeg})})\lambda^{(\mathrm{Tl})}(Z_\infty)]\\ =&(-\Delta\mu^{(\mathrm{Tx})} - \Delta\mu^{(\mathrm{Rdeg})})\mu_1\pi(\mathrm{ON}) + (-\Delta\mu^{(\mathrm{Tl})} - \Delta\mu^{(\mathrm{Pdeg})})\frac{\mu_1\mu_2}{\delta_1}\pi(\mathrm{ON})\\ =&(-\Delta\mu^{(\mathrm{Tx})} - \Delta\mu^{(\mathrm{Rdeg})})\delta_1\,\mathsf{E}[R_\infty] + (-\Delta\mu^{(\mathrm{Tl})} - \Delta\mu^{(\mathrm{Pdeg})})\delta_2\,\mathsf{E}[P_\infty].\end{aligned}$$

Partioning the chemical potentials into length dependent and independent contributions, we obtain the relation

$$-\Delta\mu^{(\mathrm{Tx})} - \Delta\mu^{(\mathrm{Rdeg})} = e_r\,l_r + \bar{e}_r \qquad -\Delta\mu^{(\mathrm{Tl})} - \Delta\mu^{(\mathrm{Pdeg})} = e_p\,l_p + \bar{e}_p$$

as used in the main part.

In conclusion, the energy dissipation rate (in units of $k_\mathrm{B}\,T\,s^{-1}$) of a single gene at stationarity is given as the sum of the entropy production rate of the promoter and $\beta\dot{Q}^{(\mathrm{TxTl})}(t)$.

#### S3.5 Entropy Production Rate on a Finite State Space

Much of the literature on the entropy production rate uses a description based on the infinitesimal generator matrix $\mathbf{\Lambda}$ of a CTMC instead of a reaction network description (*1*, *2*, *9*). A connection between the two pictures is established by using a one-hot coding of the reaction network. In our case, the conversion is employed to make use of Schnakenberg's method (*9*) to compute the entropy production rate of the promoter.

Let $Z$ be N-dimensional with a finite state space $\mathcal{Z} = \{e_1, \ldots, e_N\} \subset \{0,1\}^N$, where $e_i$ is the unit vector with a one at position $i$. Further, the set of reactions be $\{(i,j) \in \mathbb{N}^2 \cap [1,N]^2 : j > i\}$ with associated stoichiometric change vector $\nu_{(i,j)} = e_j - e_i$. For $i < j$ we define the reversible conversion reaction

$$\mathcal{R}_{(i,j)}: \quad Z_i \underset{\lambda^-_{(i,j)}}{\overset{\lambda^+_{(i,j)}}{\rightleftharpoons}} Z_j \quad \text{with} \quad \lambda^+_{(i,j)}(z) = \Lambda_{ij}\,\mathbf{1}(z = e_i), \qquad \lambda^-_{(i,j)}(z) = \Lambda_{ji}\,\mathbf{1}(z = e_j).$$

In the context of an open CRN, this definition implies that each reaction $\mathcal{R}_{(i,j)}$ is associated with a unique change vector of chemostat species. Plugging the definition of the rates into (S16) we obtain

$$\frac{\mathrm{d}}{\mathrm{d}t} H^{\mathrm{tot}}(t) = \sum_{i=1}^{N} \sum_{(k,j):\,k<j} \ln\left( \frac{p(e_i,t)\,\Lambda_{kj}\,\mathbf{1}(e_i = e_k)}{p(e_i + e_j - e_k, t)\,\Lambda_{jk}\,\mathbf{1}(e_i + e_j - e_k = e_j)} \right)$$
$$\cdot \left( \Lambda_{kj}\,\mathbf{1}(e_i = e_k) p(e_i,t) - \Lambda_{jk}\,\mathbf{1}(e_i + e_j - e_k = e_j) p(e_i + e_j - e_k, t) \right),$$

which simplifies to

$$\frac{\mathrm{d}}{\mathrm{d}t} H^{\mathrm{tot}}(t) = \sum_{i=1}^{N} \sum_{j:\,i<j} \ln\left( \frac{p(e_i,t)\,\Lambda_{ij}}{p(e_j,t)\,\Lambda_{ji}} \right) \left( \Lambda_{ij}\, p(e_i,t) - \Lambda_{ji}\, p(e_j,t) \right).$$

### S4 Schnakenberg's Cycle Representation of the Entropy Production Rate

After the derivation of the entropy production rate and it's meaning in the context of open chemical reaction networks, we here want to present the method of Schnakenberg (*9*) to derive the entropy production rate in the steady state. In particular, we want to introduce the expression

$$e_p = \sum_{o \in \mathcal{O}} F(o)\, A(o) \tag{S17}$$

and how to efficiently evaluate it.

While Equation (S17) appears to be more complex then the previous definition of $e_p$, the approach of Schnakenberg provides beautiful aspects for the practical applicability and interpretation of the entropy production rate. As so, we refer the interested reader to (*9*), and especially Sections VIII and IX thereof, for the derivation by Schnakenberg and (*11*) for a more extensive reproduction than we provide here.

The central aspect of Schnakenberg's treatment is the consideration of cycles within the CTMC's state space. In particular, a cycle $o$ is a sequence of reversible reactions connecting states, such that starting at any of these states and following the reactions in the forward direction, one reaches the initial state again. Within this graph centric approach, (*9*) obtains a so called fundamental set of cycles $\mathcal{O}$ by employing a spanning tree based approach. Starting with an arbitrary spanning tree $T$ of the state space graph, the edges, respectively reaction pairs, not included in $T$ are referenced by $\zeta$ and denoted as chords. Adding the arbitrary chord $\varsigma \in \zeta$ to $T$ yields the cycle $o_\varsigma$, from which we remove all edges not contributing to the circle. The fundamental set is obtained by setting $\mathcal{O} = \{o_\varsigma : \varsigma \in \zeta\}$. In addition, (*9*) fixes for each reaction pair $m$ and cycle $o_\varsigma$ a sense of orientation and introduces

$$\phi_m(o_\varsigma) = \begin{cases} 1 & \text{if } m \text{ is parallel to } o_\varsigma, \\ -1 & \text{if } m \text{ is antiparallel to } o_\varsigma, \\ 0 & \text{if } m \text{ is not in } o_\varsigma \end{cases}$$

which gives rise to the relative orientation of $m$ with respect to $o_\varsigma$.

Next, we define for each cycle $o_\varsigma$, the corresponding (probability) flux $F(o_\varsigma)$ and affinity, respectively force, $A(o_\varsigma)$. To simplify notation, we identify the index $i$ with the state $z$ and index $j$ with state $z + \nu_m$, which is reached after the reaction reaction $m$ happened. The quantities of interest are then defined as

$$A(o_\varsigma) = \sum_{m \in o_\varsigma} \phi_m(o_\varsigma)\, A_m \qquad A_m = \log\left(\frac{\pi_i\, \Lambda_{ij}}{\pi_j\, \Lambda_{ji}}\right) \tag{S18}$$

$$F(o_\varsigma) = \phi_\varsigma(o_\varsigma)\, \bar{J}_\varsigma \qquad \bar{J}_\varsigma = \pi_i\, \Lambda_{ij} - \pi_j\, \Lambda_{ji}. \tag{S19}$$

Following (9), the entropy production rate is defined as

$$e_p = \sum_{o_\varsigma \in \mathcal{O}} F(o_\varsigma)\, A(o_\varsigma) \tag{S20}$$

while the applicability of Kirchhoff's Current Law, allowing the use of $\bar{J}_\varsigma$, limits this expression to the steady state.

#### S4.1 Entropy Production Rate of the Eight State Promoter

Within this section, we want to show the relation between entropy production rate and energy dissipation rate at the example of the eight state promoter model being the basis for our technology mapping. In particular, we show the equation

$$e_p = F(o_1)\,\beta\,\mu(o_1) + F(o_2)\,\beta\,\mu(o_2) + F(o_3)\,\beta\,\mu(o_3)$$

where $J_o$ is the net flux along cycle $o$ and $\beta\,\mu_o$ is the corresponding driving force.

Following Schnakenberg (*9*), we start with the state space as being depicted in Figure S2**A**. For this particular structure, Kirchhoff's matrix tree theorem tells us that 56 spanning trees exist and we present our exemplary choice $T$ in Figure S2**B**. The corresponding chords $\zeta = \{\varsigma_1, \varsigma_2, \varsigma_3\}$ are given by the edges $\varsigma_1 = (z_5, z_6)$, $\varsigma_2 = (z_6, z_7)$, and $\varsigma_3 = (z_7, z_8)$. Next, we obtain the cycle $o_i \in \mathcal{O}$ by joining the chord $\varsigma_i \in \zeta$ with the tree T and removing all edges not constituting the resulting loop. At the example of $o_1$, we add $\varsigma_1 = (z_5, z_6)$ to $T$, yielding the cycle over states $z_5$, $z_1$, $z_2$, and $z_6$ to state $z_5$ again. States $z_3$, $z_4$, $z_7$ and $z_8$ are not part of any cycle and so aren't the edges associated. Figure S2**C** presents the cycle $o_1$ obtained by removing the edges and states not included in the cycle. The same approach applies to $o_2$ and $o_3$, where Figures S2**D** and **E** present the corresponding cycles. We complete this step by assigning to each cycle the orientation presented in the respective figure (see Figures S2**C-E**).

To derive the entropy production rate, we first have to determine the thermodynamic forces and fluxes associated to the cycles. For this, we make use of Equations (S18) and (S19). Considering the reaction pairs in the defined orientation gives

$$F(o_1) = \pi_6\,\Lambda_{65} - \pi_5\,\Lambda_{56} \qquad A(o_1) = \ln\left(\frac{\Lambda_{12}\,\Lambda_{26}\,\Lambda_{65}\,\Lambda_{51}}{\Lambda_{21}\,\Lambda_{62}\,\Lambda_{56}\,\Lambda_{15}}\right)$$

$$F(o_2) = \pi_7\,\Lambda_{76} - \pi_6\,\Lambda_{67} \qquad A(o_2) = \ln\left(\frac{\Lambda_{23}\,\Lambda_{37}\,\Lambda_{76}\,\Lambda_{62}}{\Lambda_{32}\,\Lambda_{73}\,\Lambda_{67}\,\Lambda_{26}}\right)$$

$$F(o_3) = \pi_8\,\Lambda_{87} - \pi_7\,\Lambda_{78} \qquad A(o_3) = \ln\left(\frac{\Lambda_{34}\,\Lambda_{48}\,\Lambda_{87}\,\Lambda_{73}}{\Lambda_{43}\,\Lambda_{84}\,\Lambda_{78}\,\Lambda_{37}}\right)$$

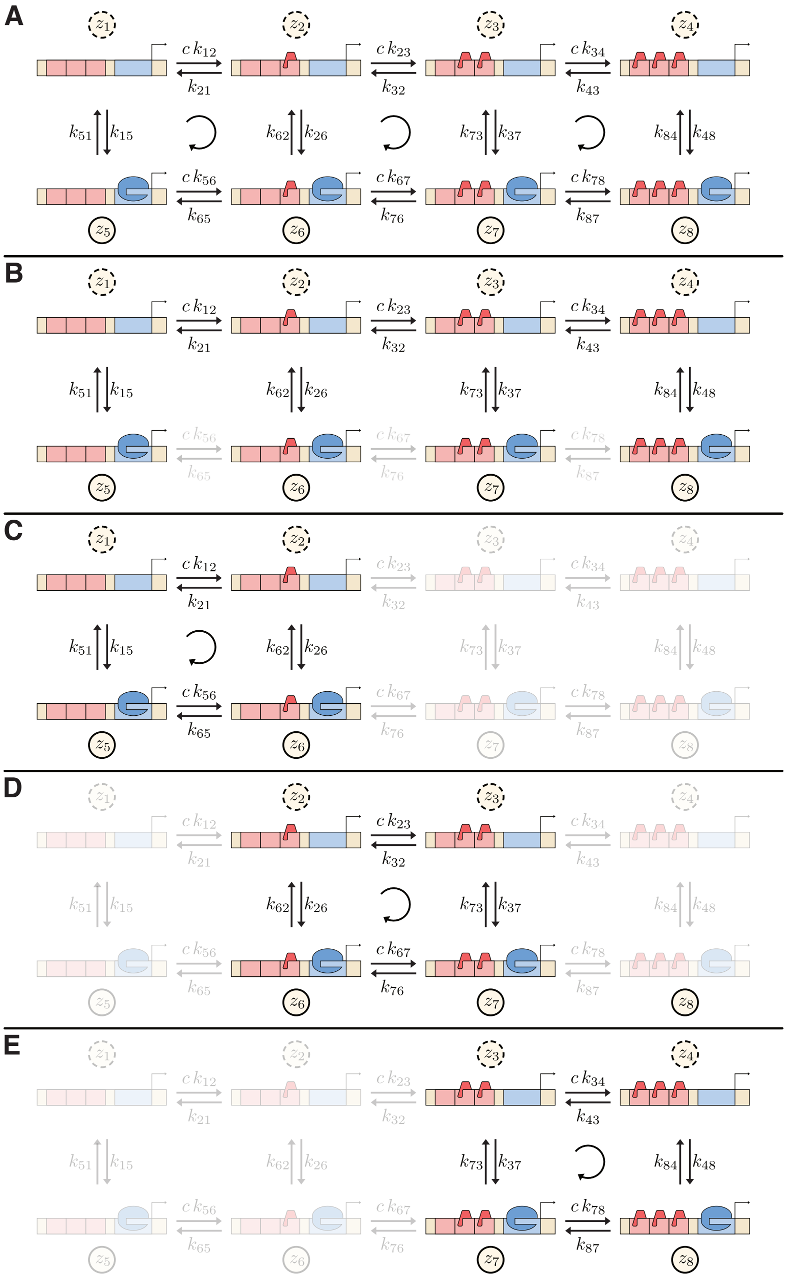


Figure S2: **Derivation of entropy production rate following Schnakenberg (9). A:** The state space of the CTMC under concideration. **B:** One of the spanning trees, whereby the greyed out reaction pairs are the excluded transitions, constituting the set of chords. **C-E:** The cycles obtained by joining the tree with each of the chords and removing all other states and reactions (greyed out).

where $F(o_i)$ directly follows from (S18) and $A(o_i)$ already includes the simplification

$$
\begin{aligned}
A(o_i) &= \sum_{(j_1,j_2)\in o_i} \phi_m(o_i)\, \ln\left(\frac{\pi_{j_1}\,\Lambda_{j_1j_2}}{\pi_{j_2}\,\Lambda_{j_2j_1}}\right) \\
&= \ln\left(\prod_{(j_1,j_2)\in o_i}\left(\frac{\pi_{j_1}\,\Lambda_{j_1j_2}}{\pi_{j_2}\,\Lambda_{j_2j_1}}\right)^{\phi_m(o_i)}\right) \\
&= \ln\left(\frac{\pi_{j_1}\,\Lambda_{j_1j_2}}{\pi_{j_2}\,\Lambda_{j_2j_1}}\,\frac{\pi_{j_2}\,\Lambda_{j_2j_3}}{\pi_{j_3}\,\Lambda_{j_3j_2}}\,\frac{\pi_{j_3}\,\Lambda_{j_3j_4}}{\pi_{j_4}\,\Lambda_{j_4j_3}}\,\frac{\pi_{j_4}\,\Lambda_{j_4j_1}}{\pi_{j_1}\,\Lambda_{j_1j_4}}\right) \\
&= \ln\left(\frac{\Lambda_{j_1j_2}\,\Lambda_{j_2j_3}\,\Lambda_{j_3j_4}\,\Lambda_{j_4j_1}}{\Lambda_{j_2j_1}\,\Lambda_{j_3j_2}\,\Lambda_{j_4j_3}\,\Lambda_{j_1j_4}}\right)
\end{aligned}
$$

which makes use of the cycles being closed and where we assumed that the state indices $j$ are ordered in the direction of the cycle. As so, the affinity or thermodynamic force of a cycle explicitly solely depends on the propensities and is independent of the actual state probabilities.

Inserting the previous into Equation (S20), we obtain the entropy production rate.

$$
\begin{aligned}
e_p &= \sum_{o_i\in\mathcal{O}} F(o_i)\,A(o_i) \\
&= \quad (\pi_6\,\Lambda_{65} - \pi_5\,\Lambda_{56})\,\ln\left(\frac{\Lambda_{12}\,\Lambda_{26}\,\Lambda_{65}\,\Lambda_{51}}{\Lambda_{21}\,\Lambda_{62}\,\Lambda_{56}\,\Lambda_{15}}\right) \\
&\quad + (\pi_7\,\Lambda_{76} - \pi_6\,\Lambda_{67})\,\ln\left(\frac{\Lambda_{23}\,\Lambda_{37}\,\Lambda_{76}\,\Lambda_{62}}{\Lambda_{32}\,\Lambda_{73}\,\Lambda_{67}\,\Lambda_{26}}\right) \\
&\quad + (\pi_8\,\Lambda_{87} - \pi_7\,\Lambda_{78})\,\ln\left(\frac{\Lambda_{34}\,\Lambda_{48}\,\Lambda_{87}\,\Lambda_{73}}{\Lambda_{43}\,\Lambda_{84}\,\Lambda_{78}\,\Lambda_{37}}\right)
\end{aligned}
$$

By recalling the thermodynamic consistency relation for open chemical reaction networks, we can simplify this expression further. Using the current notation for Equation (S12) for reasons of brevity, the relation is

$$
\begin{aligned}
\frac{\Lambda_{ij}}{\Lambda_{ji}} &= \exp\left(-\beta\left(\Delta g_{ij} - \Delta\mu_{ij}\right)\right) \\
\Delta g_{ij} &= g_j - g_i = -\Delta g_{ji}
\end{aligned}
$$

where $g_i$ is the Gibbs free energy of the state $z_i$ and $\Delta\mu_{ij}$ is the chemical potential for transitioning from state $z_i$ to state $z_j$. For a closed loop $o$ in the state space of our CTMC, the forward and backward propensities satisfy

$$\prod_{(i,j)\in o} \frac{\Lambda_{ij}}{\Lambda_{ji}} = \exp\left(\beta\,\mu(o)\right) \qquad \text{with} \qquad \mu(o) = \sum_{(i,j)\in o} \mu_{ij}$$

which follows directly by inserting the thermodynamic consistency relation into the left side. For the entropy production rate, this yields

$$e_p = F(o_1)\,\beta\,\mu(o_1) + F(o_2)\,\beta\,\mu(o_2) + F(o_3)\,\beta\,\mu(o_3)$$

which is the entropy production rate expressed in terms of the chemical potential and the flux along the CTMC's cycles. In this context, we raise the reader's attention to the dependency on the cognate transcription factor's concentration. Despite not explicitly stated, the propensities $\Lambda_{ij}$ depend on $c$ and it's influence is covered by the chemical potentials $\mu(o_i)$.

### S5 ARCTIC and Energy Aware Technology Mapping

ARCTIC is a technology mapping framework first introduced in (*12*) and extended in (*13*) as well as in this work. Given a genetic gate library and the specification of the Boolean function, the framework provides methods to efficiently synthesize circuits for the desired Boolean function. As a unique feature, it considers structural variants of circuits as candidate topologies for gate assignment optimization. With simulated annealing, Branch-and-Bound, and exhaustive enumeration, three optimization algorithms are available for the gate assignment. Exhaustive enumeration is the traditional approach of evaluating all valid assignments of genetic gates and is in turn computationally expensive. Branch-and-Bound performs an implicit enumeration, providing a beneficial performance and can also guarantee exactness. However, the method requires well adapted models providing guarantees on monotonicity and

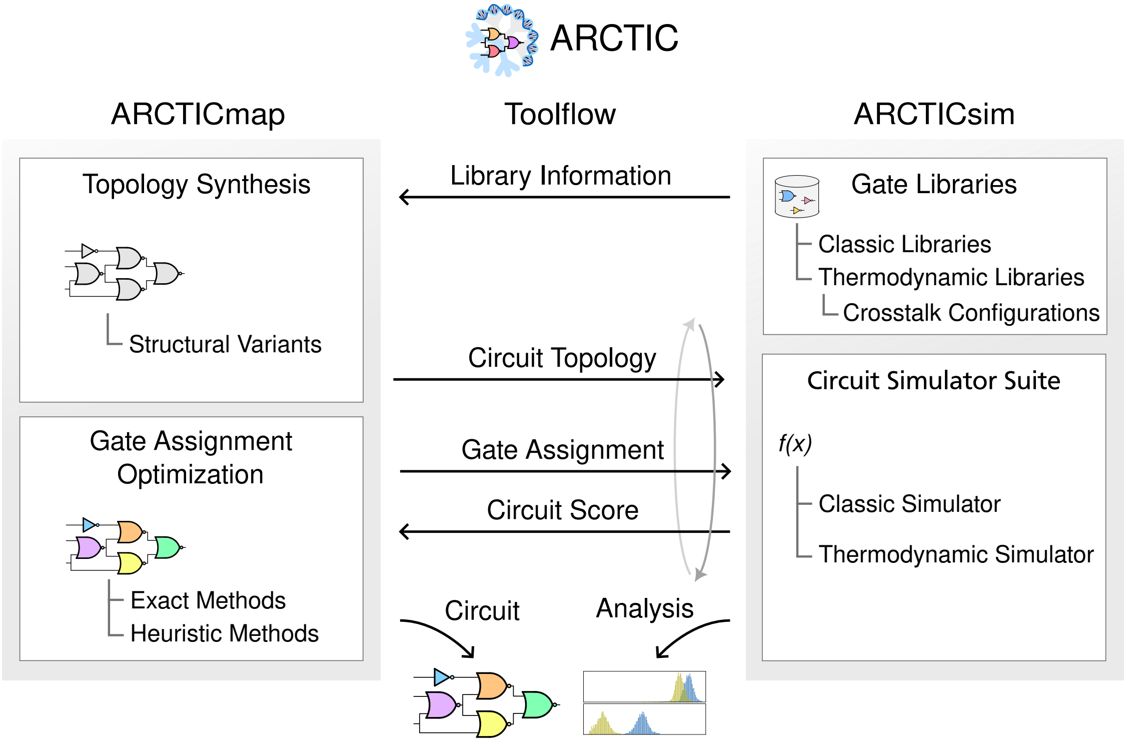


Figure S3: **Overview of the technology mapping framework ARCTIC.**

boundedness of gate transfer functions. Simulated annealing is the most versatile method, as it is a flexible heuristic. It allows for optimizing arbitrary objective functions and extended by context information (*12*), it provides an adequate performance.

The mapping part is complemented by genetic circuit simulators putting a special focus on robustness and context effects. To this end, ARCTIC includes with the E-Score (*12*) (see Methods of main part) a scoring method accounting for output distributions. This is utilized by the particle based simulation used in this work and allows to evaluate the circuit's performance in the presence of cell-to-cell variability and intrinsic noise. Besides, (*13*) presents a thermodynamic equilibrium model accounting for context effects such as cross-talk among gates, titration of transcription factors to non-cognate binding sites, and interference of host molecules with the circuit.

As the **Energy Aware Technology Mapping** makes use of the model presented in this work, we subsequently describe the creation of the genetic gate library used for the present evaluation.

#### S5.1 Library Creation

In order to highlight the adaptability to different scenarios and already existing libraries, we use the user constraint file (UCF) of Chen et al. (*14*) as starting point for the creation of our genetic gate library. Providing all the data relevant to Cello (*14*–*16*), the UCF includes information such as cytometry data, toxicity, DNA sequences, and Hill curve parameters for genetic logic gates implemented and evaluated in the yeast Saccharomyces cerevisiae (S. cerevisiae). The first step in library creation is to translate the entries within the UCF to entries compatible with our framework. This mainly involves providing new identifiers, reshaping entries and stripping of the information currently of no use to us, such as the `motif_library` collection. In the following paragraphs, we consider the most important collections further and provide the information relevant for intercomparability.

**Device** The central aspect of our library is the device. It represents the actual genetic logic gate, which is constructed of a promoter, a protein, and the UTR, respectively Kozak sequence. As all the previously mentioned parts are included indivdually, the device collection holds a reference to the parts of relevance. In addition, it gives rise to the logic gate primitives realizable, which are in our case NOT and NOR. To identify which gates can be used jointly in a circuit, the group tag indicates the transcription factor used. By this, the technology mapping algorithm can exclude gate assignments featuring a single group more than once.

**Promoter** The promoter entry includes all the entries relevant for the characterization of the promoter model employed. In particular, these are the cognate transcription factors, references to the corresponding DNA sequences, information for the proximity based gate assignment (*12*), and the actual parameters of the model. These include the transcriptional activities of all promoter states and the reaction rate constants obtained through parameter estimation. In the following, we present the prerequisites for this process.

A key aspect of the Cello UCF is, that the values provided are in units of RPU or related to it. RPU stands for relative promoter unit and relates the strength of a promoter to a reference promoter (*14*, *15*, *17*). In comparison, the model presented in this work is defined with protein abundance in mind. However, considering the derivation of the values presented in the UCF, one observes that these relate to the fluorescence intensity of some reporter protein and are translated to RPU by assuming that the promoter activity is proportional to the protein expression level (*15*, *17*). In the model's parameter estimation, we partially adapt this view and assume, that fluorescence intensity is proportional to the expression level. Due to the significant mismatch between the order of magnitude of RPU and protein concentration, we rescale the protein abundance in dependence to the values of $\mu_1$, $\mu_2$, $\delta_1$, and $\delta_2$. In particular, the rescaled protein level $\tilde{P}$ is

$$\tilde{P} = P \, \frac{\delta_1 \, \delta_2}{\mu_1 \, \mu_2} \tag{S21}$$

where $P$ is the actual protein abundance. Equation (S21) eliminates the concern with respect to the order of magnitude and at the same time preserves the coefficient of variation of the associated distribution. As so, there is no loss of probabilistic information while obtaining a representation compatible with RPU.

Figure S4 reassembles the setting of genetic gate characterization, respectively the one employed for the measurement of the cytometry data, at the example of the transcription factor phlF1. The parameter estimation approach reassembles this setting to derive the parameters of the promoter model. By the previously introduced rescaling, the transcriptional activity parameters $a_i$ are in the range of RPU, making the models response characteristic comparable to the models provided in (*14*).

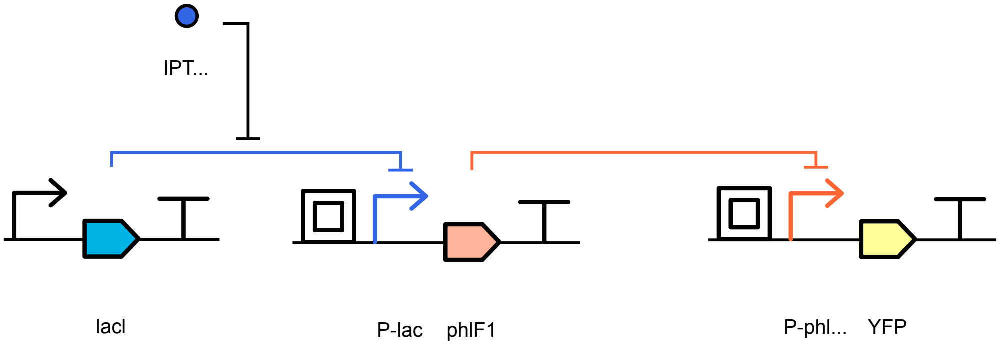


Figure S4: **Genetic gate characterization.** This figure presents the gene circuit for the characterization of the genetic logic gates. The setup consist of a constitutive promoter expressing the protein lacI, which can act as transcription factor or bind to the inducer IPTG. This transcription factor represses it's cognate promoter, which realizes the input to the gate under consideration, here P1_PhlF. The expression level of the reporter protein succeeding the gate is in practice measured via flow cytometry and thereafter normalized to RPU (*15*).

**Protein** The protein collection provides for each transcription factor the information relevant for the model. In detail, these are the rates for the RNA and protein dynamics and the information on the energetics. As introduced subsequently, we make use of the following rates during library creation of S. cerevisiae. The transcription rate is $\mu_1 = 6.\bar{6}\,10^{-3}\,s^{-1}$, the translation rate is $\mu_2 = 55.\bar{5}\,10^{-3}\,s^{-1}$, and the corresponding degradation rates are $\delta_1 = 0.58\,10^{-3}\,s^{-1}$ and $\delta_2 = 0.29\,10^{-3}\,s^{-1}$. Furthermore, we derive from the DNA sequences encoding the proteins the RNA and protein lengths $l_r$ and $l_p$. For the RNA length $l_r$, we additionally take the provided UTR into account, as it is part of the transcript but not of the protein the DNA sequence encodes for. The protein length $l_p$ in amino acids is then the sequence length divided by the codon size (i.e. three). In combination with the parameters $e_r = 16$, $\bar{e}_r = 0$, $e_p = 42$, and $\bar{e}_p = 52$, this provides all the information relevant to determine the energetic cost of a single RNA and protein molecule.

As the previous rates are host dependent, they are only applicable for S. cerevisae and need to be adapted for other organisms such as E. coli.

#### S5.2 Parameter Estimation

The parameter estimation approach for our particular promoter model instantiation has to identify the values of the 22 parameters matching best to the provided data. The first step is to reimplement the experimental setting used to characterize the parts in (*14*, *15*) *in silico*. This setup is shown in Figure S4 and consists of two genes, whereby the genetic gate is realized by the transcription factor encoded by the first gene and the promoter on the second gene. In comparison to (*14*), we omit the gene constitutively expressing lacI, the transcription factor binding to the inducer IPTG and allowing for induction, by replacing it *in silico* with a degenerate promoter directly translating inducer levels to promoter activities. The required information is not included in the UCF but is provided as supplementary data to the work of Chen et al. (*14*). Adding the reporter protein to the end of the gene cascade, we follow the setting of (*14*). Next, the inducer levels are associated to the corresponding cytometry measurements, respectively to the associated mean and variance.

The parameter matching is achieved by minimizing the error function measuring the difference between the distribution resulting from the model setup and the data provided. As the cytometry data is given in terms of RPU, we rescale the reporter output to promoter activity levels, yielding the model's output in RPU. This is described by SI Equation (S21). The error function measures the logarithmic distance between the model's mean and standard deviation to the reference data. To account for precisely matching the saturated regions, we weight the errors at the smallest and largest input conditions accordingly. In addition, a penalization is added in case the current response characteristic is not monotonous, which ensures well behaving models in domains not covered by the experimental data.

To ease optimization, the reaction rate constants for the reactions between to transcriptional activity levels are initialized to feature an inverter characteristic, while the others are chosen randomly. Furthermore, we constrain all rate constants to the range $[10^{-5}, 10^{5}]$ and the activity level of the active levels to be at least the maximum mean value encountered in the cytometry data and exceeding it by factor two at most. The inactive level is upper bounded by the smallest mean value and is allowed to undercut it by half. For each genetic gate, the best parameter set out of ten optimizations is chosen and included in the respective promoter entry in the gate library.

### S6 Relative Promoter Units, Relative Expression Units, and the Characterization of Genetic Logic Gates in Cello

Relative Promoter Units (RPU) are commonly used in synthetic biology as they enable the comparison of promoters in different contexts and contribute to the standardization of the field (*18*, *19*). Nielsen et al. (*15*) and Chen et al. (*14*) use relative promoter units (RPU) to characterize their gates and sensors. The RPUs are derived from the flow cytometry measurements of the constructs and by normalizing with respect to previously obtained values. The formula is

$$RPU = \frac{YFP - YFP_0}{YFP_{RPU} - YFP_0}$$

where the median fluorescence of the construct $YFP$, the RPU standard plasmid $YFP_{RPU}$, and the autofluorescence $YFP_0$ are set in relation. Within (*17*), a predecessor of (*15*) already introducing the genetic gates, the concept relative expression units (REU) is used to refer to the values later concidered as RPU.

Figure S4 visualizes an exemplary plasmid for the characterization of a gate's reponse characteristic in RPU, following the description of Nielsen et al. (*20*) which extends on Stanton et al. (*17*) and matches Chen et al. (*14*). Both, the input and output of a gate are characterized in relative promoter units (RPU). The output RPU are derived from the flow cytometry measurements of YFP given distinct ligand concentrations and the conversion to RPU described previously. To complete the gate description, the ligand concentration needs to be replaced by the input promoter activity. This is done by expressing YFP directly by the promoter sensitive to the transcription factor capable of binding to the ligand. The result is the expression level depending on the ligand concentration, from which one can derive the respective input RPU. By associating the input RPU to the output RPU of the gate, the complete gate description is obtained. Figure (*21*) in the supplement of Nielsen et al. showcases this visually.

### S7 Gene Expression Kinetics in *S. cerevisiae*

In this section, we present key rate constants for transcription, translation, and degradation processes in *S. cerevisiae*. Most values are derived from a combination of mechanistic understanding of biochemical reactions on one hand (*22*), and experimental estimates on the other (*23*).

Table S1: Transcription, translation, and degradation rate constants for *S. cerevisiae*.

| **Rate constant** | **C** | **Unit** | **Value** |
|---|---|---|---|
| Cell volume | $k$ | $\mu m^3$ | 0.4 – 2.5 (*23*) |
| Transcription rate | $\tilde{\lambda}_1$ | $nt.s^{-1}$ | 40 (*24*) |
| Translation rate | $\tilde{\lambda}_2$ | $aa.s^{-1}$ | 3 – 10 (*23*) |
| Transcription rate | $\lambda_1$ | $mRNA.h^{-1}$ | 2 – 30 (*25*) |
| Translation efficiency | $\lambda_2$ | $protein.mRNA^{-1}.h^{-1}$ | 20 – 2000 (*26*) |
| mRNA half life | $t_{m_{1/2}}$ | $min$ | 20 (*23*) |
| Protein | $t_{p_{1/2}}$ | $min$ | 40 (*23*) |
| mRNA degradation rate | $\delta_m$ | $s^{-1}$ | $5.8 \cdot 10^{-4}$ (*23*, *27*) |
| protein degradation rate | $\delta_p$ | $s^{-1}$ | $2.9 \cdot 10^{-4}$ (*23*) |

#### S7.1 Transcription Rates and Energy Dependence

Transcription is catalyzed by a single enzyme, RNA polymerase, which derives its energy from the hydrolysis of nucleoside triphosphates (NTPs), the substrates for RNA synthesis.

$$NTP + RNA_n \rightarrow PPi + RNA_{n+1}$$

Although the initiation and termination phases require additional energy, the overall cost of transcription is predominantly governed by the elongation phase, where each nucleotide addition consumes one NTP.

**Transcription Rates** For our gene expression model, the transcription rate in mRNA per hour ($mRNA \cdot h^{-1}$) is relevant as it directly indicates the gene expression levels. According to (25), the mRNA transcription rate in *S. cerevisiae* is approximately 2 to 30 $mRNA \cdot h^{-1}$. For our simulations, we chose a translation rate value of 200 $protein \cdot mRNA^{-1} \cdot h^{-1}$

#### S7.2 Translation Rates and Energy Dependence

**Energy Dependence** Elongation is the step that consumes the most energy during the entire protein synthesis process. The process is well-studied, allowing each reaction to be enumerated for an accurate estimation of the cost (22, 28).

- Aminoacyl-tRNA charging:

$$\text{AA} + \text{ATP} \rightarrow \text{AA-AMP} + \text{PPi}$$
$$\text{AA-AMP} + \text{tRNA} \rightarrow \text{AA-tRNA} + \text{AMP}$$

- EF-Tu hydrolyzes one GTP molecule into a GDP and is used to load the aminoacyl-tRNA in the A site of the ribosome.
- Then, the translocation of tRNAs and mRNA hydrolyzes one GTP into a GDP.

Initiation and termination processes in translation also require energy. The search for the first AUG codon involves ATP hydrolysis into ADP and inorganic phosphate (Pi) (1 to 4 ATP molecules per initiation event). During initiation, an initiation factor hydrolyzes one GTP molecule into GDP and Pi. In termination, release factors hydrolyze one GTP molecule into GDP and Pi (*22*).

**Translation Rates** For our gene expression model, the translation rate in proteins per hour ($protein \cdot mRNA^{-1} \cdot h^{-1}$) is relevant as it directly indicates the protein synthesis levels. Reported values for *S. cerevisiae* range from approximately 20 to 2000 $protein \cdot mRNA^{-1} \cdot h^{-1}$ (*26*), with a maximum around 4000 $protein \cdot mRNA^{-1} \cdot h^{-1}$ (*29*). For our simulations, we chose a translation rate value of 200 $protein \cdot mRNA^{-1} \cdot h^{-1}$.

#### S7.3 Degradation and Dilution Rates

**RNA Degradation** A few enzymes that participate in RNA degradation require ATP, but most do not, and this energy requirement can often be neglected (*30*).

**RNA Degradation Rate** Degradation rates are often represented with half-life times, using the conversion: $\delta_1 = \ln(2)/t_{1/2}$. For RNA degradation rates in *S. cerevisiae*, $t_{1/2} \approx$ 20 min so $\delta_1 \approx 5.8 \cdot 10^{-4}$ s$^{-1}$ (*23*, *27*).

**Protein Degradation** In contrast to RNA degradation, protein degradation requires external energy in the form of ATP. Specifically, the process of polyubiquitylation and subsequent degradation by the proteasome involve ATP-dependent reactions. Estimates suggest that the degradation of a single protein can require the hydrolysis of over 100 to a few hundred molecules of ATP (*31*). Since it depends on the protein, and that dilution mostly make the protein decay, we approximate the cost to zero.

**Protein Degradation Rate** For protein degradation in *S. cerevisiae* we have $t_{1/2} \approx 40$ min therefore $\delta_2 \approx 2.9 \cdot 10^{-4}$ s$^{-1}$ (*23*).

**Dilution Rates** The previous decay rate, include both active degradation, and dilution. The most important effect of the decrease of concentration of a protein, is the dilution, caused by the growth of the microorganism. We have, either for mRNA, or protein

$$\delta = \delta_{degradation} + \delta_{dilution}$$

For the dilution rate, it is directly linked with the doubling time:

$$\delta_{dilution} = 1/\tau$$

with $\delta$ being the dilution rate, and $\tau$ being the time interval of the cell cycle. For *S. cerevisiae*, the doubling time is about 90 minutes, so $\delta_{dilution} = 1.85 \times 10^{-4}\ s^{-1}$ (*23*).

### S8 Extended Figures and Tables

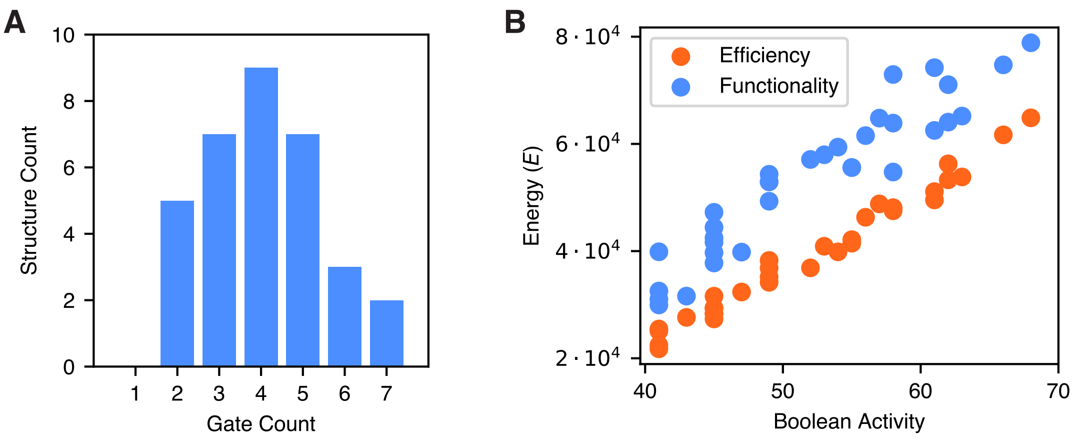


Figure S5: **Characteristics of benchmark circuits. A:** The gate count distribution of the circuits within the benchmark set. **B:** Comparison of Boolean activity for energy efficiency and functionally optimized circuits.

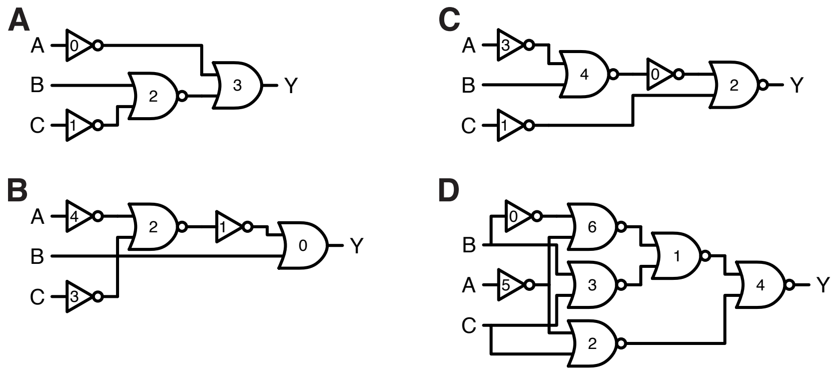


Figure S6: **Circuit structures. A:** 0x2F. **B:** 0xDF. **C:** 0x20. **D:** 0x81. Each gate is labelled with its ID.

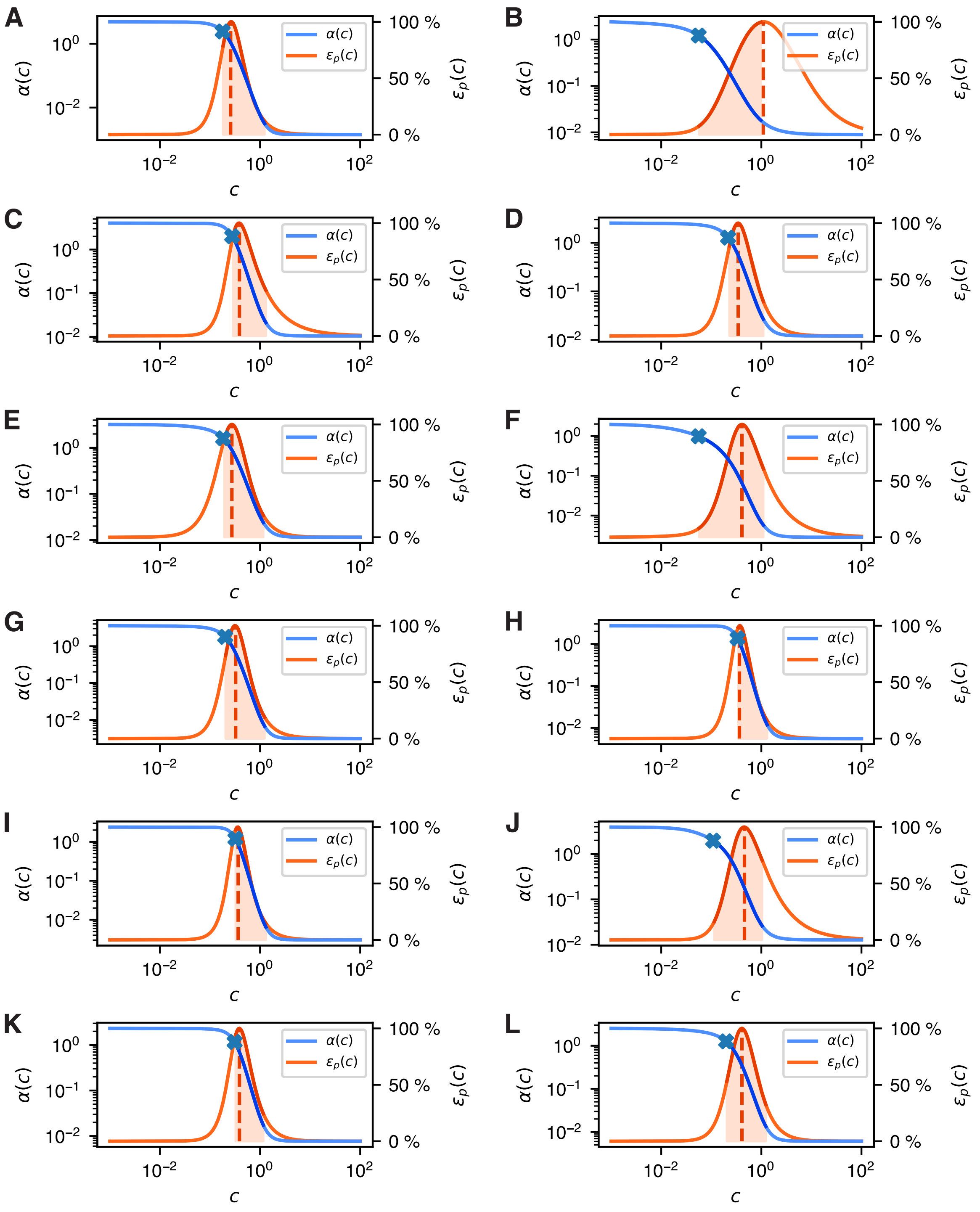


Figure S7: **Entropy production rate $\epsilon_p(c)$ and average promoter activity $\alpha(c)$ as functions of transcription factor abundance.** **A:** P1_BM3RI **B:** P1_CI **C:** P1_CI434 **D:** P1_HKCI **E:** P1_IcaR **F:** P1_LexA **G:** P1_PhlF **H:** P1_PsrA **I:** P1_QacR **J:** P2_CI **K:** P2_CI434 **L:** P2_LexA

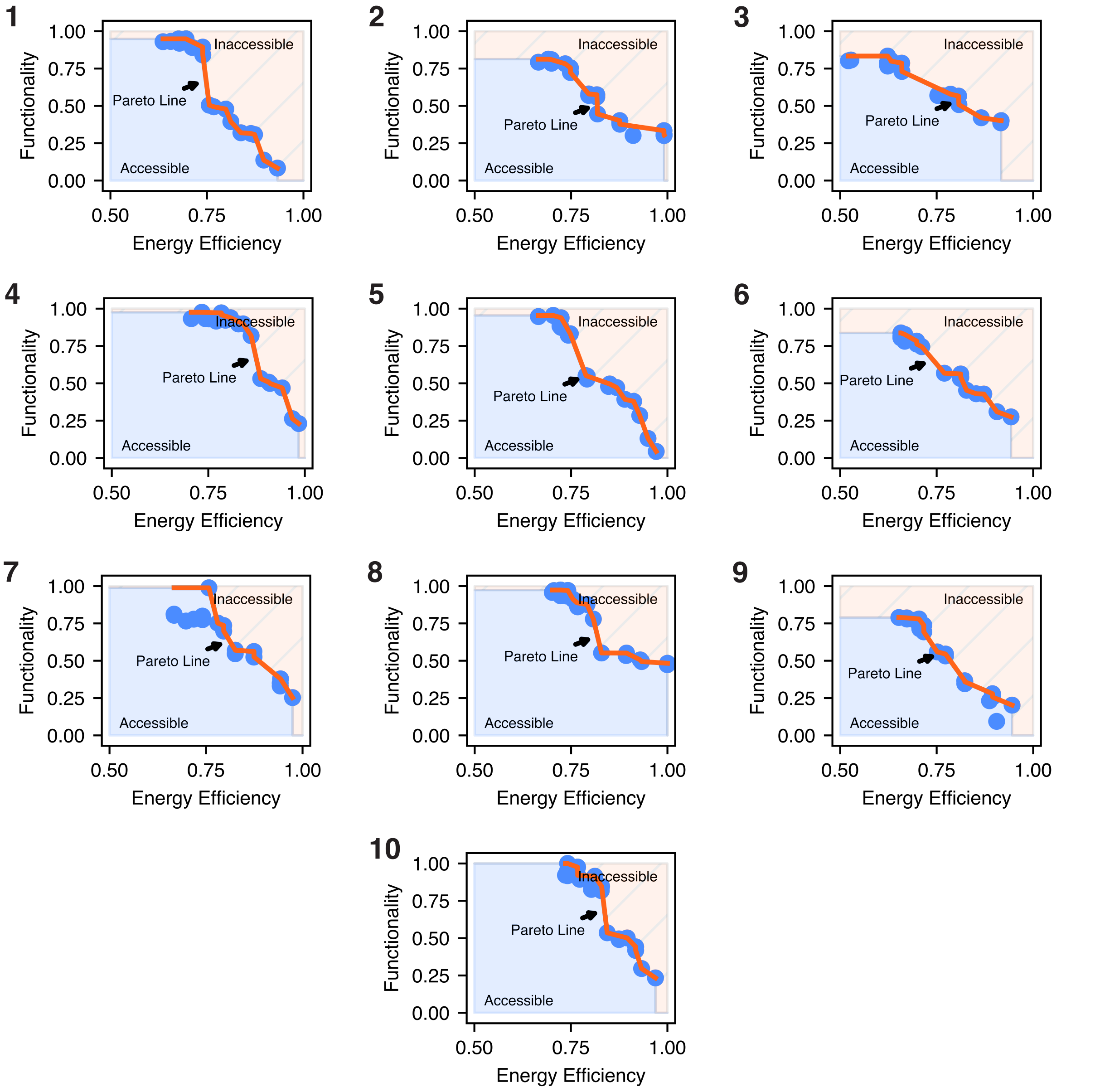


Figure S8: **Pareto fronts of structural variants of circuit 0xF7.** The structure IDs are indicated by the bold numbers in the figure.

Table S2: Energy per gate for all Boolean assignments for circuit 0x2F (Energy opt.) and mean values in italic. Figure S6**A** presents the corresponding structure.

| A | B | C | NOR2_2 | NOT_0 | NOT_1 | OR2_3 |
|---|---|---|---|---|---|---|
| 0.00 | 0.00 | 0.00 | 12544.80 | 63.67 | 18.02 | 8375.74 |
| 0.00 | 0.00 | 0.00 | 60.62 | 63.67 | 10833.12 | 20563.13 |
| 0.00 | 0.00 | 0.00 | 25810.57 | 63.67 | 18.02 | 8374.46 |
| 0.00 | 0.00 | 0.00 | 13326.38 | 63.67 | 10833.12 | 8375.43 |
| 0.00 | 0.00 | 0.00 | 12544.80 | 7506.98 | 18.02 | 40.17 |
| 0.00 | 0.00 | 0.00 | 60.62 | 7506.98 | 10833.12 | 12227.56 |
| 0.00 | 0.00 | 0.00 | 25810.57 | 7506.98 | 18.02 | 38.89 |
| 0.00 | 0.00 | 0.00 | 13326.38 | 7506.98 | 10833.12 | 39.86 |
| *0.00* | *0.00* | *0.00* | *12935.59* | *3785.32* | *5425.57* | *7254.41* |

Table S3: Energy per gate for all Boolean assignments for circuit 0x2F (Func. opt.) and mean values in italic. Figure S6**A** presents the corresponding structure.

| A | B | C | NOR2_2 | NOT_0 | NOT_1 | OR2_3 |
|---|---|---|---|---|---|---|
| 0.00 | 0.00 | 0.00 | 22249.87 | 11.47 | 66.06 | 18288.26 |
| 0.00 | 0.00 | 0.00 | 135.04 | 11.47 | 7788.78 | 42972.25 |
| 0.00 | 0.00 | 0.00 | 32185.15 | 11.47 | 66.06 | 18288.22 |
| 0.00 | 0.00 | 0.00 | 10070.33 | 11.47 | 7788.78 | 18289.74 |
| 0.00 | 0.00 | 0.00 | 22249.87 | 14076.13 | 66.06 | 24.27 |
| 0.00 | 0.00 | 0.00 | 135.04 | 14076.13 | 7788.78 | 24708.26 |
| 0.00 | 0.00 | 0.00 | 32185.15 | 14076.13 | 66.06 | 24.23 |
| 0.00 | 0.00 | 0.00 | 10070.33 | 14076.13 | 7788.78 | 25.74 |
| *0.00* | *0.00* | *0.00* | *16160.10* | *7043.80* | *3927.42* | *15327.62* |

Table S4: Gate output values for all Boolean assignments for circuit 0x2F (Energy opt.) and mean values in italic. Figure S6**A** presents the corresponding structure.

| A | B | C | NOR2_2 | NOT_0 | NOT_1 | OR2_3 |
|---|---|---|---|---|---|---|
| 0.01 | 0.00 | 0.00 | 0.00 | 1.63 | 2.32 | 1.64 |
| 0.01 | 0.00 | 1.80 | 2.39 | 1.63 | 0.01 | 4.02 |
| 0.01 | 2.46 | 0.00 | 0.00 | 1.63 | 2.32 | 1.64 |
| 0.01 | 2.46 | 1.80 | 0.00 | 1.63 | 0.01 | 1.64 |
| 1.30 | 0.00 | 0.00 | 0.00 | 0.00 | 2.32 | 0.01 |
| 1.30 | 0.00 | 1.80 | 2.39 | 0.00 | 0.01 | 2.39 |
| 1.30 | 2.46 | 0.00 | 0.00 | 0.00 | 2.32 | 0.01 |
| 1.30 | 2.46 | 1.80 | 0.00 | 0.00 | 0.01 | 0.01 |
| *0.65* | *1.23* | *0.90* | *0.60* | *0.82* | *1.16* | *1.42* |

Table S5: Gate output values for all Boolean assignments for circuit 0x2F (Func. opt.) and mean values in italic. Figure S6**A** presents the corresponding structure.

| A | B | C | NOR2_2 | NOT_0 | NOT_1 | OR2_3 |
|---|---|---|---|---|---|---|
| 0.00 | 0.00 | 0.01 | 0.00 | 3.58 | 4.03 | 3.58 |
| 0.00 | 0.00 | 1.30 | 4.83 | 3.58 | 0.02 | 8.41 |
| 0.00 | 1.80 | 0.01 | 0.00 | 3.58 | 4.03 | 3.58 |
| 0.00 | 1.80 | 1.30 | 0.00 | 3.58 | 0.02 | 3.58 |
| 2.46 | 0.00 | 0.01 | 0.00 | 0.00 | 4.03 | 0.00 |
| 2.46 | 0.00 | 1.30 | 4.83 | 0.00 | 0.02 | 4.83 |
| 2.46 | 1.80 | 0.01 | 0.00 | 0.00 | 4.03 | 0.00 |
| 2.46 | 1.80 | 1.30 | 0.00 | 0.00 | 0.02 | 0.01 |
| *1.23* | *0.90* | *0.65* | *1.21* | *1.79* | *2.03* | *3.00* |

Table S6: Energy per gate for all Boolean assignments for circuit 0xDF (Energy opt.) and mean values in italic. Figure S6**B** presents the corresponding structure.

| A | B | C | NOR2_2 | NOT_1 | NOT_3 | NOT_4 | OR2_0 |
|---|---|---|---|---|---|---|---|
| 0.00 | 0.00 | 0.00 | 23940.44 | 66.96 | 74.12 | 16.22 | 8296.52 |
| 0.00 | 0.00 | 0.00 | 12854.61 | 72.30 | 8739.06 | 16.22 | 8182.20 |
| 0.00 | 0.00 | 0.00 | 23940.44 | 66.96 | 74.12 | 16.22 | 20837.56 |
| 0.00 | 0.00 | 0.00 | 12854.61 | 72.30 | 8739.06 | 16.22 | 20723.24 |
| 0.00 | 0.00 | 0.00 | 11183.42 | 75.99 | 74.12 | 9755.99 | 8105.47 |
| 0.00 | 0.00 | 0.00 | 97.58 | 17852.22 | 8739.06 | 9755.99 | 25.41 |
| 0.00 | 0.00 | 0.00 | 11183.42 | 75.99 | 74.12 | 9755.99 | 20646.51 |
| 0.00 | 0.00 | 0.00 | 97.58 | 17852.22 | 8739.06 | 9755.99 | 12566.45 |
| *0.00* | *0.00* | *0.00* | *12019.01* | *4516.87* | *4406.59* | *4886.11* | *12422.92* |

Table S7: Energy per gate for all Boolean assignments for circuit 0xDF (Func. opt.) and mean values in italic. Figure S6**B** presents the corresponding structure.

| A | B | C | NOR2_2 | NOT_1 | NOT_3 | NOT_4 | OR2_0 |
|---|---|---|---|---|---|---|---|
| 0.00 | 0.00 | 0.00 | 25030.36 | 58.76 | 20.05 | 63.67 | 24892.68 |
| 0.00 | 0.00 | 0.00 | 9907.19 | 82.43 | 12057.13 | 63.67 | 24840.65 |
| 0.00 | 0.00 | 0.00 | 25030.36 | 58.76 | 20.05 | 63.67 | 37433.72 |
| 0.00 | 0.00 | 0.00 | 9907.19 | 82.43 | 12057.13 | 63.67 | 37381.69 |
| 0.00 | 0.00 | 0.00 | 15239.64 | 62.91 | 20.05 | 7506.98 | 24883.91 |
| 0.00 | 0.00 | 0.00 | 116.26 | 22235.24 | 12057.13 | 7506.98 | 17.54 |
| 0.00 | 0.00 | 0.00 | 15239.64 | 62.91 | 20.05 | 7506.98 | 37424.95 |
| 0.00 | 0.00 | 0.00 | 116.26 | 22235.24 | 12057.13 | 7506.98 | 12558.58 |
| *0.00* | *0.00* | *0.00* | *12573.36* | *5609.84* | *6038.59* | *3785.32* | *24929.22* |

Table S8: Gate output values for all Boolean assignments for circuit 0xDF (Energy opt.) and mean values in italic. Figure S6**B** presents the corresponding structure.

| A | B | C | NOR2_2 | NOT_1 | NOT_3 | NOT_4 | OR2_0 |
|---|---|---|---|---|---|---|---|
| 0.00 | 0.00 | 0.01 | 0.01 | 1.62 | 2.09 | 2.39 | 1.62 |
| 0.00 | 0.00 | 1.30 | 0.01 | 1.60 | 0.01 | 2.39 | 1.60 |
| 0.00 | 2.46 | 0.01 | 0.01 | 1.62 | 2.09 | 2.39 | 4.08 |
| 0.00 | 2.46 | 1.30 | 0.01 | 1.60 | 0.01 | 2.39 | 4.05 |
| 1.80 | 0.00 | 0.01 | 0.01 | 1.58 | 2.09 | 0.00 | 1.59 |
| 1.80 | 0.00 | 1.30 | 3.08 | 0.00 | 0.01 | 0.00 | 0.00 |
| 1.80 | 2.46 | 0.01 | 0.01 | 1.58 | 2.09 | 0.00 | 4.04 |
| 1.80 | 2.46 | 1.30 | 3.08 | 0.00 | 0.01 | 0.00 | 2.46 |
| *0.90* | *1.23* | *0.65* | *0.78* | *1.20* | *1.05* | *1.20* | *2.43* |

Table S9: Gate output values for all Boolean assignments for circuit 0xDF (Func. opt.) and mean values in italic. Figure S6**B** presents the corresponding structure.

| A | B | C | NOR2_2 | NOT_1 | NOT_3 | NOT_4 | OR2_0 |
|---|---|---|---|---|---|---|---|
| 0.01 | 0.00 | 0.00 | 0.01 | 4.87 | 2.53 | 1.63 | 4.87 |
| 0.01 | 0.00 | 1.80 | 0.01 | 4.86 | 0.01 | 1.63 | 4.86 |
| 0.01 | 2.46 | 0.00 | 0.01 | 4.87 | 2.53 | 1.63 | 7.32 |
| 0.01 | 2.46 | 1.80 | 0.01 | 4.86 | 0.01 | 1.63 | 7.31 |
| 1.30 | 0.00 | 0.00 | 0.01 | 4.87 | 2.53 | 0.00 | 4.87 |
| 1.30 | 0.00 | 1.80 | 4.03 | 0.00 | 0.01 | 0.00 | 0.00 |
| 1.30 | 2.46 | 0.00 | 0.01 | 4.87 | 2.53 | 0.00 | 7.32 |
| 1.30 | 2.46 | 1.80 | 4.03 | 0.00 | 0.01 | 0.00 | 2.46 |
| *0.65* | *1.23* | *0.90* | *1.02* | *3.65* | *1.27* | *0.82* | *4.88* |

Table S10: Energy per gate for all Boolean assignments for circuit 0x20 (Energy opt.) and mean values in italic. Figure S6**C** presents the corresponding structure.

| A | B | C | NOR2_2 | NOR2_4 | NOT_0 | NOT_1 | NOT_3 | O |
|---|---|---|---|---|---|---|---|---|
| 0.00 | 0.00 | 0.00 | 21096.40 | 11173.56 | 76.02 | 18.02 | 74.12 | 15.70 |
| 0.00 | 0.00 | 0.00 | 8612.22 | 11173.56 | 76.02 | 10833.12 | 74.12 | 22.72 |
| 0.00 | 0.00 | 0.00 | 21299.99 | 24306.17 | 66.93 | 18.02 | 74.12 | 15.70 |
| 0.00 | 0.00 | 0.00 | 8815.80 | 24306.17 | 66.93 | 10833.12 | 74.12 | 22.01 |
| 0.00 | 0.00 | 0.00 | 12550.05 | 87.71 | 17937.86 | 18.02 | 8739.06 | 16.90 |
| 0.00 | 0.00 | 0.00 | 65.86 | 87.71 | 17937.86 | 10833.12 | 8739.06 | 12204.56 |
| 0.00 | 0.00 | 0.00 | 21190.53 | 13220.34 | 71.75 | 18.02 | 8739.06 | 15.70 |
| 0.00 | 0.00 | 0.00 | 8706.35 | 13220.34 | 71.75 | 10833.12 | 8739.06 | 22.38 |
| *0.00* | *0.00* | *0.00* | *12792.15* | *12196.94* | *4538.14* | *5425.57* | *4406.59* | *1541.96* |

Table S11: Energy per gate for all Boolean assignments for circuit 0x20 (Func. opt.) and mean values in italic. Figure S6**C** presents the corresponding structure.

| A | B | C | NOR2_2 | NOR2_4 | NOT_0 | NOT_1 | NOT_3 | O |
|---|---|---|---|---|---|---|---|---|
| 0.00 | 0.00 | 0.00 | 41960.48 | 11036.64 | 97.42 | 11.47 | 63.67 | 7.26 |
| 0.00 | 0.00 | 0.00 | 22253.22 | 11036.64 | 97.42 | 14076.13 | 63.67 | 7.31 |
| 0.00 | 0.00 | 0.00 | 41959.73 | 23171.35 | 79.74 | 11.47 | 63.67 | 7.26 |
| 0.00 | 0.00 | 0.00 | 22252.46 | 23171.35 | 79.74 | 14076.13 | 63.67 | 7.31 |
| 0.00 | 0.00 | 0.00 | 19784.58 | 50.88 | 23181.99 | 11.47 | 7506.98 | 7.35 |
| 0.00 | 0.00 | 0.00 | 77.32 | 50.88 | 23181.99 | 14076.13 | 7506.98 | 24842.10 |
| 0.00 | 0.00 | 0.00 | 41960.31 | 12185.96 | 92.39 | 11.47 | 7506.98 | 7.26 |
| 0.00 | 0.00 | 0.00 | 22253.05 | 12185.96 | 92.39 | 14076.13 | 7506.98 | 7.31 |
| *0.00* | *0.00* | *0.00* | *26562.64* | *11611.21* | *5862.89* | *7043.80* | *3785.32* | *3111.65* |

Table S12: Gate output values for all Boolean assignments for circuit 0x20 (Energy opt.) and mean values in italic. Figure S6**C** presents the corresponding structure.

| A | B | C | NOR2_2 | NOR2_4 | NOT_0 | NOT_1 | NOT_3 | O |
|---|---|---|---|---|---|---|---|---|
| 0.01 | 0.00 | 0.00 | 0.00 | 0.01 | 1.58 | 2.32 | 2.09 | 0.00 |
| 0.01 | 0.00 | 1.80 | 0.00 | 0.01 | 1.58 | 0.01 | 2.09 | 0.00 |
| 0.01 | 2.46 | 0.00 | 0.00 | 0.01 | 1.62 | 2.32 | 2.09 | 0.00 |
| 0.01 | 2.46 | 1.80 | 0.00 | 0.01 | 1.62 | 0.01 | 2.09 | 0.00 |
| 1.30 | 0.00 | 0.00 | 0.00 | 3.10 | 0.00 | 2.32 | 0.01 | 0.00 |
| 1.30 | 0.00 | 1.80 | 2.39 | 3.10 | 0.00 | 0.01 | 0.01 | 2.39 |
| 1.30 | 2.46 | 0.00 | 0.00 | 0.01 | 1.60 | 2.32 | 0.01 | 0.00 |
| 1.30 | 2.46 | 1.80 | 0.00 | 0.01 | 1.60 | 0.01 | 0.01 | 0.00 |
| *0.65* | *1.23* | *0.90* | *0.30* | *0.78* | *1.20* | *1.16* | *1.05* | *0.30* |

Table S13: Gate output values for all Boolean assignments for circuit 0x20 (Func. opt.) and mean values in italic. Figure S6**C** presents the corresponding structure.

| A | B | C | NOR2_2 | NOR2_4 | NOT_0 | NOT_1 | NOT_3 | O |
|---|---|---|---|---|---|---|---|---|
| 0.01 | 0.00 | 0.00 | 0.00 | 0.02 | 4.03 | 3.58 | 1.63 | 0.00 |
| 0.01 | 0.00 | 2.46 | 0.00 | 0.02 | 4.03 | 0.00 | 1.63 | 0.00 |
| 0.01 | 1.80 | 0.00 | 0.00 | 0.01 | 4.03 | 3.58 | 1.63 | 0.00 |
| 0.01 | 1.80 | 2.46 | 0.00 | 0.01 | 4.03 | 0.00 | 1.63 | 0.00 |
| 1.30 | 0.00 | 0.00 | 0.00 | 3.86 | 0.01 | 3.58 | 0.00 | 0.00 |
| 1.30 | 0.00 | 2.46 | 4.86 | 3.86 | 0.01 | 0.00 | 0.00 | 4.86 |
| 1.30 | 1.80 | 0.00 | 0.00 | 0.02 | 4.03 | 3.58 | 0.00 | 0.00 |
| 1.30 | 1.80 | 2.46 | 0.00 | 0.02 | 4.03 | 0.00 | 0.00 | 0.00 |
| *0.65* | *0.90* | *1.23* | *0.61* | *0.98* | *3.03* | *1.79* | *0.82* | *0.61* |

Table S14: Energy per gate for all Boolean assignments for circuit 0x81 (Energy opt.) and mean values in italic. Figure S6**D** presents the corresponding structure.

| A | B | C | NOR2_1 | NOR2_2 | NOR2_3 | NOR2_4 | NOR2_6 | NOT_0 | NOT_5 | O |
|---|---|---|---|---|---|---|---|---|---|---|
| 0.00 | 0.00 | 0.00 | 14539.62 | 12551.63 | 93.57 | 40.61 | 23572.42 | 74.12 | 12.01 | 13868.55 |
| 0.00 | 0.00 | 0.00 | 152.39 | 22291.39 | 12130.65 | 8504.09 | 23572.42 | 74.12 | 12.01 | 57.61 |
| 0.00 | 0.00 | 0.00 | 190.56 | 12551.63 | 8688.63 | 7891.38 | 12486.60 | 8739.06 | 12.01 | 68.74 |
| 0.00 | 0.00 | 0.00 | 145.62 | 22291.39 | 20725.70 | 8623.54 | 12486.60 | 8739.06 | 12.01 | 55.91 |
| 0.00 | 0.00 | 0.00 | 14548.54 | 60.11 | 93.57 | 15229.01 | 11206.30 | 74.12 | 14742.41 | 30.89 |
| 0.00 | 0.00 | 0.00 | 161.31 | 9799.88 | 12130.65 | 8356.92 | 11206.30 | 74.12 | 14742.41 | 59.89 |
| 0.00 | 0.00 | 0.00 | 17771.60 | 60.11 | 8688.63 | 15228.40 | 120.46 | 8739.06 | 14742.41 | 30.89 |
| 0.00 | 0.00 | 0.00 | 17726.66 | 9799.88 | 20725.70 | 43.33 | 120.46 | 8739.06 | 14742.41 | 13869.01 |
| *0.00* | *0.00* | *0.00* | *8154.54* | *11175.75* | *10409.64* | *7989.66* | *11846.44* | *4406.59* | *7377.21* | *3505.19* |

Table S15: Energy per gate for all Boolean assignments for circuit 0x81 (Func. opt.) and mean values in italic. Figure S6**D** presents the corresponding structure.

| A | B | C | NOR2_1 | NOR2_2 | NOR2_3 | NOR2_4 | NOR2_6 | NOT_0 | NOT_5 | O |
|---|---|---|---|---|---|---|---|---|---|---|
| 0.00 | 0.00 | 0.00 | 22720.66 | 13695.91 | 87.59 | 77.53 | 21530.74 | 63.67 | 16.22 | 24841.61 |
| 0.00 | 0.00 | 0.00 | 153.39 | 27760.57 | 16615.68 | 22251.32 | 21530.74 | 63.67 | 16.22 | 7.31 |
| 0.00 | 0.00 | 0.00 | 192.48 | 13695.91 | 8752.76 | 22248.23 | 12801.98 | 7506.98 | 16.22 | 7.31 |
| 0.00 | 0.00 | 0.00 | 154.29 | 27760.57 | 25280.48 | 22251.26 | 12801.98 | 7506.98 | 16.22 | 7.31 |
| 0.00 | 0.00 | 0.00 | 22742.76 | 33.50 | 87.59 | 19707.91 | 8773.73 | 63.67 | 9755.99 | 7.35 |
| 0.00 | 0.00 | 0.00 | 175.49 | 14098.16 | 16615.68 | 22250.39 | 8773.73 | 63.67 | 9755.99 | 7.31 |
| 0.00 | 0.00 | 0.00 | 19114.14 | 33.50 | 8752.76 | 19708.93 | 44.94 | 7506.98 | 9755.99 | 7.35 |
| 0.00 | 0.00 | 0.00 | 19075.95 | 14098.16 | 25280.48 | 78.42 | 44.94 | 7506.98 | 9755.99 | 24839.61 |
| *0.00* | *0.00* | *0.00* | *10541.15* | *13897.03* | *12684.13* | *16071.75* | *10787.85* | *3785.32* | *4886.11* | *6215.65* |

Table S16: Gate output values for all Boolean assignments for circuit 0x81 (Energy opt.) and mean values in italic. Figure S6**D** presents the corresponding structure.

| A | B | C | NOR2_1 | NOR2_2 | NOR2_3 | NOR2_4 | NOR2_6 | NOT_0 | NOT_5 | O |
|---|---|---|---|---|---|---|---|---|---|---|
| 0.00 | 0.01 | 0.00 | 0.00 | 0.00 | 2.50 | 2.71 | 0.01 | 2.09 | 2.32 | 2.71 |
| 0.00 | 0.01 | 1.80 | 1.33 | 0.00 | 0.01 | 0.01 | 0.01 | 2.09 | 2.32 | 0.01 |
| 0.00 | 1.30 | 0.00 | 1.24 | 0.00 | 0.02 | 0.01 | 0.01 | 0.01 | 2.32 | 0.01 |
| 0.00 | 1.30 | 1.80 | 1.35 | 0.00 | 0.01 | 0.01 | 0.01 | 0.01 | 2.32 | 0.01 |
| 2.46 | 0.01 | 0.00 | 0.00 | 2.39 | 2.50 | 0.01 | 0.01 | 2.09 | 0.01 | 0.01 |
| 2.46 | 0.01 | 1.80 | 1.31 | 0.00 | 0.01 | 0.01 | 0.01 | 2.09 | 0.01 | 0.01 |
| 2.46 | 1.30 | 0.00 | 0.00 | 2.39 | 0.02 | 0.01 | 3.05 | 0.01 | 0.01 | 0.01 |
| 2.46 | 1.30 | 1.80 | 0.00 | 0.00 | 0.01 | 2.71 | 3.05 | 0.01 | 0.01 | 2.71 |
| *1.23* | *0.65* | *0.90* | *0.65* | *0.60* | *0.64* | *0.69* | *0.77* | *1.05* | *1.16* | *0.69* |

Table S17: Gate output values for all Boolean assignments for circuit 0x81 (Func. opt.) and mean values in italic. Figure S6**D** presents the corresponding structure.

| A | B | C | NOR2_1 | NOR2_2 | NOR2_3 | NOR2_4 | NOR2_6 | NOT_0 | NOT_5 | O |
|---|---|---|---|---|---|---|---|---|---|---|
| 0.00 | 0.01 | 0.00 | 0.01 | 0.00 | 3.77 | 4.86 | 0.01 | 1.63 | 2.39 | 4.86 |
| 0.00 | 0.01 | 2.46 | 4.03 | 0.00 | 0.01 | 0.00 | 0.01 | 1.63 | 2.39 | 0.00 |
| 0.00 | 1.30 | 0.00 | 4.03 | 0.00 | 0.02 | 0.00 | 0.01 | 0.00 | 2.39 | 0.00 |
| 0.00 | 1.30 | 2.46 | 4.03 | 0.00 | 0.01 | 0.00 | 0.01 | 0.00 | 2.39 | 0.00 |
| 1.80 | 0.01 | 0.00 | 0.01 | 3.56 | 3.77 | 0.00 | 0.02 | 1.63 | 0.00 | 0.00 |
| 1.80 | 0.01 | 2.46 | 4.03 | 0.00 | 0.01 | 0.00 | 0.02 | 1.63 | 0.00 | 0.00 |
| 1.80 | 1.30 | 0.00 | 0.01 | 3.56 | 0.02 | 0.00 | 3.16 | 0.00 | 0.00 | 0.00 |
| 1.80 | 1.30 | 2.46 | 0.01 | 0.00 | 0.01 | 4.86 | 3.16 | 0.00 | 0.00 | 4.86 |
| *0.90* | *0.65* | *1.23* | *2.02* | *0.89* | *0.95* | *1.22* | *0.80* | *0.82* | *1.20* | *1.22* |

20. Ref. *15*, Supplement.

21. Ref. *15*, Figure S39.

22. Alberts, B., Johnson, A., Lewis, J., Raff, M., Roberts, K., and Walter, P. *Molecular Biology of the Cell*, 6th ed.; Garland Science: New York, NY, 2017.

23. Milo, R., and Phillips, R. *Cell Biology by the Numbers*; Garland Science, 2015.

24. Coulon, A., Ferguson, M. L., de Turris, V., Palangat, M., Chow, C. C., and Larson, D. R. (2014) Kinetic competition during the transcription cycle results in stochastic RNA processing. *eLife 3*, e03939.

25. Pelechano, V., Chavez, S., and Perez-Ortin, J. E. (2010) A Complete Set of Nascent Transcription Rates for Yeast Genes. *PLOS One 5*, e15442.

26. Lahtvee, P.-J., Sánchez, B. J., Smialowska, A., Kasvandik, S., Elsemman, I. E., Gatto, F., and Nielsen, J. (2017) Absolute Quantification of Protein and mRNA Abundances Demonstrate Variability in Gene-Specific Translation Efficiency in Yeast. *Cell Systems 4*, 495–504.

27. Wang, Y., Liu, C. L., Storey, J. D., Tibshirani, R. J., Herschlag, D., and Brown, P. O. (2002) Precision and functional specificity in mRNA decay. *Proceedings of the National Academy of Sciences 99*, 5860–5865.

28. Wagner, A. (2005) Energy Constraints on the Evolution of Gene Expression. *Molecular biology and evolution 22*, 1365–1374.

29. Hausser, J., Mayo, A., Keren, L., and Alon, U. (2019) Central dogma rates and the trade-off between precision and economy in gene expression. *Nature Communications 10*, 68.

30. Houseley, J., and Tollervey, D. (2009) The Many Pathways of RNA Degradation. *Cell 136*, 763–776.

31. Benaroudj, N., Zwickl, P., Seemüller, E., Baumeister, W., and Goldberg, A. L. (2003) ATP Hydrolysis by the Proteasome Regulatory Complex PAN Serves Multiple Functions in Protein Degradation. *Molecular Cell 11*, 69–78.
